# Ecological Drivers of Nontuberculous Mycobacteria in Aquatic Systems: Biodiversity, Niche Competition, and Pathogen Emergence

**DOI:** 10.64898/2026.02.02.701937

**Authors:** Marine Combe, Amar Bouam, Sylvestre Dizoe, Bernard Davoust, Emilien Drancourt, Dylan Messeca, Alice Valentini, Romain Blaizot, Rodolphe Elie Gozlan

**Affiliations:** ISEM, Université de Montpellier, CNRS, IRD, EPHE, Montpellier, France; Aix-Marseille Univ., IRD, MEPHI, IHU Méditerranée Infection, Marseille, France; Programme National de Lutte contre l’Ulcère de Buruli (PNLUB), Kongouanou Care Center, Yamoussoukro, Côte d’Ivoire; Aix-Marseille Univ., RITMES, AP-HM, SSA, IHU Méditerranée Infection, Marseille, France; SPYGEN, Savoie Technolac, BP 274 Le Bourget-du-Lac, France; Service de Dermatologie, Centre Hospitalier Andrée Rosemon, Cayenne, French Guiana

**Keywords:** Biogeography, Bacterial communities, Freshwater, Tropical, eDNA

## Abstract

Microbial diversity remains largely unexplored across environments and scales, notably because at local scales many microbial taxa exist under a dormant state. Microbial biogeography is shaped by edaphic and ecological drivers, and shifts in microbial community composition are frequently associated with host community structure and health. Nontuberculous mycobacteria represent a striking example of environmental microorganisms with opportunistic pathogenic potential. Unfortunately, data on their diversity, distribution, and ecological interactions in aquatic environments remain limited. However, understanding competition for niche space and the role of abiotic and biotic factors shaping their biogeography is crucial for predicting disease emergence and transmission. Here we aimed at i) identifying microhabitat abiotic and biotic drivers influencing their distribution, ii) assessing the predictability of their diversity and distribution across continents, and iii) examining potential exclusion or associations between pathogenic and nonpathogenic mycobacterial species. By deploying an eDNA-based metabarcoding approach from freshwater samples collected in urban and rural sites in French Guiana and Côte d’Ivoire, we have boosted our understanding of environmental mycobacteria ecology by highlighting the influence of habitat type, abiotic factors, and microbial interactions on mycobacterial distribution. In addition, the detection of pathogenic species further highlighted the importance of environmental reservoirs in mycobacterial disease transmission.

## Introduction

Anthropogenic pressures, including climate change, deforestation and intensive agriculture, and urban expansion are key drivers of biodiversity changes, and shifts in microbial community compositions, ultimately influencing the emergence of infectious diseases (Daszak et al., 2000; Morris et al., 2014; Wu et al., 2016; Morris et al., 2016; Cable et al., 2017; Stat et al., 2017; Reverter et al., 2020). In ecological theory, biodiversity is recognized as fundamental to ecosystem resilience (Stat et al., 2017; Jagadesh et al., 2019) and has been hypothesized to mitigate infectious diseases risks through the “dilution effect” although this remains debated (Keesing et al., 2006; Keesing & Ostfeld, 2021).

Tropical freshwater ecosystems illustrate these dynamics, where for example deforestation-driven aquatic food web alterations reduce top fish predators, favouring low-trophic taxa that harbor *Mycobacterium ulcerans* (*M. ulcerans*) the causative agent of Buruli ulcer, thereby increasing human exposure risk to the pathogen (Morris et al., 2016). These shifts align with the pathobiome concept in human and animal health, where microbiome disturbances promote pathogen proliferation (Bass et al., 2019; Pitlik & Koren, 2017). Pathogenic transitions from diverse, stable microbiomes to dysbiotic states occur under multiple abiotic and biotic stressors, including temperature, host immune responses, and microbiota composition (Lai et al., 2015). Thus, unraveling microbial diversity and interactions is not only fundamental to microbial biogeography and community assembly but also to understanding the relationships between biodiversity, ecosystem stability, and health within the One Health framework (Combe & Gozlan, 2024).

Microbial diversity remains largely unexplored across environments and scales (Curtis et al. 2002). The classical principle proposed by Baas Becking (1934), *“Everything is everywhere, but, the environment selects,”* suggests that microorganisms are globally distributed but often remain under a dormant state poorly amenable to routine culture, until specific abiotic or biotic conditions trigger their proliferation (de Wit & Bouvier, 2006). At local scales, dormancy may provoke that many microbial taxa exist below culture-based detection, reinforcing the need for complementary molecular approaches beyond traditional enrichment cultures. Advances in environmental DNA (eDNA)-based metabarcoding have significantly enhanced biosurveillance and biodiversity assessments across multiple habitats, including air, sediments, soil, and aquatic systems (Valentini et al. 2016; Drummond et al. 2015; Olds et al. 2016; Clare et al., 2021; Nuñez et al., 2021).

Microbial biogeography is shaped by edaphic and ecological drivers. For instance, soil bacterial diversity is primarily structured by pH, with higher diversity in neutral compared to acidic soils (Fierer and Jackson, 2006). Similarly, shifts in microbial community composition are frequently associated with host community structure and health (Bass et al. 2019). In ecosystems, trophic interactions influence pathogen dispersal; for example, *Lumbricus terrestris* earthworms contribute to *Mycobacterium bovis* spread in Brazil (Barbier et al., 2016), while transmission of *M. ulcerans* is likely mediated through food web interactions (Combe et al. 2017; Hammoudi et al., 2020). Understanding the distribution of potential pathogens and their interactions with macrofauna is therefore essential for assessing biodiversity-health relationships and predicting disease emergence risks.

Nontuberculous mycobacteria (NTM) represent a striking example of environmental microorganisms with opportunistic pathogenic potential. These bacteria are widely distributed across ecosystems (Nichols et al., 2004, Combe et al. 2023) and act as opportunistic pathogens affecting human and animal activities, including natural waters (fresh, brackish, and marine), drinking water systems, soils (Falkinham, 2002, 2009), aquaria, aquaculture reservoirs (Yanong & Pouder, 2010), and wastewater (Amha et al., 2017; Combe M, unpublished data). Their associated diseases pose significant health risks and economic burdens (Douine et al. 2017), particularly in aquaculture and fish farming in developing countries (Phung et al., 2013). Among NTM species infecting humans, the *Mycobacterium avium* complex (MAC) is most frequently implicated, causing lymphadenitis in children, respiratory infections in the elderly, and disseminated disease in immunocompromised individuals (Nichols et al., 2004). Other species, such as *Mycobacterium fortuitum*, *Mycobacterium abscessus*, *Mycobacterium chelonae,* and *Mycobacterium smegmatis* are associated with skin infections (De Groote and Johnson, 2004), while *Mycobacterium haemophilum* induces cutaneous and systemic infections majority in immunocompromised patients (Straus et al. 1994). In low- and middle-income countries, Mycolactone-Producing-Mycobacteria (MPM), including *M. ulcerans*, *Mycobacterium shinshuense*, *Mycobacterium liflandii*, and *Mycobacterium pseudoshottsii* are responsible for Buruli ulcer, a debilitating skin disease (Douine et al., 2017; Combe et al. 2017, 2023). Notably, *M. gilvum* has been identified from cutaneous ulcers clinically mimicking the ones of Buruli ulcers in patients in French Guiana, sometimes as co-infections with unidentified mycobacterial species (Combe et al. 2020).

Environmental factors shape NTM proliferation in aquatic habitats rich in organic matter or animal excreta, as well as in disturbed or polluted swamps characterized by high temperatures, low *pH*, low dissolved oxygen, and elevated zinc levels (Oloya et al., 2008; Jagadesh et al., 2019). Furthermore, NTM species exhibit distinct environmental preferences (e.g. brackish vs freshwater systems) and metabolic strategies (e.g. slow *vs* rapid growth) (Cangelosi et al., 2004), suggesting that local selective pressures constrain their biogeography. Despite their ubiquity, competition for niche space between NTM remains an understudied ecological process (Stubbendieck et al. 2016). Niche partitioning likely occurs through *(i)* resource exploitation, where one bacterium limits competitor growth by depleting shared resources, or *(ii)* interference, where antagonistic factors hinder competitors. Co-infections in animals and humans may arise from both antagonism and synergism between pathogens (Otta et al., 2014; Chen et al., 2017), leading to diverse outcomes. For example, interactions between *Aeromonas* strains enhanced virulence compared to single-strain infections (Mosser et al., 2015); co-infection with *Bolbophorus damnificus* and *Edwardsiella ictaluri* increased mortality rates in *Ictalurus punctatus* (Labrie et al., 2004); and in *Caenorhabditis elegans*, co-infection with *Acinetobacter baumannii* and *Candida albicans* suppressed *C. albicans* virulence (Peleg et al., 2008).

Among NTM, all MPM species isolated from humans, fish, and frogs produce mycolactone, an immunosuppressive toxin encoded on a virulence plasmid (Doig et al. 2012), which may enhance their competitive fitness in host tissues or environmental reservoirs. Additionally, cooperative interactions such as quorum sensing (QS), metabolite and gene product sharing (Black Queen Hypothesis, Loss, 2012), and horizontal gene transfer of virulence factors (Derome et al., 2016) may facilitate the emergence of opportunistic pathogens. The mechanisms may explain the occurrence of *M. chelonae*, *M. smegmatis* and *M. gilvum,* a non-MPM mycobacteria lacking the virulence plasmid, as co-infections in Buruli ulcer cases in Côte d’Ivoire and French Guiana (Nguetta et al., 2018; Combe et al., 2020, 2023).

Despite their known health and economic impacts, data on NTM diversity, distribution, and ecological interactions in aquatic environments remain limited. Understanding competition for niche space and the role of abiotic and biotic factors shaping their biogeography is crucial for predicting disease emergence and transmission. This study aims to: (1) identify microhabitat abiotic drivers (*pH*, oxygen, conductivity, temperature, phosphorus, nitrogen, carbon, C/N ratio, substrate) influencing NTM distribution; (2) evaluate the role of aquatic macro-biodiversity in shaping NTM patterns; (3) assess the predictability of NTM diversity and distribution across continents and (4) examine potential exclusion or associations between pathogenic and nonpathogenic mycobacterial species.

## Materials & Methods

### Study sites

In French Guiana (FG), 13 aquatic sites (ponds and oxbows) were sampled along the coast in mid-July 2017, at the onset of the dry season. These included 8 urban and 5 rural sites (Table 1; Fig. 1A; Supplementary Fig. S1; maps were generated using QGIS v3.36.2). Sampling in Côte d’Ivoire (CI) occurred in May 2018 (early rainy season) and included 12 urban or peri-urban water bodies near Bouaflé, Sinfra and Yamoussoukro (Fig. 1B; Supplementary Fig. S2). All sites were located in Buruli ulcer-endemic areas and were selected because clinically confirmed cases had been diagnosed there by physicians from the National Buruli Ulcer Control Program (PNLUB). Additionally, *M. ulcerans* had previously been detected at these sites, both in water bodies (Combe et al., 2019) and on plants (Hammoudi et al., 2020).

**Figure 1.**
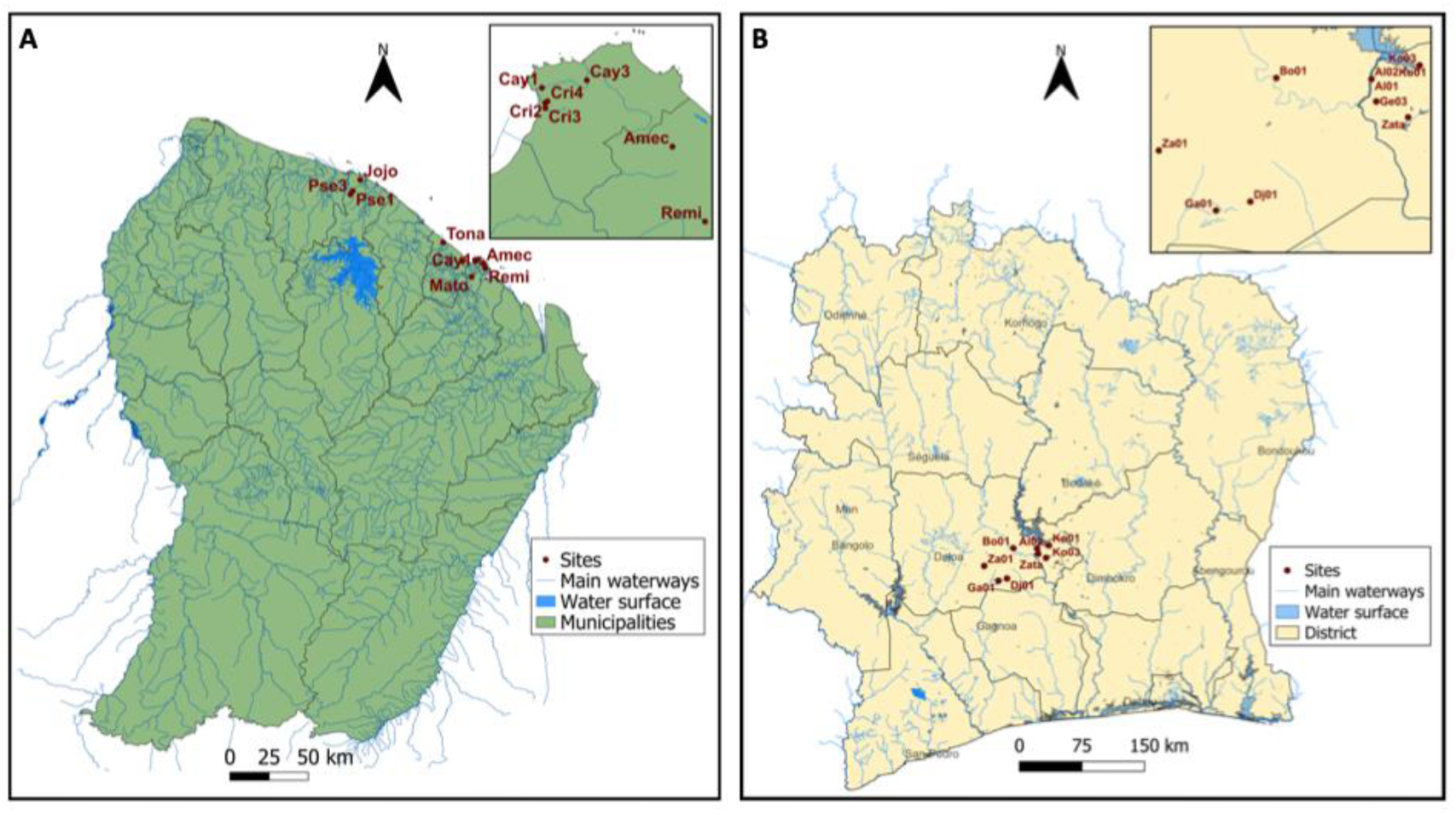
Localization of sampling sites in French Guiana (A) and Côte d’Ivoire (B). Sampling sites are indicated in red (see Table 1 for site abbreviations).

**Table 1.** Study sites in French Guiana (FG) and Côte d’Ivoire (CI). W: water samples; SS: surface sediment samples; SC: sediment core samples

| Country | Site name | Abbreviation | Latitude | Longitude | Area | Samples |
| --- | --- | --- | --- | --- | --- | --- |
| FG | Jojo | Jojo | 5,3941 | -52,992017 | Rural | W, SS |
| FG | Piste St Elie 1 | Pse1 | 5,32946 | -53,03592 | Rural | W, SS |
| FG | Piste St Elie 3 | Pse3 | 5,31043 | -53,04884 | Rural | W, SS |
| FG | Tonate | Tona | 5,03535 | -52,516483 | Rural | W, SS |
| FG | Matoury | Mato | 4,838083 | -52,35325 | Rural | W, SS |
| FG | Soula | Soul | 4,929067 | -52,403817 | Urban | W, SS |
| FG | Cayenne 1 | Cay1 | 4,93505 | -52,32441 | Urban | W, SS |
| FG | Cayenne 3 | Cay3 | 4,93786 | -52,31704 | Urban | W, SS, SC |
| FG | Crique 2 | Cri2 | 4,92908 | -52,33273 | Urban | W, SS, SC |
| FG | Crique 3 | Cri3 | 4,92697 | -52,33201 | Urban | W, SS |
| FG | Crique 4 | Cri4 | 4,93037 | -52,33123 | Urban | W, SS |
| FG | Ame Claire | Amec | 4,91411 | -52,2864 | Urban | W, SS, SC |
| FG | Remire | Remi | 4,88733 | -52,27473 | Urban | W, SS, SC |
| CI | Zagueta | Za01 | 6,783342 | -6,067334 | Urban | W, SS |
| CI | Bouaflé | Bo01 | 6,977789 | -5,752347 | Urban | W, SS |
| CI | Djenedougoufla | Dj01 | 6,6465 | -5,821809 | Urban | W, SS |
| CI | Gbalibonou | Ga01 | 6.622534 | -5.913820 | Urban | W, SS |
| CI | Zatta | Zata | 6,872015 | -5,398372 | Urban | W, SS |
| CI | Geoffroy, Toumobokro | Ge02 | 6,914796 | -5,484185 | Urban | W, SS |
| CI | Geoffroy, Toumobokro | Ge03 | 6,914796 | -5,484185 | Urban | W, SS |
| CI | Allaiyaokro/Allanykro | Al01 | 6,974977 | -5,497005 | Urban | W, SS |
| CI | Allaiyaokro/Allanykro | Al02 | 6,974977 | -5,497005 | Urban | W, SS |
| CI | Kongouanou | Ko03 | 7,011667 | -5,367778 | Urban | W, SS |
| CI | Kongouanou | Ko01 | 7,011667 | -5,367778 | Urban | W, SS |
| CI | NA | Vo02 | NA | NA | Urban | W, SS |

NA : unknown information

### Environmental sampling

In French Guiana, sampling was carried out under the French Nagoya Protocol (#166344). In Côte d’Ivoire, sampling was carried out after submitting a request for authorization under the Nagoya Protocol to the Ministry, and in consultation with local collaborators from the National Buruli Ulcer Control Program (PNLUB). At each site, 20×100 mL water samples were collected from different micro-habitats (total of 2 L) using a sterile ladle and transferred to 2 L Whirl-Pak® bags. Water was immediately filtered through 0.45 µm VigiDNA^®^ filter capsules (SPYGEN, France) using a sterile syringe until depletion or clogging. Capsules were filled with 80 mL CL1 conservation buffer (SPYGEN, France) and stored at room temperature before DNA extraction. Physicochemical parameters (temperature, oxygen, pH, conductivity) were recorded (Supplementary Table S1). Surface sediments (0-1 cm) were sampled at all site using a sterile Falcon tube and stored at 4°C for 24 h before DNA extraction. Additionally, sediment cores from four urban sites in FG were collected at depths of 0-1 cm; 1-2 cm; 2-3 cm; 3-4 cm; 4-5 cm; 5-10 cm and subsampled at each sediment layer for DNA extractions and physicochemical analysis (P, N, C, C/N ratio, granulometry) (Supplementary Table S1).

### Aquatic invertebrate sampling

Invertebrates were collected using a 25 cm x 45 cm dip net (500 µm mesh) from five areas per site, sampling across microhabitats. The net was moved from the sediment surface upward over a 1m² area. Specimens were sorted and preserved in 70% alcohol for taxonomic identification at the family level (Supplementary Table S2).

### DNA extractions

For water, the DNA extraction was performed following the protocol described in (Pont et al., 2018). each filtration capsule was agitated for 15 min on S50 Shaker (Cat Ingenieurbüro™) at 800 rpm and the CL1 buffer inside the capsule was emptied into a 50 mL tube that was then centrifuged for 15 min at 15,000 × g. The supernatant was removed with a sterile pipette by leaving 15 mL of liquid at the bottom of the tube. Subsequently, 33 mL of ethanol and 1.5 mL of 3M sodium acetate were added to each 50 mL tube that were stored overnight at -20°C. The tubes were centrifuged at 15,000 × g for 15 min at 6°C and the supernatant was discarded. Then, 720 µL of ATL Buffer from the DNeasy Blood & Tissue Extraction Kit (QIAGEN GmbH, Hilden, Germany) were added, the tubes were vortexed and the supernatants were transferred into 2 mL tubes previously filled with 20 µL of Proteinase K. Tubes were incubated at 56°C for two hours. Subsequently, DNA extractions were performed using the NucleoSpin® Soil kit (MACHEREY-NAGEL GmbH & Co., Düren, Germany) starting from step 6 and following the manufacturer’s instructions. Elution was performed by adding 100 µL of SE buffer twice. Negative extraction controls were included in each batch of extraction to monitor any possible contamination. Total DNA was also extracted from 250 mg of sediments using the PowerSoil DNA extraction kit (QIAGEN) and following the manufacturer’s recommendations, including a negative control as well. Extracted DNA was stored at -20°C.

### Detection of mycolactone-producing-mycobacteria (MPM) by qPCR

To detect and quantify the relative abundance of MPM DNA in sediment samples we performed two TaqMan qPCR runs targeting the insertion sequence *IS*2404 and the ketoreductase B (KR) domain of the mycolactone polyketide synthase gene (Combe et al., 2023). To amplify *IS*2404 genetic marker, we used the following primer and probes: *IS*2404 forward primer 5’-ATTGGTGCCGATCGAGTTG-3’, *IS*2404 reverse primer 5’-TCGCTTTGGCGCGTAAA-3’ and *IS*2404 probe FAM-CACCACGCAGCATTCTTGCCGT-BHQ1. For KR amplification we used KR forward primer 5’-TCACGGCCTGCGATATCA-3’, KR reverse primer 5’-TTGTGTGGGCACTGAATTGAC-3’, and KR probe FAM-ACCCCGAAGCACTG-MGBNFQ. The qPCR reaction consisted of 1X TaqMan Gene Expression Master Mix (LifeTechnologies), 0.3 μM (final concentration) of each primer, 0.1 μM (final concentration) of the probe, 5 μl of DNA and water adjusted to a final volume per reaction of 25 μl. For KR we followed the same protocol except that we used the probe at a final concentration of 0.25 μM. An internal positive control (IPC) was added in each *IS*2404 reaction in order to test for the presence of PCR inhibitors in the environmental samples. As positive controls we used the *M. ulcerans* DNA at a concentration of 10^5^ bacteria/mL and negative (DNA replaced by sterile water) controls were included in each run. *M. ulcerans* positive control consisted of genomic DNA purified from a cultured strain from French Guiana (strain 1G897) and provided by Dr. Laurent Marsollier (ATOMYCA, Université d’Angers, France). This positive control was also used to run standard curves based on serial dilutions of purified DNA from 10^5^ to 10^0^ bacteria/mL (in triplicates). Standard curves allowed us to determine a threshold value to consider positive results (CT-values < 38). The assays were run in an Applied Biosystems 7300 Real Time PCR system, with the following program: one cycle at 50°C for 2 min, one cycle at 95°C for 10 min, followed by 45 cycles at 95°C for 15 sec and at 60°C for 1 min. Only samples with cycle threshold values < 38 for both *IS*2404 and KR markers were considered as positives (Combe et al., 2019). In all assays the negative controls remained negative.

### Primer design and in silico validation

Group-specific primers (22 in total) were designed using the *ecoprimers* software (Riaz et al., 2011) on a collection of 16S ribosomal DNA (rDNA) sequences, the most commonly used indicator gene for bacterial biodiversity assessment (Horner-Devine et al. 2004), of the order Corynebacteriales (including all sequences present on GenBank on the 25^th^ of August 2017). We selected only primer pairs with a difference of Tm < 4°C, a GC percentage comprises between 40% and 80%, indicated as GG in the output file, with a fragment length < 200 bp in order to capture degraded DNA, a Barcoding coverage (*Bc,* Ficetola et al. 2010)and Barcoding specificity (*Bs,* Ficetola et al. 2010) > 0.95 and with < 5 repetitions of the same base. Primers retained were aligned to *M. tuberculosis* 16S and their positions on the sequence were used to determinate the groups of primers that bind a similar genomic region using the function *Mclust* in the R package Mclust. Then, one primer pair per group (22 groups) were randomly choosen and tested *in silico* using the ecoPCR program (Bellemain et al., 2010; Ficetola et al., 2010) against the entire set of DNA sequences available from the EMBL-European Nucleotide Archive (release 132, standard sequences) and with a maximum of three mismatches in the entire primer sequence. For each primer, a sequence logo was generated (Crooks et al., 2004) based on *in silico* PCR results for the target and non-target group. A mismatch analysis was performed, both for the target and for the non-target taxonomic group to assess specificity. Five other primer pairs available in the literature and previously used for the detection of *Mycobacterium sp.* (Kox et al., 1995; Tobler et al., 2006; Phung et al., 2013; Pontiroli et al., 2013; Radomski et al., 2013) were tested using the same protocol. Finally, one primer pair (MycoF: 5’-CCACACCGCAAAAGCTTT-3’; MycoR: 5’-CGTGCTTAACACATGCAA-3’) was selected from the newly designed ones since it was the most conserved among the *Mycobacterium* group, it amplified a low number of non-target groups, and it amplified a fragment showing the smallest length polymorphism among the 27 primer pairs tested.

### DNA amplification and sequencing

DNA amplifications were performed in a final volume of 25 μL, using 3 μL of DNA extract as template. The amplification mixture consisted of 1 U of AmpliTaq Gold DNA Polymerase (Applied Biosystems, Foster City, CA), 10 mM of Tris-HCl, 50 mM of KCl, 2.5 mM of MgCl2, 0.2 mM of each dNTP, 0.2 μM of each primer, and 0.2 µg/µL of bovine serum albumin (BSA, Roche Diagnostic, Basel, Switzerland). Primers were 5’-labeled with an eight-nucleotide unique tag (with at least three differences between any pair of tags) allowing the assignment of each sequence to the corresponding sample during sequence analysis. Tags for forward and reverse primers were identical for each sample. The PCR mixture was denatured at 95°C for 10 min, followed by 40 cycles of 30 sec at 95°C, 30 sec at 55°C and 1 min at 72°C, and followed by a final elongation at 72°C for 7 min. The PCR was performed in a room dedicated to amplify DNA, with negative air pressure and physically separated from the DNA extraction rooms (with positive air pressure). Four PCR replicates were run per sample. One negative PCR (ultrapure water, with 12 replicates), two positive controls composed of pooled DNA extracts of seven *Mycobacterium* species DNA (*M. ulcerans*, *M. marinum*, *M. fortuitum*, *M. avium*, *M. chelonae*, *M. absessus*, *M. smegmatis*; *M. leprae; M. lepromatosis;* Supplementary Table S3) provided by Dr. Jean-Christophe Avarre (UMR ISEM, Montpellier, France) and three negative controls (DNA replaced by sterile water) for extractions were analyzed in parallel to monitor any possible contaminations and the sensitivity of the primer pairs.

Amplified DNAs were quantified using capillary electrophoresis (QIAxcel; QIAGEN) and purified using the MinElute PCR purification kit (QIAGEN). Purified PCR products were then pooled in equal volumes to achieve an expected theoretical sequencing depth of 700,000 reads per sample. Library preparation and sequencing were performed at Fasteris (Geneva, Switzerland). Libraries were prepared following the MetaFast protocol (Fateris, https://www.fasteris.com/dna/?q=content/metafast-protocol-amplicon-metagenomic-analysis) and a paired-end sequencing (2 x 125 bp) was carried out using an Illumina MiSeq (Illumina, San Diego, CA, USA) with the MiSeq Kit v3 (Illumina, San Diego, CA, USA) following the manufacturer’s instructions.

### Bioinformatics

Sequence reads were analyzed using the programs implemented in the OBITools package (http://metabarcoding.org/obitools; Boyer et al., 2016). Forward and reverse reads were assembled using illuminapairedend program and assigned to each sample using the ngsfilter program. Sequences shorter than 20 bp, occurring < 10 times per sample or labelled as “internal” by the obiclean program (corresponding most likely to PCR/sequencing errors) were discarded. The taxonomic assignment of MOTUs (Molecular Operational Taxonomic Units) was performed using the program ecotag, with a reference database built by using the 7753 complete genomes of *Mycobacterium sp.* (available on GenBank on the 29^th^ of September 2017, Supplementary Table S4), and resulting in a total of 42 MOTUs. MOTUs showing < 97% similarity with our reference database were removed since this 97% threshold is commonly used to define bacterial species (Stackebrandt & Goebel, 1994). Finally, all sequences with a frequency of occurrence below 0.002 per taxon were discarded.

### Analysis of mycobacteria communities’ diversity and spatial variability

Statistics were performed in RStudio (v2023.12.1). NA values in sequencing data, indicating absent MOTUs, were converted to zero. Genetic sequences and invertebrate taxonomic data were log10(x+1) transformed to improve visualization and meet statistical assumptions. Group differences were assessed via Student’s t test (normal data) or Wilcoxon Mann Whitney test (non-normal data) with a 5% alpha threshold. Post-hoc corrections (Tukey or the Dunn’s test with a Benjamini-Hochberg adjustment) were applied when necessary, with a Kruskall Walis test.

#### Alpha Diversity Analysis

(vegan package version 2.6.4) - Alpha diversity was assessed using the Shannon index to quantify the diversity of MOTU communities at each site. This index accounts for both species richness and evenness, providing a comprehensive measure of community structure. The analysis helps determine whether some sites host more diverse or more evenly distributed MOTU assemblages compared to others. Beta Dissimilarity Analysis (vegan package version 2.6.4) - To compare the composition of MOTU communities between sites, beta dissimilarity was evaluated using the Jaccard index. This measure captures differences in MOTUs presence or absence, allowing the identification of compositional variations across sampling locations. Such an approach is crucial for understanding spatial heterogeneity in microbial communities.

#### MOTU distribution in sediment cores

To characterize the vertical distribution of NTM MOTUs within sediment cores, an ANOVA was performed on MOTU richness (i.e., the number of distinct MOTUs per site/layer). This statistical test determines whether significant differences exist in MOTU diversity across sediment layers, helping to assess stratification patterns and potential environmental influences on microbial distribution.

#### Multidimensional Scaling Analysis

(vegan package version 2.6.4) - Non-Metric Multidimensional Scales (NMDS) was employed to visualize site relationships based on similarities and dissimilarities in MOTU composition. This technique reduces data dimensionality while preserving relative distances, making it particularly effective for representing ecological datasets and identifying clustering trends in microbial communities.

#### Hierarchical Clustering Analysis (HCA) and Principal Component Analysis (PCA) (hclust function and prcomp function of the stats package version 4.3.3, respectively)

HCA was used to classify sites based on their MOTU composition, distinguishing clusters of similar and dissimilar sites. The optimal dendrogram cut was determined using the inertia loss scree method. Additionally, PCA was performed on transformed raw data (centered but not scaled) to further reduce dataset complexity while maintaining variance structure. This approach aids in identifying dominant gradients influencing community composition.

#### Linear Discriminant Analysis (LDA) and Random Forest (RF) (caret package version 6.0.94)

LDA was applied following the PCA to formally define site grouping and identify key discriminant variables. By maximizing separation between predefined groups, LDA seeks highlights the most influential features distinguishing microbial communities. A 3-fold, 200 repetition cross-validation ensured model robustness. To further compare patterns between FG and CI, a RF analysis was conducted leveraging its ability to produce accurate predictions and assess variable importance.

#### MOTU Interaction Analysis (qgraph package version 1.9.8.)

Network analysis was performed to investigate potential correlation patterns, i.e. associations or exclusions, among MOTUs, for each type of habitat (rural, urban, water, surface sediment, core sediment). This method helps uncover co-occurrence patterns, which can reveal ecological interactions such as mutualistic or competitive relationships among microbial taxa. A Spearman’s correlation between each MOTU, commonly used for non-linear datasets, was calculated and correlations > 0 identified from pairwise comparisons were considered to further run and visualize a partial correlation network by using the function “pcor” in the qgraph package in R (Epskamp et al., 2012), where each node represents a MOTU and each edge stands for a correlation between the nodes, assuming that the bigger the edges (lines), the stronger the link between two MOTU.

#### Mycobacteria-Invertebrate Relationships (coinertia function of the ade4 package version 1.7.22)

To explore the relationship between mycobacteria MOTUs and invertebrate communities, a co-inertia analysis was conducted. This technique identifies covariation patterns between two datasets, providing insights into potential ecological interactions. It is particularly useful for understanding how microbial diversity correlates with macro-organism distributions, shedding light on functional linkages in the studied environment.

## Results

Illumina MiSeq paired-end sequencing yielded 73,881,081 reads, with 21,497,222 reads (298,573 per sample on average) assigned to 39 mycobacterial MOTUs at 97% similarity threshold, including an unidentified *Mycobacterium sp.* (Myco) MOTU. Whilst we focused only on NTM, Myco_group4 (Gp04) also encompassed *M. tuberculosis* due to its shared 16S region sequence targeted here with other MOTUs (Table 2). In FG, sequencing from water and sediments produced 36,800,559 reads, with 12,245,891 (510,246 per sample on average) assigned to MOTUs (Supplementary Fig. S3A), while sediment cores generated 19,919,577 reads, with 5,767,468 (240,311 per sample on average) assigned (Supplementary Fig. S3B). In CI, 17,160,945 reads were obtained, with 3,483,863 (145,161 per sample on average) assigned to MOTUs (Supplementary Fig. S3C). Also, a *M. lepromatosis* MOTU, a causative agent of leprosy, was included to assess its presence in environmental samples, but was not detected. A total of 27 MOTUs were found in both countries while some were only found country-specific. In FG, unique MOTUS (N=11) included *M. phlei* (Phle), *M. liflandii* (Lifl), *M. immunogenum* (Immu), *M. goodi* (Good), *M. avium* (Aviu), *M. avium* complex (Gp13), Myco-Group8 (Gp08 including *M. marinum*/*M. ulcerans)*, Myco-Group7 (Gp07 including *M. sinense*/*M. terrae*), Myco_Group4 (Gp04 including *M. africanum*/*M. bovis/M. canettii/ M. caprae/M. tuberculosis*), Myco_Group3 (Gp03 including *M. absessus*/*M. chelonae/M. saopaulense/M. sp. QIA-37/M. stephanolepidis*) and Myco_Goup10 (Gp10 including *M. chimaera/M. indicus pranii/M. intracellulare/M. sp. MOTT36Y/M. yongonense*), whereas *M. aurum* (Auru) was exclusive to CI (Table 2; Supplementary Fig. S4).

**Table 2:**
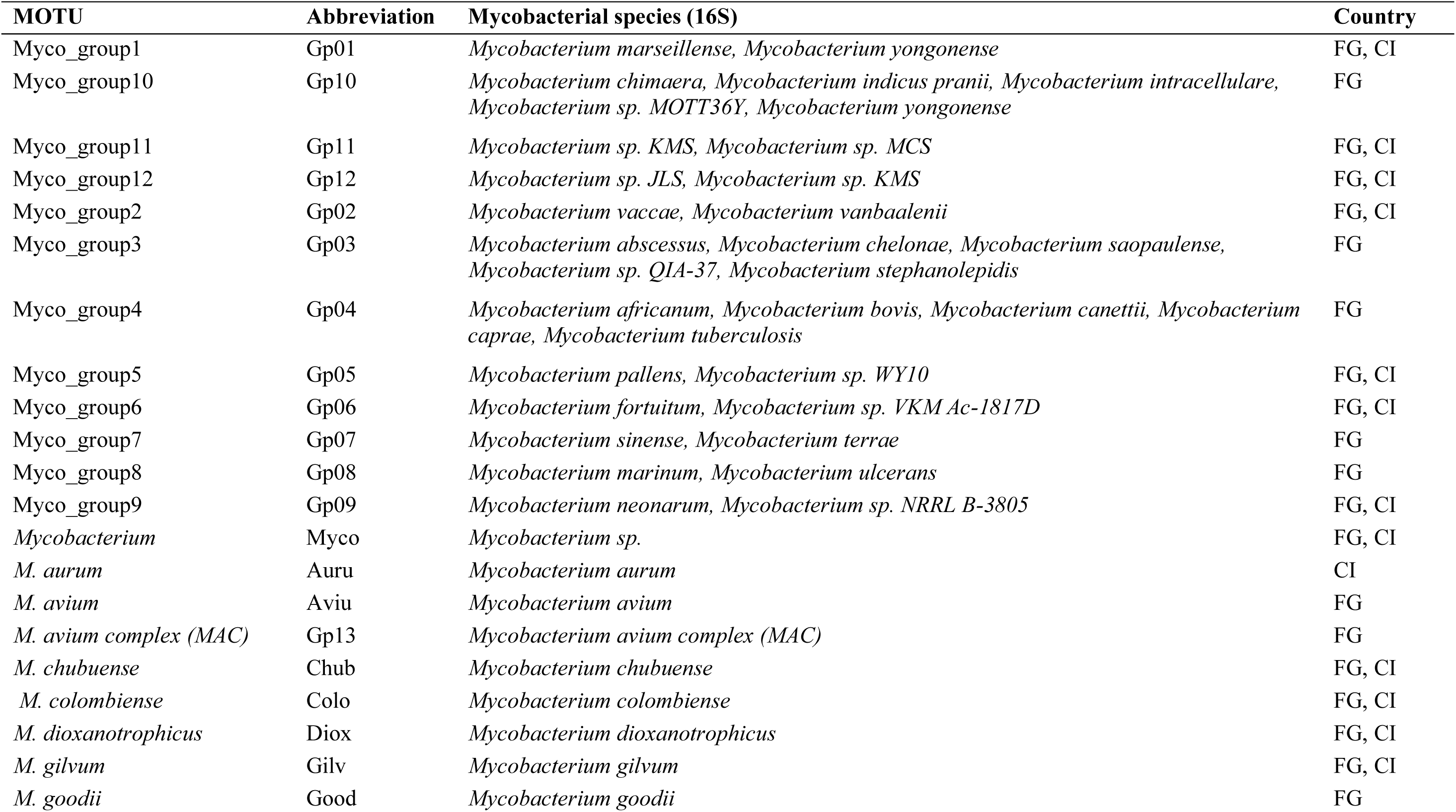
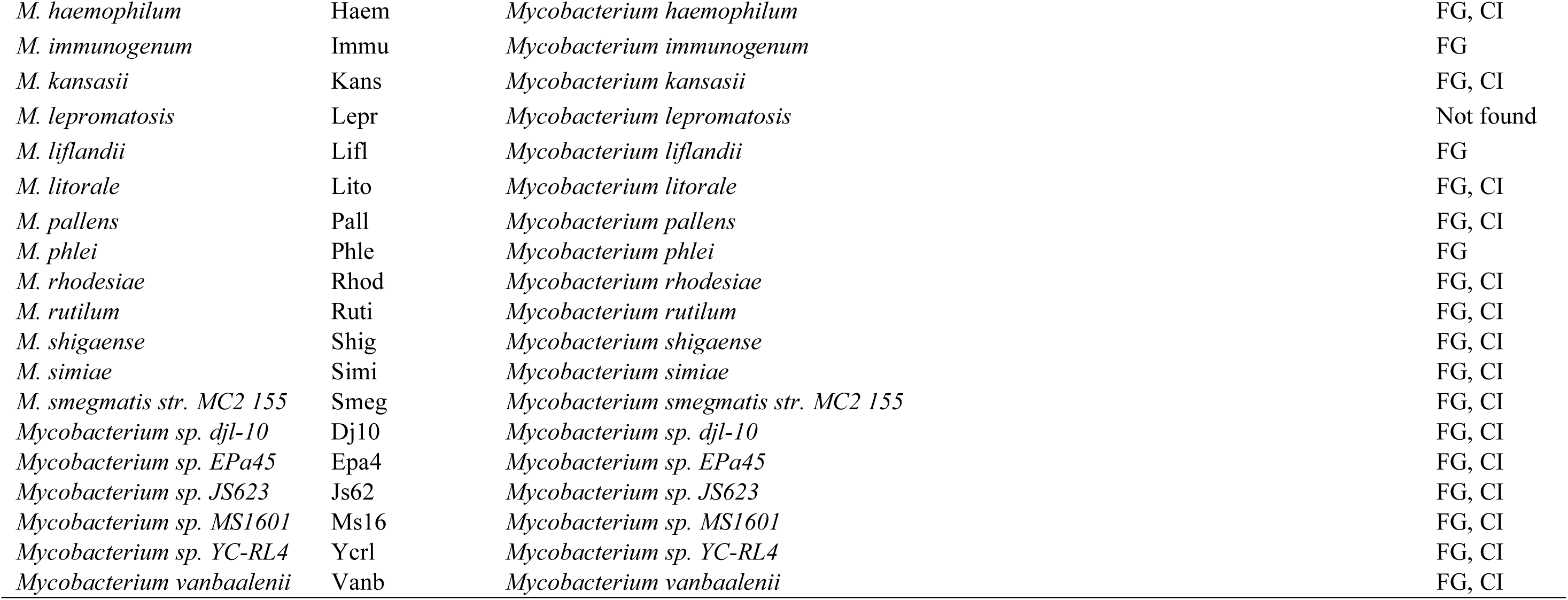
Dataset of the mycobacterial MOTUs identified by eDNA analyses of water and sediments (surface and core) in French Guiana (FG) and Côte d’Ivoire (CI). The name and the abbreviation of each MOTU are indicated, as well as the different mycobacterial species belonging to each MOTU based on their 16S genetic sequence.

### Diversity and distribution of mycobacterial MOTUs in FG freshwater habitats

#### Urban vs. Rural Sites

MOTU richness (alpha diversity) varied across sites, with the lowest at Tonate (Tona 0.53) and the highest at Jojo (Jojo 2,67), indicating differences in MOTU diversity and distribution balance (Table 3). Hierarchical Cluster Analysis (HCA) grouped sites into urban and rural clusters with further subgroups based on sample type (water vs. sediment; Fig. 2A). However, some sites deviated from this pattern, such as Jojo water (Jojo_W) clustering with sediment samples, or Piste St Elie 3 sediment (Pse3_S) clustering with water samples. Similarly, urban sites Remire sediment (Remi_S) and Cayenne 1 sediments (Cay1_S) clustered with urban water samples.

**Figure 2.**
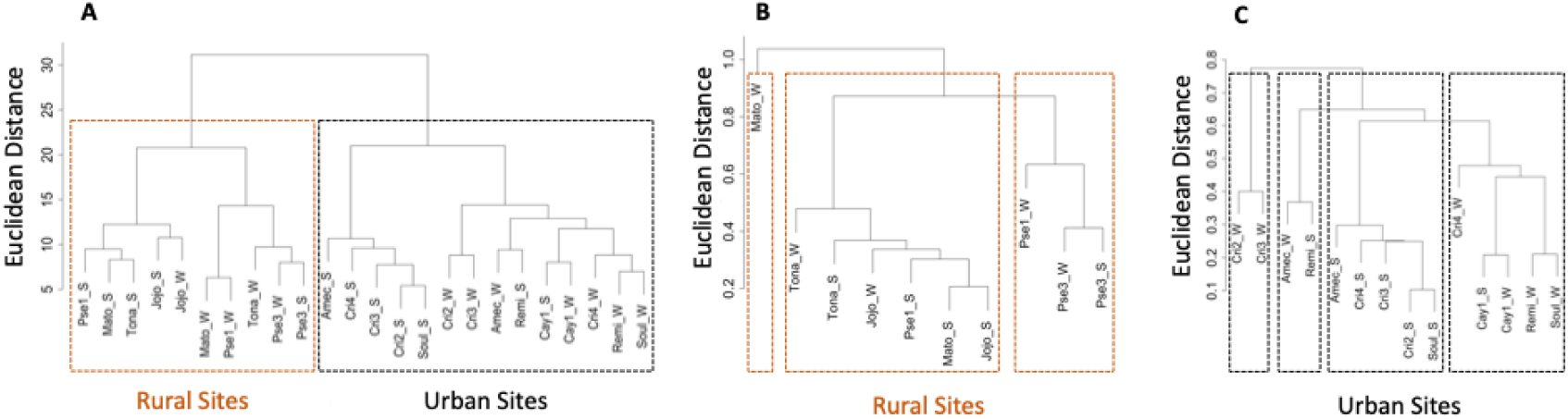
**A:** Hierarchical Cluster Analysis (HCA) of log-transformed data (reads) collected in French Guiana (FG). **B & C:** HCA of the dissimilarity matrix (Jaccard index). We used the Euclidean distance and the wardD2 jump. Site names are indicated (see Table 1 for site abbreviations) as well as the type of sample collected such as water (W) or surface sediment (S). **B**: rural sites; **C:** urban sites.

**Table 3.**
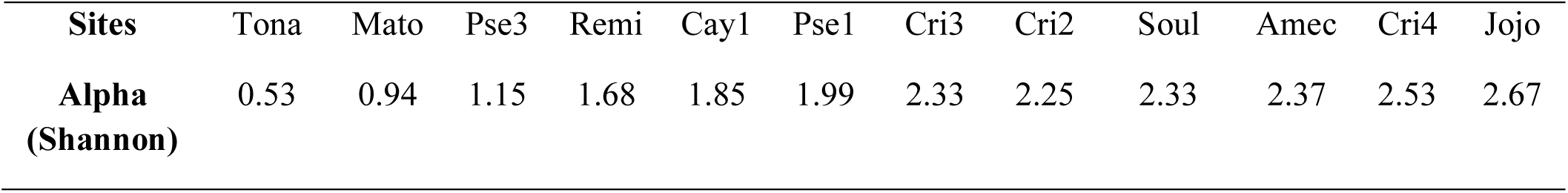
Alpha (Shannon) diversity on raw sequence data and for each site studied in French Guiana (FG). Data for both water and surface sediment were combined in order to obtain the global diversity of the site.

| **Sites** | Tona | Mato | Pse3 | Remi | Cay1 | Pse1 | Cri3 | Cri2 | Soul | Amec | Cri4 | Jojo |
| --- | --- | --- | --- | --- | --- | --- | --- | --- | --- | --- | --- | --- |
| **Alpha**<br>**(Shannon)** | 0.53 | 0.94 | 1.15 | 1.68 | 1.85 | 1.99 | 2.33 | 2.25 | 2.33 | 2.37 | 2.53 | 2.67 |

Principal Component Analysis (PCA) revealed distinct MOTU compositions between urban and rural sites (Fig. 3A). *M. shigaense* (Shig), *M. haemophilum* (Haem), and Myco_goup4 (Gp04) were associated with rural sites, while Myco_group6 (Gp06), *M. vanbaaleni* (Vanb) and *M. avium complex* (Gp13) were predominantly found in urban environments (Fig. 3B). Additionally, urban sites showed a segregation between water and surface sediment samples (Fig. 3A). Linear Discriminant Analysis (LDA) achieved 89% classification accuracy, identifying *M. shigaense* (Shig), Myco_group 6 (Gp06), and *M. haemophilum* (Haem) as key discriminants between urban and rural sites (Supplementary Fig. S5). The 100% site specificity of *M. shigaense* (rural) and Myco_group6 (urban) suggest their distribution is strongly linked to site micro-characteristics.

**Figure 3.**
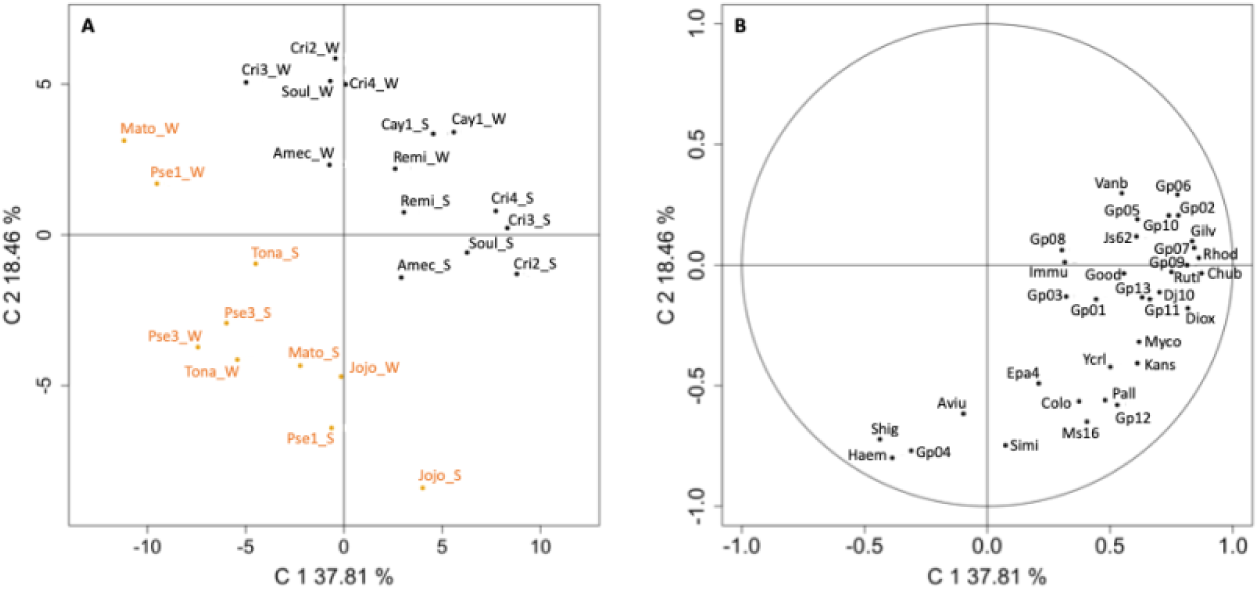
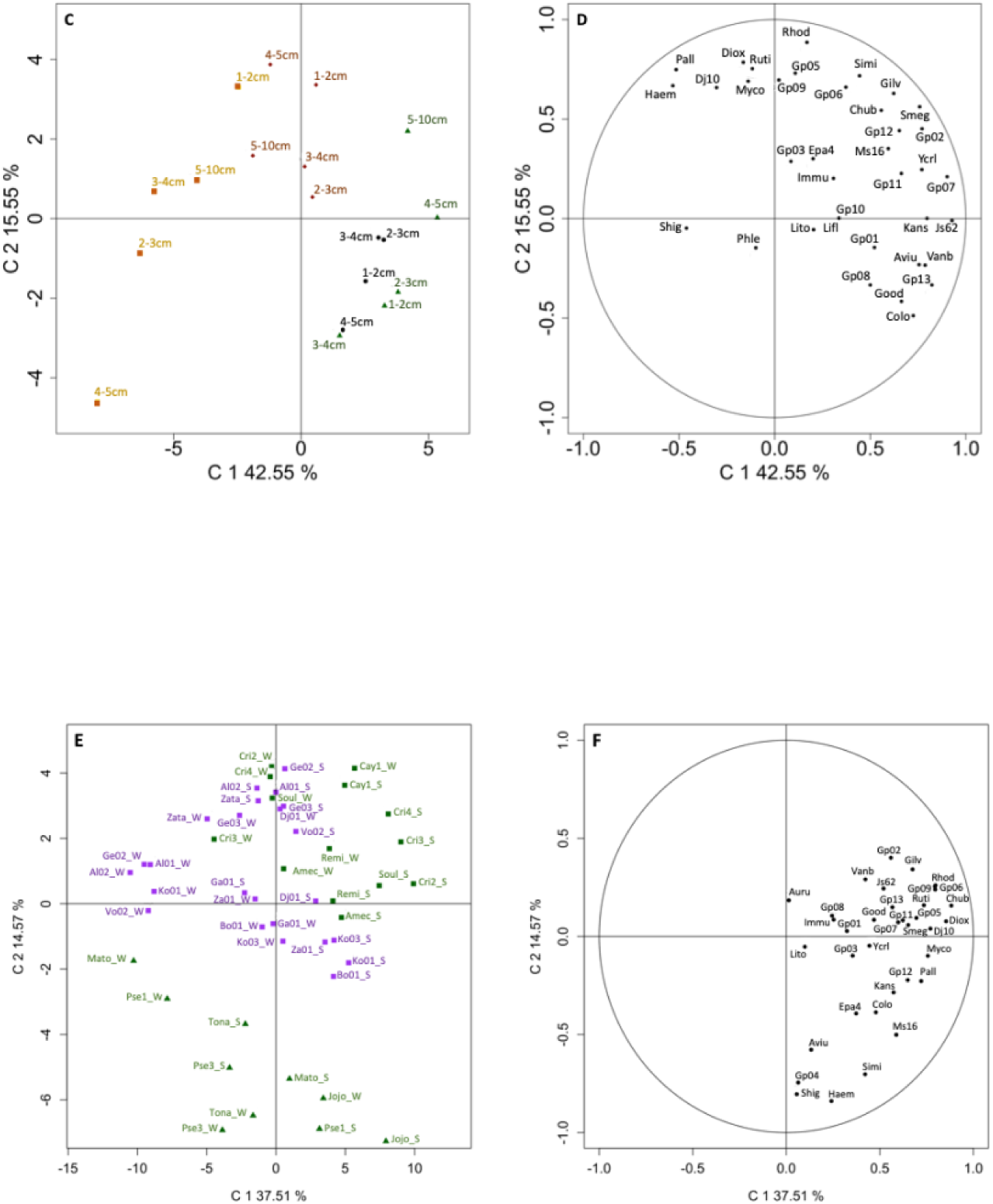
**A & B:** Principal Component Analysis (PCA) on log-transformed read data. Axis 1 explains 37.81% of the variability of the dataset while axis 2 explains 18.46% of this variability. **A**: cloud of the sites. Rural sites are indicated in orange while urban sites are indicated in black; **B**: circle of correlations between MOTUs and axes. The name of each site and each MOTU is indicated (see Table 1, 2). S: surface sediment samples; W: water samples. **C & D:** Unreduced centered PCA on log10(x+1) transformed core data. The Cay3_5-10 cm value was excluded as it was not representative. All sediment layers for the 4 sites (1-2 cm, 2-3 cm, 3-4 cm, 4-5 cm, 5-10 cm) are represented. **C:** cloud of individuals (sites); Orange square = Amec, Brown diamond = Remi, Green triangle = Cri2, Black circle = Cay3. **D:** circle of correlations between MOTUs and components. Axis C1 explains 42.55% of the variability of the dataset and axis C2 15.55%. **E & F:** Unreduced centered PCA on log10(x+1) transformed French Guiana (FG) and Côte d’Ivoire (CI) MOTU data. C1 explains 37.51% of the variability in the dataset and C2 14.57%. **E**: cloud of individuals (sites); **F**: circle of correlations between MOTUs and components. The sites were colored according to the country (FG in green and CI in purple), the triangles represent sites in rural areas and the squares in urban areas.

Given the urban vs. rural clustering, we further analyzed MOTU composition separately for each. Using beta dissimilarity (Jaccard index) HCA identified three rural site groups (Fig. 2B). Sites Piste St Elie 1 (Pse1) and Piste St Elie 3 (Pse3) clustered due to their upstream-downstream proximity, but Piste St Elie 1 sediment (Pse1_S) diverged, suggesting mycobacterial diversity is influenced more by microhabitat conditions than watershed dynamics. Similarly, Matoury water (Mato_W) clustered separately, indicating local abiotic/biotic factors shape mycobacterial distribution. For urban sites, HCA identified four distinct clusters (Fig. 2C), with Crique 4 water (Cri4_W) forming a unique group, suggesting a distinct MOTU composition compared to other urban sites in the same cluster.

#### Water vs Surface Sediment Samples

Alpha diversity (Shannon index) did not significantly differ between urban (U) and rural (R) sites or between water (W) and sediment (S) samples (ANOVA, *p* = 0.126; Fig. 4A) nor when comparing only water vs. sediment (Wilcoxon Mann–Whitney, *p* = 0.101; Fig. 4B). However, median alpha diversity tended to be higher in surface sediments (R_S, U_S) than in water samples (R_W, U_W; Fig. 4A). In contrast, the number of MOTUs was significantly higher in surface sediments than in water across all comparisons (R_S vs. R_W, U_S vs. U_W, and U_S vs. R_W; ANOVA, p = 0.002; Fig. 4C), with a marked difference between sediment and water samples (t-test, p < 0.001; Fig. 4D). These findings indicate greater mycobacterial MOTU diversity in surface sediments compared to water samples.

**Figure 4.**
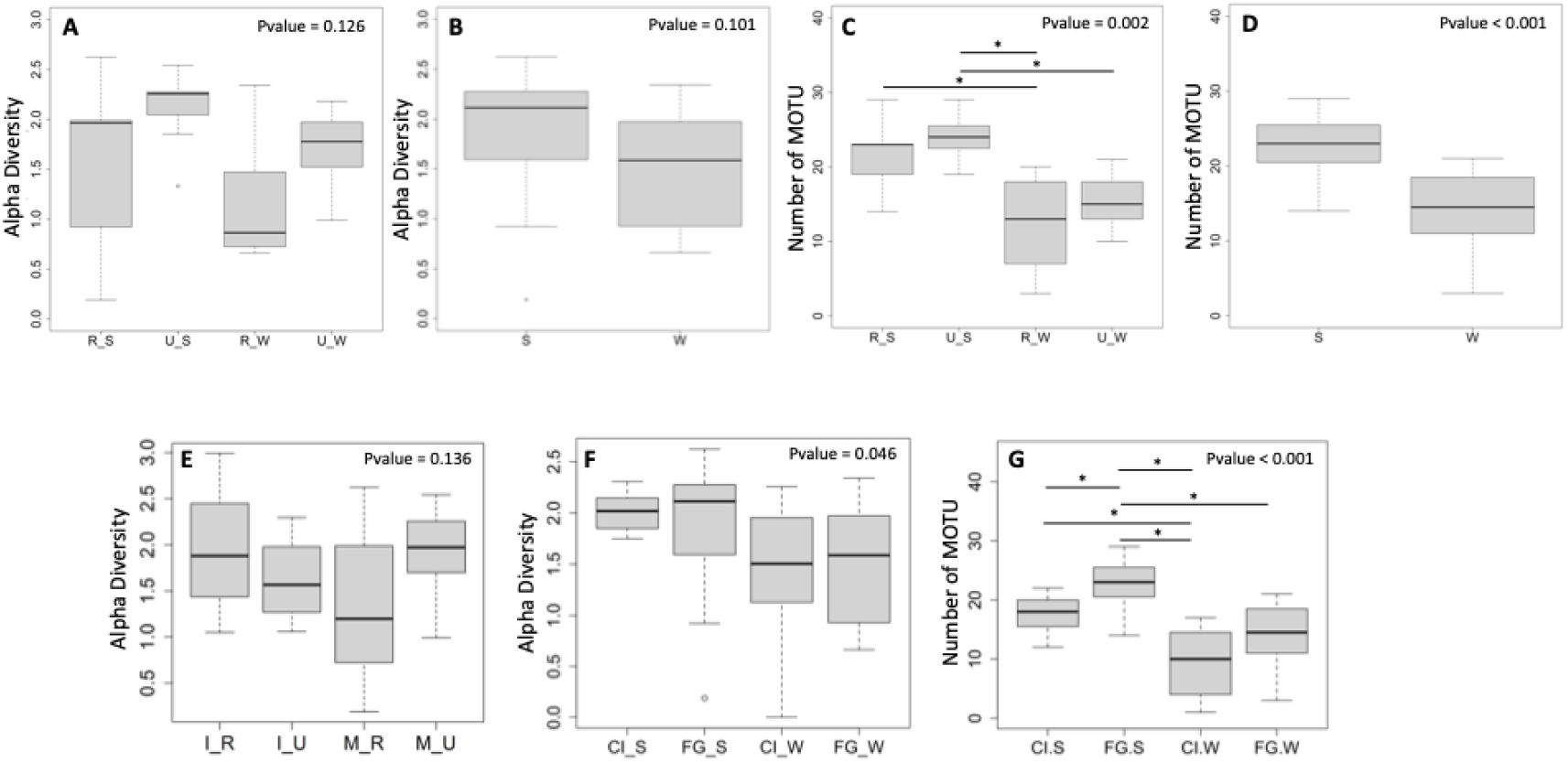
MOTU distribution and diversity in French Guiana (FG). **A:** Alpha (Shannon) diversity for each type of site and for each type of samples. **B:** Alpha (Shannon) diversity for surface sediments and water samples. **C:** Distribution of the number of MOTUs for each type of site and for each type of samples. **D:** Distribution of the number of MOTUs for surface sediments and water samples. **E:** Distribution of alpha diversity (Shannon index) for invertebrates (I) and mycobacteria (M) communities in rural and urban environments. **F & G:** Distribution of alpha diversity (Shannon) (F) and the number of MOTUs (G) by country and by sample type. U: urban sites; R: rural sites; S: surface sediment samples; W: water samples; FG: French Guiana; CI: Côte d’Ivoire. The black bar indicates the median and the moustaches the extremum. Circles indicate extreme values, i.e. values greater than 1.5 times the interquartile space and the segments with * represent significant differences.

#### Vertical Distribution of Mycobacterial MOTUs

Core sampling from four FG sites (Amec, Remi, Cay3, Cri2) revealed no significant differences in MOTU richness across sediment depths (ANOVA, *p* = 0.744; Supplementary Fig. S6), despite a trend of decreasing MOTU numbers with depth. However, MOTU richness varied significantly between sites, with Amec exhibiting lower diversity than Cay3 and Cri2 (Kruskal Wallis, *p* = 0.002; Supplementary Fig. S7). HCA grouped the sites into two clusters: Cay3/Cri2 and Amec/Remi, likely reflecting similar habitat typologies within each group (Supplementary Fig. S8).

PCA confirmed these grouping, indicating distinct mycobacterial communities (Fig. 3C). *M. pallens* (Pall) and *M. haemophilum* (Haem) were preferentially found in Amec/Remi, while *M. avium complex* (Gp13) was more represented in Cay3/Cri2 sediment cores (Fig. 3D). LDA identified Gp13, Pall, and sediment phosphorus (P) as key discriminants between site groups (accuracy = 0.87; Supplementary Fig. S9). MOTU Gp13 varied significantly between (Amec and Cay3, Amec and Cri2, and Cri2 and Remi (ANOVA, *p* = 0.002; Fig. 5A), while phosphorus levels differed between nearly all sites (ANOVA, *p* <0.001; Fig. 5B). The distribution of MOTUs Pall, Vanb, Js62, and Haem also showed significant site-specific differences (Kruskal Walis, *p* < 0.005; Fig. 5C-F). Overall, the clustering of sites Cri2/Cay3 vs. Amec/Remi was supported by both microbial and environmental variables, with site-specific mycobacterial distributions likely driven by local abiotic and biotic factors.

**Figure 5.**
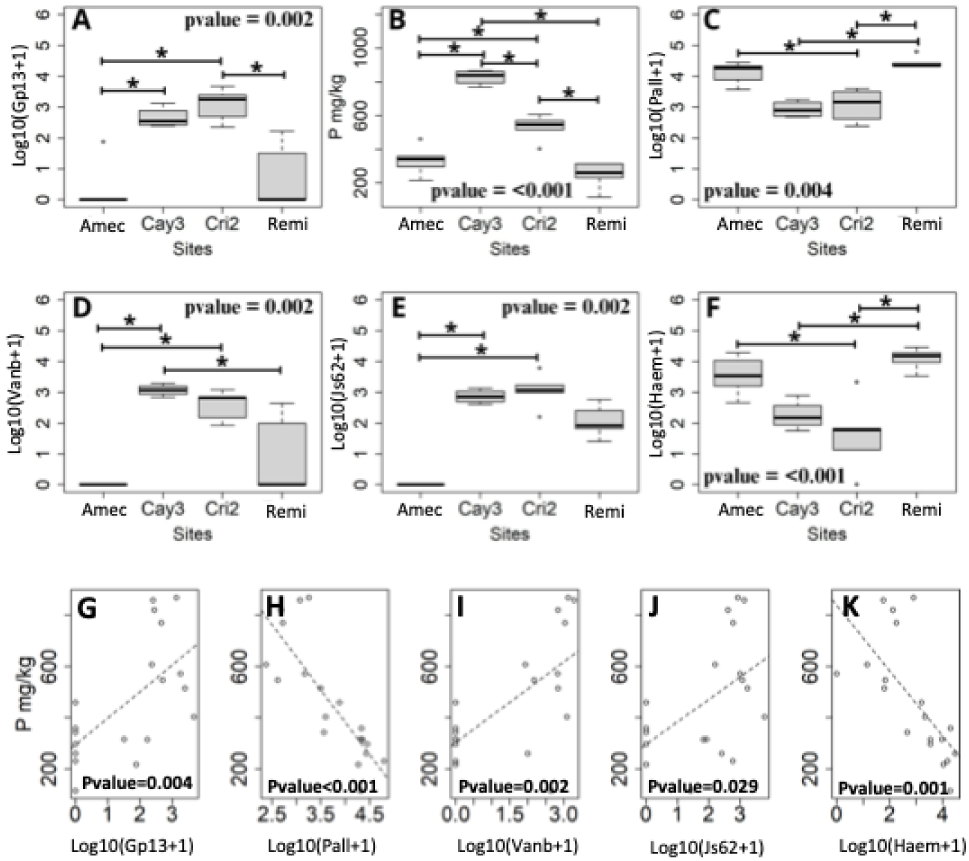
**A-F:** Distribution of the MOTU Gp13, Pall, Vanb, Js62 and Haem and the phosphorus (P) according to the 4 sites sampled for sediment layers (1-2cm, 2-3cm, 3-4cm, 4-5cm, 5-10cm). The black bar indicates the median and the whiskers the extremum. The circle indicates an extreme value i.e. greater than 1.5 times the interquartile range and the segments with “*” represent significant contrasts. **G-K:** Relationship between phosphorus (P) levels and different MOTUs found in the sediment layers (1-2cm, 2-3cm, 3-4cm, 4-5cm, 5-10cm) of four aquatic sites (Amec, Remi, Cri2, Cay3) analyzed in French Guiana (FG).

#### Abiotic Drivers of Mycobacterial MOTU Distribution

To explore abiotic influences on mycobacterial MOTU distribution, we analyzed key physico-chemical parameters in urban and rural sites. Although no significant differences were detected (Wilcoxon, *p* > 0.05), urban sites tended to exhibit higher pH (Supplementary Fig. S10). However, the limited sample size warrant caution in interpretation.

Given the observed structuring of sediment-associated mycobacteria based on phosphorus levels (Fig. 5B, Supplementary Fig. S9), we examined correlations between phosphorus and specific MOTUs. *M. avium* complex (Gp13), Vanb, and Js62 were positively correlated with phosphorus (*p* < 0.05; Fig. 5G, I, J), whereas Pall and Haem showed negative correlations (*p* < 0.05; Fig. 5H, K), suggesting phosphorus availability plays a role in shaping mycobacterial communities.

We further assessed the influence of sediment composition (0-5 cm vs. 5-10 cm) across four sites (Amec, Remi, Cay3, Cri2). Amec and Remi were dominated by coarse sands (> 75%) whereas Cay3 and Cri2 contained mostly clays and fine silts (Supplementary Fig. S11). Random Forest (RF) analysis identified Pall, Haem, and fine silts (F.Slt.) as key variables distinguishing Ame/Rem from Cay3/Cri2 (Supplementary Fig. S12). MOTUs Pall and Haem were associated with coarse sand-rich sites (Amec/Remi) but absent from fine silt/clay-dominated sites (Cay3/Cri2; Supplementary Fig. S13, S14). RF Analysis further indicated that MOTUs Gp01, Diox, and Gilv along with fine sand (F.Sd), differentiated the 0-5 cm and 5-10 cm layers, with all being more prevalent in the upper layer (*p* < 0.05, Supplementary Fig. S15-S17). These results suggest a stratified distribution of mycobacterial communities, with certain MOTUs favoring specific sediment textures and depths.

#### Biotic Drivers of Mycobacterial MOTU Distribution

To explore links between mycobacterial diversity and aquatic invertebrate communities, we conducted a co-inertia analysis (CoIA). Alpha diversity (Shannon index) did not differ significantly between rural and urban sites for either invertebrates or mycobacteria (ANOVA, *p* = 0.136; Fig. 4E), suggesting comparable community diversity across FG. CoIA revealed site-specific patterns in microbial-invertebrate associations. Canonical weights highlighted key invertebrate species and mycobacterial MOTUs contributing to covariance (Fig. 6A), with longer vectors indicating stronger relationships. Site projection (Fig. 6B) showed variables agreement between microbial and invertebrate communities; for example, sites Remi and Pse3 exhibited strong concordance, whereas site Amec showed divergence. The RV coefficient (0.33) indicated weak overall correlation between datasets. These findings suggest that while some sites share similar mycobacterial and invertebrate community structures, there is generally little alignment between the two across French Guiana, indicating different ecological factors may drive their distributions.

**Figure 6.**
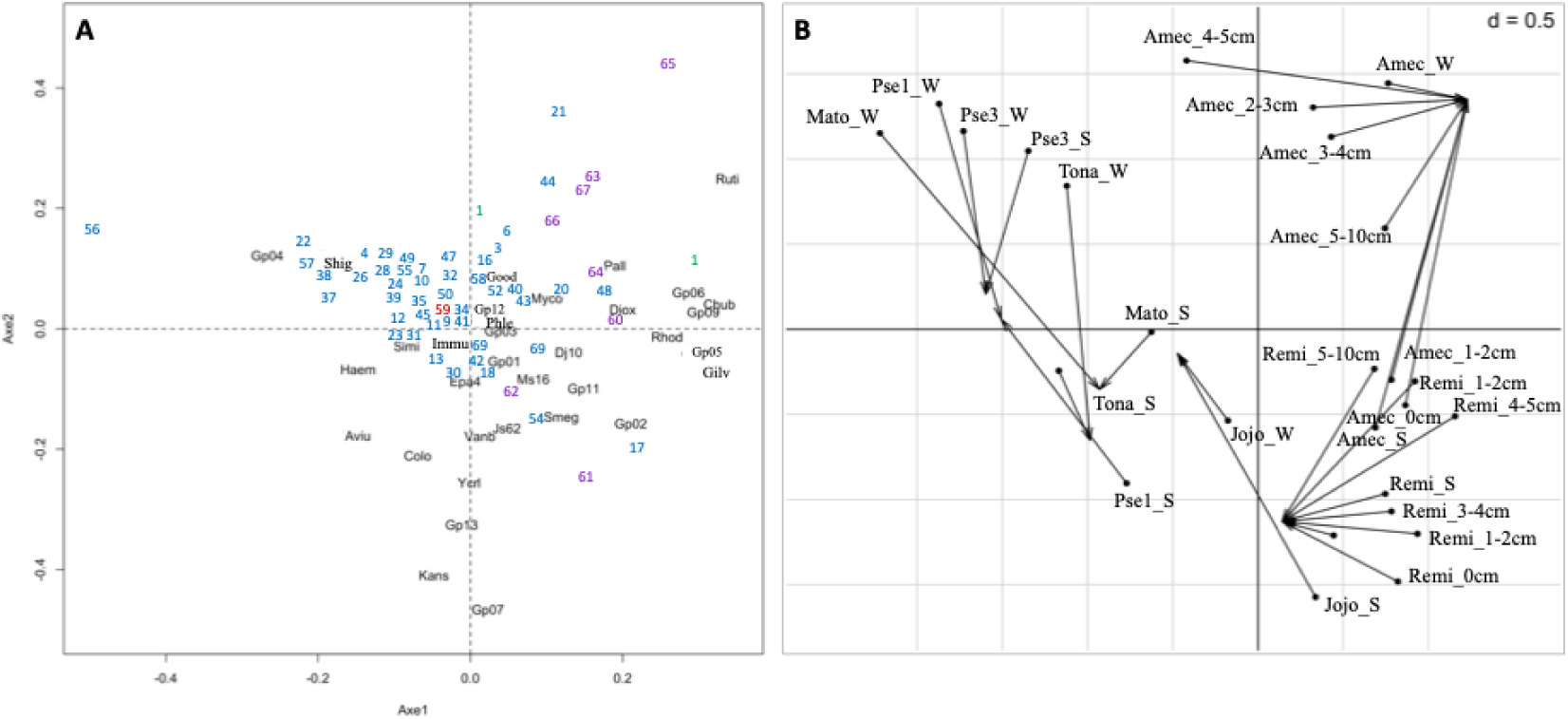
**A:** Biplot of coefficients of environmental variables (Invertebrates) and MOTUs. The plot shows the canonical weights for invertebrate variables (Y axis) and mycobacterial MOTUs (X axis) on the first two axes of co-inertia. Each invertebrate species is represented by a number (See Supplementary Table S2), and the direction and distance from the origin indicate its relative influence on the axes, and each MOTU is represented in black by its abbreviation. Variables with a large length are the most important in explaining the covariance with mycobacterial variables. **B:** Individual-line cloud (Sites). The starting point of the arrow represents the position of the observation in the first data set X, and the tip of the arrow represents its position in the second data set Y.

#### Comparison of Mycobacterial Diversity and Distribution in FG and CI

We assessed whether mycobacterial communities differed between French Guiana (FG) and Côte d’Ivoire (CI). In both countries, most MOTU diversity was captured after sampling ∼ 12 sites (Supplementary Fig. S18), suggesting a limited number of rare species. FG exhibited greater MOTU richness than CI, with an average of 38 vs. 28 MOTUs (Supplementary Fig. S4, S19). Alpha diversity initially showed significant differences between groups (Kruskal Walis test, *p* = 0.046; Fig. 4F), but post hoc tests revealed no significant contrasts. Nonetheless, FG exhibited higher overall MOTU richness than CI (ANOVA, *p* < 0.001; Fig. 4G), with sediments harboring more MOTUs than water samples in both regions. Tukey’s test confirmed that mycobacterial communities varied by geography and environment type. PCA analysis showed clustering of rural and urban sites, with CI communities resembling FG urban sites, except for peri-urban sites Ko01, Ko02, Al01, and Al02 (Fig. 3E). Certain MOTUs, such as Shig, Haem, and Gp04, were associated with rural areas, while Gp02 and Gilv were linked to urban areas (Fig. 3F). Random Forest analysis identified the MOTU Shig as the strongest discriminator between urban and rural sites in both countries (Supplementary Fig. S20). These findings suggest that some mycobacterial MOTUs are habitat-specific, while others are more ubiquitous across FG and CI.

#### Distribution of Potentially Pathogenic Mycobacteria

Several identified MOTUs correspond to mycobacterial species known to cause a human disease (Table 2; Supplementary Table S5). Some species, including *M. intracellulare*, *M. abscessus*, *M. chelonae*, *M. bovis*, *M. terrae*, *M. immunogenum, M. liflandii*, *M. marinum* and *M. ulcerans,* were detected exclusively in French Guina, while *M. fortuitum*, *M. gilvum*, *M. kansasii,* and *M. simiae* were found in both countries (FG and CI). Notably, members of the Mycolactone-Producing-Mycobacteria (MPM) complex (*M. marinum, M. ulcerans, M. liflandii*), which cause Buruli ulcer (BU), were detected at low prevalence in only two FG sites and were absent in CI by eDNA analysis. This contrasts with MPM qPCR assays, which detected three positive sites in FG, including water and sediment samples, but a 100% positivity in CI in sediment samples, while all water samples were negative (data not shown). These discrepancies could 1) question the reliability of qPCR- or eDNA-based detection or 2) suggest that MPMs occur at very low environmental prevalence, potentially affecting transmission likelihood. Additionally, *M. gilvum* was found in both FG and CI across all habitat types (water, surface and core sediment) at high prevalence, indicating its ubiquity in environmental matrices.

#### Patterns of Mycobacterial Interactions

To assess potential associations and exclusions among mycobacterial MOTUs, we constructed partial correlations networks for rural vs urban sites and different sample types (water, surface sediment, sediment core). While partial correlation is closely related to correlation, it shows that a correlation between two variables does not necessarily mean that there is a causality relationship between them, meaning that if there is a correlation between two variables, it may be possible that this correlation can be partially explained by a third variable. In FG rural sites, strong associations where mainly observed between Ruti-Gp06, Ruti-Ycrl, Gp05-Myco and Gp01-Good (0.3 *⩽* ρ *⩽* 1), while the strongest exclusions occurred between Gp03-Ruti, Rhod-Gp06, Gilv-Gp06, and Ms16-Colo (0.06 *⩽* ρ *⩽* 0.57) (Fig. 7A). Urban sites exhibited more and stronger interactions among MOTUs, with notable associations including Gp04-Colo, Gp03-Myco, Haem-Rhod, Haem-Js62, Haem-Vanb, Gp13-Rhod, Gp06-Diox, and Gp08-Gp07, whereas strong exclusions were observed for Epa4-Haem, Gp05-Dj10, and Gp03 and Ms16 (Fig. 7B). Interaction networks also varied by habitat type (Fig. 7C, D, E). In water, key associations included Simi-Ruti, Smeg-Gp13, Myco-Aviu, Good-Gp11, while exclusions involved Kans-Gp04. Surface sediments exhibited strong interactions between Gp08-Gp01, Good-Gilv, and Aviu-Gp07, with exclusions such as Colo-Immu, Immu-Good, Immu-Ms16, Gilv-Gp08, and Gp08-Gp09, among others. In sediment cores, associations included Haem-Pall, Ms16-Js62 and Simi-Gp05, while exclusions occurred for Lito-Ms16 and Gp13-Shig. Overall, interactions were stronger in urban compared to rural sites and more pronounced in surface sediments than in water or sediment cores, correlating with higher MOTU richness in surface sediments. Similar trends were observed in CI, though notable differences emerged, such as Gilv-Ruti, Gilv-Simi, Ruti-Simi, Ruti-Ms16, Myco-Epa4 and Gp09-Gp02, Ycrl-Smeg, Dj10-Js62 being highly exclusive in water and sediment samples, respectively (Fig. 7F, G), while strong associations occurred between Ycrl-Gp06 and Ms16-Kans in water samples vs Rhod-Ruti, Shig-Gp01, Gp11-Dj10, Gp06-Js61, Epa4-Lito in sediment samples. Moreover, while strong associations were observed for a few numbers of MOTUs, we also observed many MOTUs widely distributed within these ecosystems but displaying less pronounced associations with other MOTU and located at the periphery of the network. Again, these findings suggest that mycobacterial interactions occur at the microhabitat level rather than following clear biogeographic patterns.

**Figure 7.**
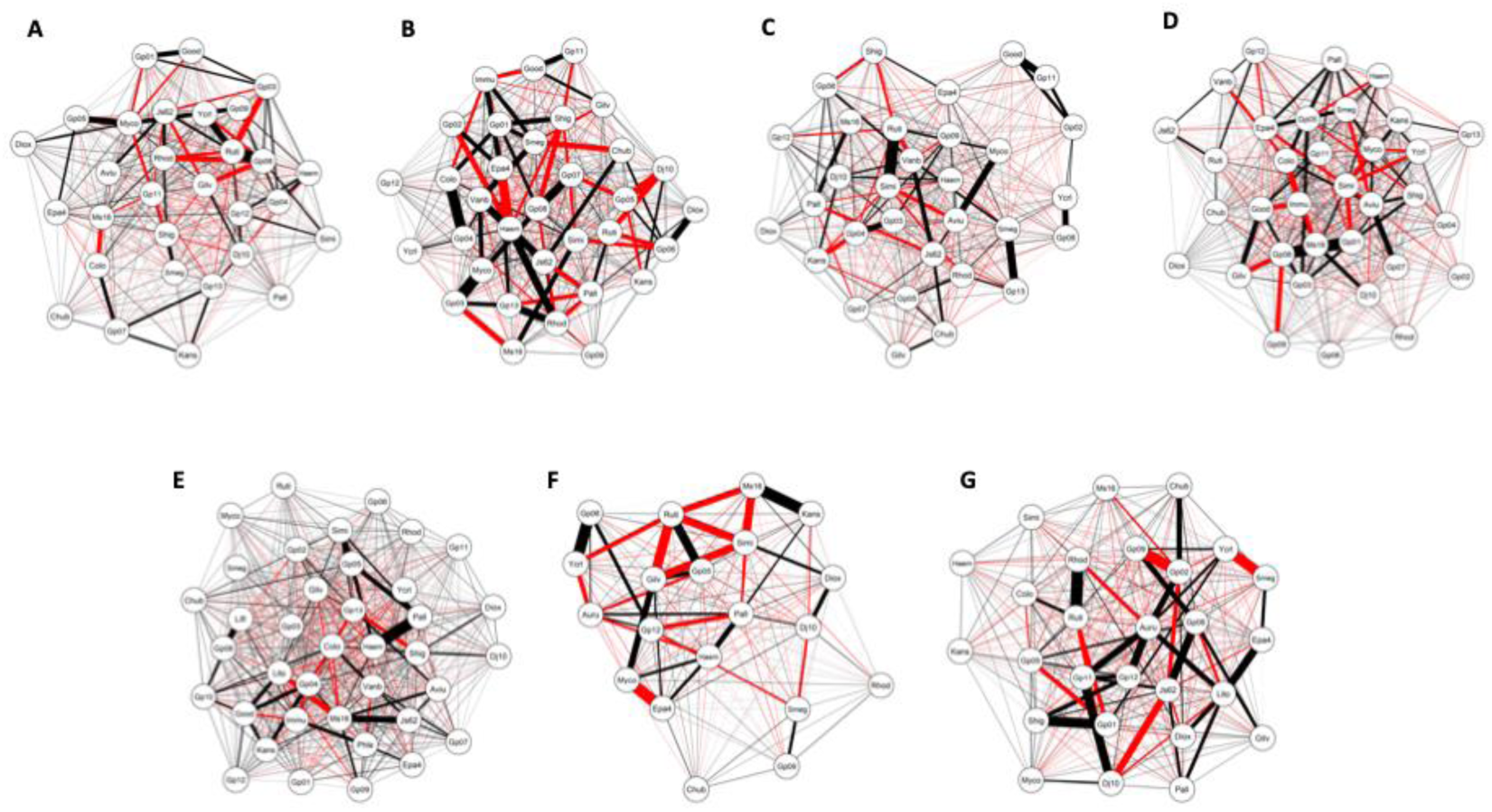
Network analysis of the partial correlation matrix between MOTUs identified in French Guiana (FG) and in Côte d’Ivoire (CI). The red line represents the exclusion links between MOTUs and the black line the associations. The thicker the lines, the more important the links. Here it is a network of partial correlations, the nodes represent the MOTUs and the links represent the partial correlations between these variables, that is to say after control for the other variables of the network. **A**: FG rural sites; **B**: FG urban sites; **C:** FG water samples (including rural and urban sites); **D:** FG surface sediment samples (including rural and urban sites); **E:** FG sediment core samples; **F:** CI water samples; **G:** CI surface sediment samples.

## Discussion

Our study provides novel insights into the diversity, distribution, and ecological interactions of environmental mycobacteria in freshwater habitats of FG and CI. Using high-throughput sequencing, we identified 39/42 mycobacterial MOTUs, including potentially pathogenic species, and demonstrated that their distribution is shaped by both abiotic and biotic factors acting at the microhabitat level. These findings have significant implications for microbial ecology, freshwater ecosystem health, and the potential emergence of zoonotic mycobacterial infections.

### Microbial Ecology and Environmental Partitioning

In FG, mycobacterial community composition varied between urban and rural sites, between water and sediment samples, and across sediment depths. The greater MOTU richness observed in sediments compared to water is consistent with previous findings that sediments provide a stable environment for microbial persistence and biofilm formation (Falkinham, 2009; Tortoli, 2006; Gomez-Alvarez et al., 2021). This is also in line with previous hypothesis that environmental (myco)bacteria, notably pathogenic ones and/or those directly transmitted from the environment, should be more constrained by local habitat characteristics such as water temperature, *pH*, oxygen, salinity, sedimentation and turbidity, biofilm formation, rainfall patterns, etc. (Combe et al., 2019).

HC and PCA analyses confirmed distinct microbial assemblages between urban and rural sites, with urban sediments exhibiting greater taxonomic richness. This could be attributed to increased nutrient availability and anthropogenic influence in urban areas, which may enhance mycobacterial diversity (Li et al., 2025). Although our results did not show a significant difference in the *pH* in urban and rural sediments, the *pH* tended to be higher in urban sediments compared to rural ones, and one could hypothesize that local micro-variations in *pH* may influence the composition of mycobacterial communities. A recent study found that soil *pH* (0-10 cm) tends to be higher in urban and peri-urban areas, which could favor the growth of certain bacterial species better adapted to these conditions (Li et al., 2023). In China, soil microbial richness was found positively correlated with *pH* but negatively correlated with total phosphorus (Jiao et al., 2016). However, concerning *pH* contrasting results were found depending on the studied environment; for instance, in lake sediments *pH* was negatively correlated with microbial richness and phylogenetic diversity (Xiong et al., 2012), while in acid soils *pH* was positively correlated with bacterial alpha diversity (Griffiths et al., 2011). In other cases, bacterial diversity was found to be the highest for near-neutral *pH* (Lauber et al., 2009). Similarly, whilst we did not find any difference between nitrogen levels in urban vs rural areas and MOTU richness, it was showed that moderate levels of nitrogen in soil and sediments enhanced the growth and the diversity of bacteria compared with environment showing high nitrogen levels (Xu et al., 2022; Li et al., 2023). Concerning mycobacteria, *M. smegmatis* (Smeg), *M. bovis* and *M. tuberculosis* (both belonging to Gp04 in the present study) were shown to efficiently assimilate nitrate in laboratory experiments (Khan et al., 2008). In line with our findings, it has been previously showed that urban freshwater habitats in FG, with algae biofilm and watered soils, were more favorable to the establishment of the MPM complex (Combe et al., 2023), including *M. ulcerans* (belonging to Gp08 in the present study) compared with rural ones (Combe et al., 2019).

Whilst DNA sedimentation could have occurred, we believe that this effect was marginal, otherwise we would have observed an accumulation of MOTU DNA through the sediment layers, which was not the case here. Rather, our results are in good agreement with other studies that found lower microbial abundances and biomass in deeper sol layers compared with surface layers (Fierer et al., 2003), and a decrease in the abundance and changes in the composition of microbial communities with sediment depth, notably for the *Mycobacterium* OTU (Myco) (Rissanen et al., 2019). Such observations may be explained in part by the micro-characteristics of these ecosystems, leading to differences in the bioavailability of carbon, nitrogen, oxygen, etc. sources necessary for these microorganisms (Gomez-Alvarez et al., 2021).

Our study showed that phosphorus (P) and substrate type were potentially important abiotic elements in the structuring of mycobacterial communities, with some mycobacteria proliferating under high levels of phosphorous (Gp13), low levels of clays, silts and high levels of coarse sands, while others were found to remain inhibited (Pall, Haem). The positive correlation between phosphorus and *M. avium* complex (Gp13), *M. vanbaaleni* (Vanb), and *Mycobacterium sp. JS623 (*Js62) suggests that nutrient availability may play a critical role in shaping mycobacterial diversity in aquatic ecosystems, and notably in sediment layers (Jiao et al., 2016). Rejas et al. (2005) showed that the reduction in bacterial abundances in depth in tropical freshwaters was mainly due to severe nutrient limitation, especially phosphorus, rather than reduced availability of organic matter. Also, the effect of soil particle size, including clay, silt and sand, were previously proposed to contribute to spatial heterogeneity and bacterial diversity in soils, with nitrifying bacteria exhibiting highest concentrations in fine silt particles (Hemkemeyer et al., 2018). Taken together these findings reinforce the idea that vertical stratification may play a crucial role in structuring microbial communities.

Interestingly, we found that urban sites in FG and CI shared similar mycobacterial communities. This observed clustering patterns further support the hypothesis that local environmental factors, rather than strict biogeographical barriers, drive mycobacterial community structure. It has been proposed that in natural soils stochasticity may remain the dominant process driving the structuration of microbial communities but that oil contamination, for instance, could increase the impact of deterministic processes (Liang et al., 2015). Indeed, oil contamination is well known to affect soil microorganisms due to toxicity and selective pressures (Labud et al., 2007) and increasing competition for nutrients (Margesin et al., 2007). In addition to environmental factors, some studies have proposed that geographic distance may limit the dispersal of microorganisms (Xiong et al., 2014; Liang et al., 2015; Wang et al., 2015; Diniz-Filho & Telles, 2000). For instance, Jiao et al. (2016) found that dispersal limitation played a significant impact in structuring bacterial communities in oil-contaminated environments. However, our findings rather support the hypothesis initially proposed by Baas Becking (1934), highlighting thus that mycobacteria are everywhere in freshwater environments, but, that abiotic parameters occurring at the micro-habitat level trigger either their dormancy or their dominance in the ecosystem.

### Biotic Interactions and Mycobacterial Network Structure

Whilst our results did not show a clear link between mycobacterial communities and those of invertebrates, this lack of direct correlation does not necessarily imply an absence of significant ecological interactions between both communities. Indeed, interactions between bacteria and aquatic invertebrates can be complex. Reported aquatic hosts of mycobacteria involve protozoans, including amoebae, mollusks, crustaceans, cnidarians, echinoderms and sponges (Davidovich et al., 2020). In freshwater ecosystems, MPM have been found associated with a wide range of macro-invertebrate taxa but with higher bacterial loads found in taxa of low and intermediate trophic level (reviewed in Combe et al., 2017). Such wide host-range can be explained by the widespread distribution of mycobacteria in the aquatic environment, explaining in part the absence of correlation between mycobacterial and invertebrate communities. Alternatively, NTM have the ability to exploit ecological niches unoccupied by other (micro)organisms, a characteristic that may also partly explain why we did not observe direct biotic interactions (Davidovich et al., 2020).

Within specific ecological niches, microbial communities form complex interaction networks (Faust & Raes, 2012), with some taxa being highly connected within and between ecological clusters, while other taxa will support lower number of connections within the network (Guimera & Amaral, 2005; Poudel et al., 2016). Moreover, taxa highly connected within and between ecological clusters are expected to support the structure of the network (Faust et al., 2015; Toju et al., 2018b) but also create ecological niche for other taxa (Toju et al., 2018a). Interaction networks are usually used to provide information into the structure and the interaction of complex microbial communities (Barberan et al., 2012) and their importance for investigating the role of microbial communities have been proven in natural ecosystems (Wagg et al., 2019; Delgado-Baquerizo et al., 2020) and for agricultural management through manipulating microbial key taxa (Shi et al., 2020). Network analysis revealed that in FG urban environments foster more and stronger interactions between mycobacterial MOTUs than rural sites, suggesting that simplified urban ecosystems, with low-level trophic networks due to biodiversity loss and human-derived pollution (Combe et al., 2019) would favor the growth of certain MOTU and enhance microbial interactions. Notable associations included notably Gp04-Colo, Gp03-Myco, and Haem-Rhod, while exclusions such as Epa4-Haem and Gp05-Dj10 highlighted competitive dynamics potentially driven by niche differentiation and resource availability (Quiang et al., 2021). Urban environments are well-known to be more suitable to pathogens. For instance, urban sewage water and standing water are known to represent a major reservoir for pathogenic *Leptospira sp.* (Casanovas-Massana et al., 2018; Combe et al., 2019); water-borne and enteric viral and bacterial diseases are at high risk in urban areas with poor sanitation infrastructure (Cable et al., 2017). Interactions also varied by habitat type in FG and in CI, with water samples showing fewer significant interactions than surface sediments. Again, this aligns with the notion that sediment matrices provide a more structured environment conducive to microbial consortia formation (Doloman and Sousa, 2024; Gomez-Alvarez et al. 2021).

Among the many interactions highlighted here, we found that a few number of MOTUs were frequently involved in such interactions, potentially suggesting their importance in the structuring and the functioning of microbial networks and more widely of the ecosystem. It is for instance the case for *M. rutilum* (Ruti), *M. haemophilum* (Haem), *M. immunogenum* (Immu), *M. gilvum* (Gilv) and *M. simiae* (Simi). *M. rutilum* and *M. gilvum*, as other environmental mycobacteria, have the ability to degrade pollutants such as polycyclic aromatic hydrocarbons (PAH), including phenanthrene (PHE) and pyrene (PYR), which are highly toxic, carcinogenic and mutagenic compounds widely distributed in the environment and more frequently encountered in urban soils (Rein et al., 2016). Such abilities suggest that these mycobacteria may function as environmental purifiers, also potentially explaining their occurrence as key species within mycobacterial networks in FG and CI. *M. haemophilum* is an emerging pathogen in humans with a reported unknown natural reservoir. Whole-genome sequence analysis revealed the basis for iron requirements in *M. haemophilum* as well as it is genetic closeness with *M. leprae* (Tufariello et al., 2015). Our study provides thus deeper information about the natural environment of this mycobacteria, found in both countries (FG and CI) but mainly involved in strong interactions in urban sites in FG. Similarly, our study revealed the natural reservoir of *M. simiae,* which remained uncertain in the literature. *M. immunogenum* is well known to be resistant to disinfection procedures, including water chlorination, ozone and addition to biocides, conditions which in fact select for more resistant mycobacterium and provide thus a preferential ecological niche (Primm et al., 2004).

Microbial communities are well known to represent a critical component to the functioning of the ecosystem, playing key roles in nutrient cycling but also food safety and disease emergence (Lin et al., 2023). Microbial communities are also known to interact within ecological niches and these “micro-communities” may function collectively (Hirano & Takemoto, 2019), and exhibit social behaviors such as biofilm formation and quorum sensing as a result of inter- and intra-species interactions ranging from competition for nutrients to cooperative networks (i.e. metabolite exchanges) (Antoniewicz, 2020). Whilst complex interactions within microbial communities often prevent their exact characterization, a better understanding of the nature of these interactions can boost our understanding of their resilience and function (Lin et al., 2023). For instance, it has been proposed that spatially structured microbial communities tend to favor cooperative behaviors (e.g. quorum sensing, commensalims). Contrarily, spatially unstructured or well-mixed communities mainly exhibit competition (e.g. parasitism, amensalism) (Nadell et al., 2016). Whilst our data could not assess the mechanisms of interactions between mycobacterial MOTU, our results revealed positive and negative interactions occurring between mycobacterial communities in freshwater ecosystems, with an observed trend for mycobacterial exclusion. Overall, the interaction matrices showed that mycobacterial communities are extremely diverse in freshwater environments in FG and in CI while some exhibit a distribution pattern specific to the microhabitat, with a few dominant MOTU being specific to certain habitat types versus a large number of low-abundance and/or low-prevalence MOTU being more ubiquitous in these environments, that we can characterize as the “rare biosphere” (Nemergut et al., 2013). In other words, the microhabitat-specific interactions suggest that whilst MOTUs should be find everywhere in freshwater ecosystems, certain mycobacteria may have preferential niches within aquatic environments, likely influenced by competition, predation, and symbiotic relationships.

### Implications for Infectious Disease Emergence

Importantly, our results revealed the widespread presence of several pathogenic mycobacteria, including the *M. avium* complex (Gp13), in freshwater environments, raising concerns about potential human and animal exposure risks. Water is widely recognized to represent the primary source of *M. avium* complex (Gp13) infection in humans and DNA-based fingerprints of *M. avium* isolates from AIDS patients were identical to the isolates found in the patient’s drinking water (Primm et al., 2004). In FG and CI, we found *M. haemophilum* a bacterium causing fish mycobacteriosis (Yanong et al., 2010) but also reported as an emerging mycobacterium causing localized skin infections or even a disseminated disease such as septic arthritis, osteomyelitis and pneumonitis in immunocompromised hosts, notably in children (Lindeboom et al., 2011). Interestingly, while not belonging to the MPM complex usually known to cause Buruli ulcer, the disease caused by *M. haemophilum* most resembles to those of *M. marinum* and *M. ulcerans*, as is the case for its genomic traits compared with other mycobacteria (Lindeboom et al., 2011). Also, some similarities, in terms of fatty acids content and proteic and lipidic properties have been identified between this mycobacterium and *M. leprae* (Besra et al., 1991). Similarly, *M. simiae* is known to cause mycobacteriosis in fish (Toranzo et al., 2004) and cervical lymphadenitis in humans (Cruz et al., 2007). *M. immunogenum*, only found in FG in this study, is known to cause hypersensitive pneumonitis in humans (Selman et al., 2012).

*M. gilvum* (Gilv) is a fast-growing microorganism that was previously found in sputum samples (Stanford & Gunthorpe, 1971), pulmonary tuberculosis infections in Brazil (Mendes de Lima et al., 2013) and in bathing water of leprosy patients in West Bengal (Lavania et al., 2014). Also, *M. gilvum* has been isolated from river sediments based on its ability to degrade polycyclic aromatic hydrocarbons (PAH), such as pyrene, as growth substrate (Lamrabet & Drancourt, 2013). Whilst this mycobacterium was believed to be non-pathogenic, it has been proposed to act as immunomodulator of the host immune system towards susceptibility of other mycobacteria such as *M. leprae* (Lavania et al., 2014). Moreover, *M. gilvum* has been found as single mycobacterium or as co-infections in Buruli ulcer skin biopsies in FG, raising questions about its mechanism of pathogenicity (Combe et al., 2020). Here, the widespread presence of *M. gilvum* across all habitat types (urban, rural, water, sediments) and its high prevalence in FG and in CI *i)* suggest that certain environmental mycobacteria may serve as indicators of ecosystem health and anthropogenic impact and *ii)* raise concerns about opportunistic infections. Overall, given the increasing incidence of NTM infections, particularly in immunocompromised populations, our findings emphasize the importance of monitoring mycobacterial communities in freshwater environments as potential sources of opportunistic infections (Johansen et al., 2020).

Surprisingly, our eDNA screening detected the Mycolactone-Producing-Mycobacteria (MPM) complex, which include closely related variants such as *M. ulcerans*, *M. shinshuense*, *M. pseudoshottsii, M. liflandii* and *M. marinum* (Combe et al., 2023), at very low environmental prevalence and only in FG (MOTUs Lifl and Gp08 only). However, all these variants have the ability to cause Buruli ulcer and were previously found in patient biopsies in Japan, CI and FG (reviewed in Combe et al., 2023). Whilst Buruli ulcer remains a neglected disease, its importance is no less since it affects nearly 5000-6000 people annually across 33 countries and represents the third most common mycobacterial disease after tuberculosis and leprosy (reviewed in Combe et al., 2023). Consequently, our results suggest that transmission dynamics of the causative mycobacterial agents causing Buruli ulcer may involve complex ecological reservoirs or undetected amplification hosts. This finding is particularly relevant given the endemicity of Buruli ulcer in West Africa (Tabah et al. 2019).

In Brazil, Mexico, the Caribbean and the USA *M. lepromatosis*, a closely related bacilli to *M. leprae*, has been reported to cause leprosy, or Hansen’s disease (Schaub et al., 2020). *M. lepromatosis* does not belong to the NTM, but rather is an obligate intracellular parasite (Schaub et al., 2020). This pathogen is mainly transmitted from human to human, probably through nasal droplets, or by contact with contaminated bathing water or cultivated soil sources, but its potential zoonotic transmission has also been proposed (Schaub et al., 2020). However, our eDNA-based metabarcoding analyses did not reveal the presence of *M. lepromatosis* in environmental samples in FG and CI, suggesting that these studied ecosystems may not represent the environmental reservoir of this mycobacterium.

Overall, the greater MOTU richness found in urban sediment samples, and the discrepancy between microbial assemblages between urban and rural areas, may have important implication for public health and urbanization. For instance, it is well known that soil represents a reservoir for the *M. avium* complex (Gp13) associated with human infection, and that individuals whose occupation involves prolonged exposure to soil are at increased risk of infection (Reed et al., 2006). Similarly, our current understanding of Buruli ulcer disease relies on higher prevalence of the MPM variants in urban environments and notably in sediment samples (Combe et al., 2019). In fact, freshwater habitats represent the natural reservoir of the mycobacterium responsible for Buruli ulcer which are spread into new habitats during flooding. However, human-disturbed environments, following for instance deforestation, agriculture and urbanization and leading to stagnant water bodies with increased water temperature, decreased *pH* and oxygen levels, algae growth, biofilm formation and also exhibiting sudden changes in foodweb structure and function, are at higher risk of disease emergence (reviewed in Combe et al., 2017). Similarly, other mycobacterium such as *M. avium* (Aviu) and *M. intracellulare* (Gp10) exhibit an acidic *pH* optimum comprised between 4.5-5.5 (George and Falkinham, 1986), and their growth is enhanced by humic and fulvic acids, the major compounds found in brown water swamps (Kirschner et al., 1999). Higher bacterial loads of *M. avium* and *M. intracellulare* have been found from water and soils of low oxygen levels (Brooks et al., 1984; Kirschner et al., 1992). Together these findings strengthen the fact that such abiotic and biotic conditions, prone to increased virulence and pathogenicity (Primm et al., 2004), are most frequently encountered in urban areas where human exposure is also enhanced.

In addition to abiotic and biotic preferential niches, human-mediated pollution spilled in urban aquatic habitats may explain the observation of many mycobacterial species known for their ability to degrade pollutants such as PAH, such as *M. rutilum* (Ruti), *M. gilvum* (Gilv), *M. haemophilum* (Haem) or *M. immunogenum* (Immu), among others. Moreover, the presence of a greater richness of mycobacterial species in urban areas, together with the pollution from other bacteria and the higher use of antimicrobial drugs (antibiotics) and disinfectant (chlorine, ozone, biocides) compared with rural areas, represent selective environments that boost either the acquisition or the expression of resistance genes by these opportunistic – and usually non-pathogenic – mycobacteria (Norton and LeChevallier, 2000). Indeed, the prevalence of environmental mycobacteria in municipal drinking water supplies is directly explained by their high innate chlorine, ozone and biocide resistance (Le Dantec et al., 2002; Norton and LeChevallier, 2000; Taylor et al., 2000). For comparison, even the weaker *M. aurum* mycobacterium is 100-fold more resistant to chlorine than *Escherichia coli* (Le Dantec et al., 2002). Along with their ability for biofilm formation, these bacilli can easily persist in flowing systems, such as water distribution system (Primm et al., 2004), even under starvation conditions or extreme temperatures, as for instance experienced in hot tap water, spas and ice machines (Nyka, 1974; Schulze-Robbecke and Buchholtz, 1992; Smeulders et al., 1999).

A deeper understanding of mycobacterial distribution patterns in the environment could help to identify hotspots of infectious disease risk in order to guide targeted surveillance programmes. Interestingly, while qPCR assays detected the MPM complex in CI, our eDNA-based sequencing approach did not, highlighting thus potential limitations in environmental DNA detection methodologies. These discrepancies underscore the need for multi-marker approaches to improve pathogen surveillance in aquatic ecosystems (Mve-Obiang et al., 2003). Importantly, in order to prevent aquatic mycobacterial diseases emergence in tropical environments we raise the importance of 1) improved infrastructure and sanitation, including access to drinking water and soap for daily disinfection of wounds, 2) access to health care facilities for treatment, 3) monitoring the mycobacterial species present in water supplies along with their chlorine susceptibility, 4) regulation of pollutant use (antibiotics, biocides, PAH, etc.) along with 5) the implementation of biodiversity restoration policies within urban and peri-urban zones which clearly represent “infectious bubbles” at high risk of disease emergence (Combe & Gozlan, 2024). Such implementations would also align with the One health strategy being currently widely adopted at a worldwide scale.

## Conclusions

This study advances our understanding of environmental mycobacteria ecology by demonstrating the influence of habitat type, abiotic factors, and microbial interactions on mycobacterial distribution. The detection of pathogenic species further highlights the importance of environmental reservoirs in mycobacterial disease transmission. Future research should focus on integrating metagenomics and functional analyses to elucidate the ecological roles of mycobacteria in freshwater ecosystems and their potential public health implications. Longitudinal studies incorporating climate variability and land-use changes may also provide further insights into the environmental drivers of mycobacterial diversity and their relevance to zoonotic disease emergence.

## Acknowledgements

This work was supported by the Centre d’Étude de la Biodiversité Amazonienne (LabEx CEBA MicroBiomes) in French Guiana. MC received a postdoctoral fellow from the Agence Nationale de la Recherche (ANR-17-CE35-0006-01 PRIME). We acknowledge Dr. Laurent Marsollier (ATOMYCA, Université d’Angers, France) and Dr. Jean-Christophe Avarre (UMR ISEM, IRD, Université de Montpellier, France) for providing pure DNA for several mycobacterial species that we used as positive controls in this study. We also thank Dr. Michel Brossard, the IRD representative in French Guiana, for facilitating the logistics for sampling. Finally, we thank Mr. Rolland Ruffine for his help in sampling in French Guiana.

## References

Amha, Y. M., Anwar, M. Z., Kumaraswamy, R., Henschel, A., & Ahmad, F. (2017). Mycobacteria in municipal wastewater treatment and reuse: microbial diversity for screening the occurrence of clinically and environmentally relevant species in arid regions. Environmental science & technology, 51(5), 3048–3056. 10.1021/acs.est.6b05580

Antoniewicz, M. R. (2020). A guide to deciphering microbial interactions and metabolic fluxes in microbiome communities. Current opinion in biotechnology, 64, 230–237. 10.1016/j.copbio.2020.07.001

Barberan, A., Bates, S.T., Casamayor, E.O., Fierer, N., (2012). Using network analysis to explore co-occurrence patterns in soil microbial communities. ISME Journal, 6, 343e351. 10.1038/ismej.2011.119

Bass, D., Stentiford, G.D., Wang, H-C., Koskella, B. Tyler, C.R. (2019). The Pathobiome in Animal and Plant Diseases. Trends in Ecology & Evolution, 34(11): 996–1008. 10.1016/j.tree.2019.07.012

Becking, L. B. (1934). Geobiologie of inleiding tot de milieukunde (No. 18-19). WP Van Stockum & Zoon.

Bellemain, Carlsen, Brochmann, Coissac, Taberlet, & Kauserud. (2010). ITS as an environmental DNA barcode for fungi: an in silico approach reveals potential PCR biases. BMC Microbiology, 10(1), 1–9. 10.1186/1471-2180-10-189

Besra, G. S., McNeil, M., Minnikin, D. E., Portaels, F., Ridell, M., & Brennan, P. J. (1991). Structural elucidation and antigenicity of a novel phenolic glycolipid antigen from Mycobacterium haemophilum. Biochemistry, 30(31), 7772–7777.

Boyer, F., Mercier, C., Bonin, A., Le Bras, Y., Taberlet, P., Coissac, E. (2016). OBITOOLS: A UNIX -inspired software package for DNA metabarcoding. Molecular Ecology Resources, 16(1), 176–182. 10.1111/1755-0998.12428

Brooks, R.W., George, K.L., Parker, B.C., Falkinham, J.O., Gruff, G. (1984). Recovery and survival of nontuberculous mycobacteria under various growth and decontamination conditions. Canadian Journal of Microbiology, 30: 1112–1117. 10.1139/m84-174

Cable, J., Barber, I., Boag, B., Ellison, A. R., Morgan, E. R., Murray, K., Pascoe, E. L., Sait, S. M., Wilson, A. J., Booth, M. (2017). Global change, parasite transmission and disease control: lessons from ecology. Philosophical Transactions of the Royal Society B: Biological Sciences, 372(1719), 20160088. 10.1098/rstb.2016.0088

Cangelosi, G., Clark-Curtis, J., Behr, M., Bull, T., Stinear, T. (2004) Biology of waterborne pathogenic mycobacteria. In Pathogenic mycobacteria in water. A guide to public health consequences, monitoring and management. In Emerging issues in water and infectious disease series. Edited by S. Pedley, J. Bartram, G. Rees, A. Dufour, J.A. Cotruvo. World Health Organization by IWA Publishing.

Casanovas-Massana, A., Costa, F., Riediger, I.N., Cunha, M., Oliveira, D.D, Mota, D.C., et al. (2018). Spatial and temporal dynamics of pathogenic Leptospira in surface waters from the urban slum environment. Water Research, 130: 176–184. 10.1016/j.watres.2017.11.068

Chen, W. C. W., Ng TzeHann, N.T., Wu JerHorng, W.J., Chen JiungWen, C.J., Wang HanChing, W.H. (2017). Microbiome dynamics in a shrimp grow-out pond with possible outbreak of acute hepatopancreatic necrosis disease. Scientific Reports, 7(1), 9395. 10.1038/s41598-017-09923-6

Clare, E.L., Economou, C.K., Faulkes, C.G., Gilbert, J.D., Bennett, F., Drinkwater, R., Littlefair, J.E. (2021). eDNAir: proof of concept that animal DNA can be collected from air sampling. PeerJ 9:e11030 10.7717/peerj.11030

Combe, M., Cherif, E., Blaizot, R., Breugnot, D., Gozlan, R. E. (2023). What about current diversity of mycolactone-producing mycobacteria? Implication for the diagnosis and treatment of buruli ulcer. International Journal of Molecular Sciences, 24(18), 13727. 10.3390/ijms241813727

Combe, M., Couppié, P., Blaizot, R., Valentini, A., Gozlan, R. E. (2020). Are all Buruli ulcers caused by Mycobacterium ulcerans? British Journal of Dermatology, 183(5), 968–970. 10.1111/bjd.19260

Combe, M., Gozlan, R. E. (2024). When the Blue Marble Health concept challenges our current understanding of One Health. One Health, 19, 100935. 10.1016/j.onehlt.2024.100935

Combe M, Gozlan RE, Jagadesh S, Velvin CJ, Ruffine R, Demar MP, et al. (2019) Comparison of Mycobacterium ulcerans (Buruli ulcer) and Leptospira sp. (Leptospirosis) dynamics in urban and rural settings. PLoS Negl Trop Dis 13(1): e0007074. 10.1371/journal.pntd.0007074

Combe M., Velvin C.J., Morris A., Garchitorena A., Carolan K., Sanhueza D., Roche B., Couppié P., Guégan J-F., Gozlan R.E. (2017). Global and local environmental changes as drivers of Buruli ulcer emergence. Emerging Microbes and Infections, 6: e20. 10.1038/emi.2017.7

Crooks, G.E., Hon, G., Chandonia, J.M., Brenner, S.E. (2004). WebLogo: a sequence logo generator. Genome research, 14(6), 1188–1190. 10.1101/gr.849004

Cruz, A. T., & Palazzi, D. L. (2007). An adolescent with fever, cough, & diffuse lymphadenopathy. The Pediatric infectious disease journal, 26(3), 275. 10.1097/01.inf.0000256756.70575.4d

Curtis, T. P., Sloan, W. T., Scannell, J. W. (2002). Estimating prokaryotic diversity and its limits. Proceedings of the National Academy of Sciences, 99(16), 10494–10499. 10.1073/pnas.142680199

Daszak, P., Cunningham, A.A., Hyatt, A.D. (2000). Emerging infectious diseases of wildlife – threats to biodiversity and human health. Science, 287(5452), 443–449. https://doi.10.1126/science.287.5452.44

Davidovich, N., Morick, D., Carella, F. (2020). Mycobacteriosis in aquatic invertebrates: A review of its emergence. Microorganisms, 8(8), 1249. 10.3390/microorganisms8081249

De Groote, M.A., Johnson, P. (2004) Pathogenic mycobacteria in water. A guide to public health consequences, monitoring and management. In Emerging issues in water and infectious disease series. Edited by S. Pedley, J. Bartram, G. Rees, A. Dufour, J.A. Cotruvo. World Health Organization by IWA Publishing.

Delgado-Baquerizo, M., Reich, P.B., Trivedi, C., Eldridge, D.J., Abades, S., Alfaro, F.D., Bastida, F., Berhe, A.A., Cutler, N.A., Gallardo, A., Garcia-Velazquez, L., Hart, S.C., Hayes, P.E., He, J.Z., Hseu, Z.Y., Hu, H.W., Kirchmair, M., Neuhauser, S., Perez, C.A., Reed, S.C., Santos, F., Sullivan, B.W., Trivedi, P., Wang, J.T., Weber-Grullon, L., Williams, M.A., Singh, B.K., (2020). Multiple elements of soil biodiversity drive eco-system functions across biomes. Nature in Ecology and Evolution, 4, 210–220. 10.1038/s41559-019-1084-y

Derome, N., Gauthier, J., Boutin, S., Llewellyn, M. (2016). Bacterial Opportunistic Pathogens of Fish. In: Hurst, C. (eds) The Rasputin Effect: When Commensals and Symbionts Become Parasitic. Advances in Environmental Microbiology, vol 3. Springer, Cham. 10.1007/978-3-319-28170-4_4

Diniz-Filho, J.A.F., Telles, M.P.d.C., (2000). Spatial pattern and genetic diversity estimates are linked in stochastic models of population differentiation. Genetics and Molecular Biology, 23, 541e544. 10.1590/S1415-47572000000300007

Doig, K. D., Holt, K. E., Fyfe, J. A., Lavender, C. J., Eddyani, M., Portaels, F., Yeboah-Manu, D., Pluschke, G., Seemann, T., & Stinear, T. P. (2012). On the origin of Mycobacterium ulcerans, the causative agent of Buruli ulcer. BMC Genomics, 13(1), 258. 10.1186/1471-2164-13-258

Doloman, A., Sousa, D.Z. (2024). Mechanisms of microbial co-aggregation in mixed anaerobic cultures. Applied Microbiology and Biotechnology, 108(1):407. 10.1007/s00253-024-13246-8

Douine M., Gozlan R.E., Nacher M., Dufour J., Reynaud Y., Elguero E., Combe M., Velvin C.J., Chevillon C., Berlioz-Arthaud A., Labbé S., Sainte-Marie D., Guégan J.F., Pradinaud R., Couppié P. (2017). Mycobacterium ulcerans infection (Buruli ulcer) in French Guiana, South America, 1969-2013: an epidemiologic study. The Lancet Planetary Health, 1: e65–73. 10.1016/S2542-5196(17)30009-8

Drummond, A. J., Newcomb, R. D., Buckley, T. R., Xie, D., Dopheide, A., Potter, B. C., Heled, J., Ross, H. A., Tooman, L., Grosser, S., Park, D., Demetras, N. J., Stevens, M. I., Russell, J. C., Anderson, S. H., Carter, A., Nelson, N. (2015). Evaluating a multigene environmental DNA approach for biodiversity assessment. GigaScience, 4(1), 46. 10.1186/s13742-015-0086-1

Epskamp, S., Cramer, A. O., Waldorp, L. J., Schmittmann, V. D., Borsboom, D. (2012). qgraph: Network visualizations of relationships in psychometric data. Journal of statistical software, 48, 1–18. 10.18637/jss.v048.i04

Falkinham, J.O. (2002). Nontuberculous mycobacteria in the environment. Clinics in Chest Medicine, 23(3), 529–551. 10.1016/S0272-5231(02)00014-X

Falkinham, J.O. (2009). Surrounded by mycobacteria: Nontuberculous mycobacteria in the human environment. Journal of Applied Microbiology, 107(2), 356–367. 10.1111/j.1365-2672.2009.04161.x

Faust, K., Lima-Mendez, G., Lerat, J.S., Sathirapongsasuti, J.F., Knight, R., Huttenhower, C., Lenaerts, T., Raes, J., (2015). Cross-biome comparison of microbial association networks. Frontiers in Microbiology 6, 1200. 10.3389/fmicb.2015.01200

Faust, K., Raes, J., (2012). Microbial interactions: from networks to models. Nature Reviews Microbiology, 10, 538e550. 10.1038/nrmicro2832

Ficetola, G. F., Coissac, E., Zundel, S., Riaz, T., Shehzad, W., Bessière, J., Taberlet, P., Pompanon, F. (2010). An In silico approach for the evaluation of DNA barcodes. BMC Genomics, 11(1), 1–10. 10.1186/1471-2164-11-434

Fierer, N., Jackson, R. B. (2006). The diversity and biogeography of soil bacterial communities. Proceedings of the National Academy of Sciences, 103(3), 626–631. 10.1073/pnas.050753510

Fierer, N., Schimel, J. P., Holden, P. A. (2003). Variations in microbial community composition through two soil depth profiles. Soil Biology and Biochemistry, 35(1), 167–176. 10.1016/S0038-0717(02)00251-1

George, K.L., Falkinham III, J.O. (1986). Selective medium for the isolation and enumeration of Mycobacterium avium-intracellulare and M. scrofulaceum. Canadian journal of microbiology, 32(1), 10–14. 10.1139/m86-003

Gomez-Alvarez, V., Liu, H., Pressman, J. G., & Wahman, D. G. (2021). Metagenomic profile of microbial communities in a drinking water storage tank sediment after sequential exposure to monochloramine, free chlorine, and monochloramine. ACS ES&T Water, 1(5), 1283–1294. 10.1021/acsestwater.1c00016.

Griffiths, R.I., Thomson, B.C., James, P., Bell, T., Bailey, M., Whiteley, A.S., (2011). The bacterial biogeography of British soils. Environmental Microbiology, 13, 1642e1654. 10.1111/j.1462-2920.2011.02480.x

Guimera, R., Amaral, L.A.N., (2005). Functional cartography of complex metabolic networks. Nature, 433, 895–900. 10.1038/nature03288

Hammoudi, N., Dizoe, S., Saad, J., Ehouman, E., Davoust, B., Drancourt, M., Bouam, A. (2020). Tracing Mycobacterium ulcerans along an alimentary chain in Côte d’Ivoire: A one health perspective. PLoS Neglected Tropical Disease, 14(5): e0008228. 10.1371/journal.pntd.0008228.

Hemkemeyer, M., Dohrmann, A.B., Christensen, B.T., Tebbe, C.C. (2018). Bacterial preferences for specific soil particle size fractions revealed by community analyses. Frontiers in Microbiology, 9, 149. 10.3389/fmicb.2018.00149

Hirano, H., Takemoto, K. (2019). Difficulty in inferring microbial community structure based on co-occurrence network approaches. BMC Bioinformatics, 20:329. 10.1186/s12859-019-2915-1

Horner-Devine M.C., Lage M., Hughes J.B., Bohannan J.M. (2004). A taxa-area relationship for bacteria. Nature, 432: 750–753. 10.1038/nature03073

Jagadesh, S., Combe, M., Couppié, P., Le Turnier, P., Epelboin, L., Nacher, M., Gozlan, R. E. (2019). Emerging human infectious diseases of aquatic origin: A comparative biogeographic approach using Bayesian spatial modelling. International Journal of Health Geographics, 18(1), 23. 10.1186/s12942-019-0188-6

Jagadesh, S., Combe, M., Couppié, P., Nacher, M. Gozlan, R.E. (2019). Global Emergence of Buruli Ulcer. EcoHealth 16, 591–593 (2019). 10.1007/s10393-019-01445-z

Jiao, S., Chen, W., Wang, E., Wang, J., Liu, Z., Li, Y., Wei, G., (2016). Microbial Succession in Response to Pollutants in Batch-enrichment Culture. Scientific reports, 6(1), 21791. 10.1038/srep21791

Jiao, S., Liu, Z., Lin, Y., Yang, J., Chen, W., Wei, G. (2016). Bacterial communities in oil contaminated soils: biogeography and co-occurrence patterns. Soil Biology and Biochemistry, 98, 64–73. 10.1016/j.soilbio.2016.04.005

Johansen, M.D., Herrmann, J.L., Kremer, L. (2020). Non-tuberculous mycobacteria and the rise of Mycobacterium abscessus. Nature Reviews Microbiology, 18(7): 392–407. 10.1038/s41579-020-0331-1

Keesing, F., Holt, R. D., Ostfeld, R. S. (2006). Effects of species diversity on disease risk. Ecology Letters, 9(4), 485–498. 10.1111/j.1461-0248.2006.00885.x

Keesing F, Ostfeld R.S. (2021). Dilution effect in disease ecology. Ecology Letters, 24(11): 2490–2505. 10.1111/ele.13875

Khan, A., Akhtar, S., Ahmad, J. N., Sarkar, D. (2008). Presence of a functional nitrate assimilation pathway in Mycobacterium smegmatis. Microbial Pathogenesis, 44(1), 71–77. 10.1016/j.micpath.2007.08.006

Kirschner, R.A., Parker, B.C., Falkinham, J.O. (1999). Humic and fulvic acids stimulate the growth of Mycobacterium avium. FEMS Microbiology Ecology, 30:327–332. 10.1111/j.1574-6941.1999.tb00660.x

Kox, L. F., Van Leeuwen, J., Knijper, S., Jansen, H. M., Kolk, A. H. (1995). PCR assay based on DNA coding for 16S rRNA for detection and identification of mycobacteria in clinical samples. Journal of Clinical Microbiology, 33(12), 3225–3233. 10.1128/jcm.33.12.3225-3233.1995

Labrie, L., Komar, C., Terhune, J., Camus, A., & Wise, D. (2004). Effect of Sublethal Exposure to the Trematode Bolbophorus spp. on the Severity of Enteric Septicemia of Catfish inChannel Catfish Fingerlings. Journal of Aquatic Animal Health, 16(4), 231–237. 10.1577/H04-011.1

Labud, V., Garcia, C., Hernandez, T., (2007). Effect of hydrocarbon pollution on the microbial properties of a sandy and a clay soil. Chemosphere 66, 1863e1871. 10.1016/j.chemosphere.2006.08.021

Lai, H. C., Ng, T. H., Ando, M., Lee, C. T., Chen, I. T., Chuang, J. C., … Wang, H. C. (2015). Pathogenesis of acute hepatopancreatic necrosis disease (AHPND) in shrimp. Fish & shellfish immunology, 47(2), 1006–1014. 10.1016/j.fsi.2015.11.008

Lamrabet, O., Drancourt, M. (2013). Mycobacterium gilvum illustrates size-correlated relationships between mycobacteria and Acanthamoeba polyphaga. Applied Environmental Microbiology, 79(5): 1606–1611. 10.1128/AEM.03765-12

Lauber, C.L., Hamady, M., Knight, R., Fierer, N., (2009). Pyrosequencing-based assessment of soil pH as a predictor of soil bacterial community structure at the continental scale. Applied & Environmental Microbiology, 75, 5111e5120. 10.1128/AEM.00335-09

Lavania, M., Turankar, R., Singh, I., et al. (2014). Detection of Mycobacterium gilvum first time from the bathing water of leprosy patient from Purulia, West Bengal. International Journal of Mycobacteriology, 3(4): 286–289. 10.1016/j.ijmyco.2014.10.005

Le Dantec, C., Duguet, J. P., Montiel, A., Dumoutier, N., Dubrou, S., & Vincent, V. (2002). Chlorine disinfection of atypical mycobacteria isolated from a water distribution system. Applied and environmental microbiology, 68(3), 1025–1032. 10.1128/AEM.68.3.1025-1032.2002

Li, M., Chen, L., Zhao, F., Tang, J., Bu, Q., Wang, X., & Yang, L. (2023). Effects of urban–rural environmental gradient on soil microbial community in rapidly urbanizing area. Ecosystem Health and Sustainability, 9, 0118. 10.34133/ehs.01188

Li, J., Zheng, Y., Achal, V. (2025). Impact of anthropogenic activities on bacterial community diversity in a coastal city: A case study from Shantou. Continental Shelf Research, 289, 105450. 10.1016/j.csr.2025.105450.

Liang, Y., Zhang, X., Zhou, J., Li, G., (2015). Long-term oil contamination increases deterministic assembly processes in soil microbes. Ecological Applications, 25, 1235e1243. 10.1890/14-1672.1

Lin, L., Du, R., Wu, Q., Xu, Y. (2023). Metabolic cooperation between conspecific genotypic groups contributes to bacterial fitness. ISME communications, 3:1–11. 10.1038/s43705-023-00250-8

Lin, Q., Li, L., Adams, J. M., Heděnec, P., Tu, B., Li, C., … & Li, X. (2021). Nutrient resource availability mediates niche differentiation and temporal co-occurrence of soil bacterial communities. Applied Soil Ecology, 163, 103965. 10.1016/j.apsoil.2021.103965

Lindeboom, J.A., Bruijnesteijn van Coppenraet, L.E., van Soolingen, D., Prins, J.M., Kuijper, E.J. (2011). Clinical manifestations, diagnosis, and treatment of Mycobacterium haemophilum infections. Clinical microbiology reviews, 24(4), 701–717. 10.1128/cmr.00020-11

Loss, A.G. (2012). The Black Queen Hypothesis: evolution of dependencies through adaptive gene loss. MBio 3, e00036–12.

Margesin, R., Hämmerle, M., Tscherko, D. (2007). Microbial activity and community composition during bioremediation of diesel-oil-contaminated soil: effects of hydrocarbon concentration, fertilizers, and incubation time. Microbial Ecology, 53, 259–269. 10.1007/s00248-006-9136-7

Mendes de Lima, C.A, Magdinier Gomes, H., Cardoso Oelemann, M.A., et al. (2013). Nontuberculous mycobacteria in respiratory samples from patients with pulmonary tuberculosis in the state of Rondônia, Brazil. Memorias do Instituto Oswaldo Cruz, 108(4): 457–462. 10.1590%2F0074-0276108042013010

Morris, A., Gozlan, R., Marion, E., Marsollier, L., Andreou, D., Sanhueza, D., Ruffine, R., Couppié, P., Guégan, J.-F. (2014). First detection of mycobacterium ulcerans dna in environmental samples from south america. PLoS Neglected Tropical Diseases, 8(1), e2660. 10.1371/journal.pntd.0002660

Morris, A. L., Guégan, J.-F., Andreou, D., Marsollier, L., Carolan, K., Le Croller, M., Sanhueza, D., Gozlan, R. E. (2016). Deforestation-driven food-web collapse linked to emerging tropical infectious disease, Mycobacterium ulcerans. Science Advances, 2(12), e1600387. 10.1126/sciadv.1600387

Mosser, T., Talagrand-Reboul, E., Colston, S. M., Graf, J., Figueras, M. J., Jumas-Bilak, E., Lamy, B. (2015). Exposure to pairs of Aeromonas strains enhances virulence in the Caenorhabditis elegans infection model. Frontiers in Microbiology, 6, 1218. 10.3389/fmicb.2015.01218

Mve-Obiang, A., Lee, R.E., Portaels, F., Small, P.L. (2003). Heterogeneity of mycolactones produced by clinical isolates of Mycobacterium ulcerans: implications for virulence. Infection and Immunity, 71(2):774–83. 10.1128/IAI.71.2.774-783.2003

Nadell, C.D., Drescher, K., Foster, K.R. (2016) Spatial structure, cooperation and competition in biofilms. Nature Reviews in Microbiology, 14:589–600. 10.1038/nrmicro.2016.84

Nemergut, D.R., Schmidt, S.K., Fukami, T., O’Neill, S.P., Bilinski, T.M., Stanish, L.F., Knelman, J.E., Darcy, J.L., Lynch, R.C., Wickey, P., Ferrenberg, S. (2013). Patterns and processes of microbial community assembly. Microbiology and Molecular Biology Reviews, 77(3), 342− 356. 10.1128/mmbr.00051-12

Nguetta, A., Coulibaly, N.D., Kouamé-Elogne, N.C., Acquah, K.J.R., Amon, A.C., Kouamé, K., Konan, N., Koffi, A., Kadio, M.C., Yao, A., et al. (2018). Phenotypic and Genotypic Characterization of Mycobacteria Isolates from Buruli Ulcer Suspected Patients Reveals the Involvement of Several Mycobacteria in Chronic Skin Lesions. American Journal of Microbiology Researches, 6, 79–87. 10.12691/ajmr-6-3-3

Nichols, G., Ford, T., Bartram, J., Dufour, A., Portaels, F. (2004) Pathogenic mycobacteria in water. A guide to public health consequences, monitoring and management. In Emerging issues in water and infectious disease series. Edited by S. Pedley, J. Bartram, G. Rees, A. Dufour, J.A. Cotruvo. World Health Organization by IWA Publishing.

Norton, C.D., LeChevallier, M.W. (2000). A pilot study of bacteriological population changes through potable water treatment and distribution. Applied and Environmental Microbiology, 66:268–276. 10.1128/AEM.66.1.268-276.2000

Nuñez, N.F., Maggia, L., Stenger, P-L., Lelievre, M., Letellier, K., Gigante, S., Manez, A., Mournet, P., Ripoll, J., Carriconde, F. (2021). Potential of high-throughput eDNA sequencing of soil fungi and bacteria for monitoring ecological restoration in ultramafic substrates: The case study of the New Caledonian biodiversity hotspot. Ecological Engineering, 173, 106416. 10.1016/j.ecoleng.2021.106416

Nyka, W. (1974). Studies on the effect of starvation on mycobacteria. Infection and immunity, 9(5), 843–850. 10.1128/iai.9.5.843-850.1974

Olds, B. P., Jerde, C. L., Renshaw, M. A., Li, Y., Evans, N. T., Turner, C. R., Deiner, K., Mahon, A. R., Brueseke, M. A., Shirey, P. D., Pfrender, M. E., Lodge, D. M., Lamberti, G. A. (2016). Estimating species richness using environmental DNA. Ecology and Evolution, 6(12), 4214–4226. 10.1002/ece3.2186

Oloya, J., Opuda-Asibo, J., Kazwala, R., Demelash, A. B., Skjerve, E., Lund, A., Johansen, T. B., Djonne, B. (2008). Mycobacteria causing human cervical lymphadenitis in pastoral communities in the Karamoja region of Uganda. Epidemiology and Infection, 136(5), 636–643. 10.1017/S095026880700900

Otta, S. K., Arulraj, R., Ezhil Praveena, P., Manivel, R., Panigrahi, A., Bhuvaneswari, T., … & Ponniah, A. G. (2014). Association of dual viral infection with mortality of Pacific white shrimp (Litopenaeus vannamei) in culture ponds in India. Virus Disease, 25, 63–68. 10.1007/s13337-013-0180-x

Peleg, A. Y., Tampakakis, E., Fuchs, B. B., Eliopoulos, G. M., Moellering Jr, R. C., & Mylonakis, E. (2008). Prokaryote–eukaryote interactions identified by using Caenorhabditis elegans. Proceedings of the National Academy of Sciences, 105(38), 14585–14590. 10.1073/pnas.0805048105

Phung, T.N., Caruso, D., Godreuil, S., Keck, N., Vallaeys, T., & Avarre, J.-C. (2013). Rapid detection and identification of nontuberculous mycobacterial pathogens in fish by using high-resolution melting analysis. Applied and Environmental Microbiology, 79(24), 7837–7845. 10.1128/AEM.00822-13

Pitlik, S.D., Koren, O. (2017). How holobionts get sick–toward a unifying scheme of disease. Microbiome 5, 64. 10.1186/s40168-017-0281-7

Pont, D., Rocle, M., Valentini, A., Civade, R., Jean, P., Maire, A., Roset, N., Schabuss, M., Zornig, H., Dejean, T. (2018). Environmental DNA reveals quantitative patterns of fish biodiversity in large rivers despite its downstream transportation. Scientific Reports, 8(1), 10361. 10.1038/s41598-018-28424-8

Pontiroli, A., Khera, T. T., Oakley, B. B., Mason, S., Dowd, S. E., Travis, E. R., … Wellington, E. M. (2013). Prospecting environmental mycobacteria: combined molecular approaches reveal unprecedented diversity. PLoS One, 8(7), e68648. 10.1371/journal.pone.0068648

Poudel, R., Jumpponen, A., Schlatter, D., Paulitz, T., Gardener, B.M., Kinkel, L., Garrett, K., (2016). Microbiome networks: a systems framework for identifying candidate microbial assemblages for disease management. Phytopathology, 106, 1083–1096. 10.1094/PHYTO-02-16-0058-FI

Primm, T.P., Lucero, C.A., Falkinham, J.O. (2004). Health impacts of environmental mycobacteria. Clinical Microbiology Review, 17(1): 98–106. 10.1128/CMR.17.1.98-106.2004

Radomski, N., Roguet, A., Lucas, F. S., Veyrier, F. J., Cambau, E., Accrombessi, H., … Moulin, L. (2013). atpE gene as a new useful specific molecular target to quantify Mycobacterium in environmental samples. BMC microbiology, 13, 1–13. 10.1186/1471-2180-13-277

Reed, C., Von Reyn, C.F., Chamblee, S., Ellerbrock, T.V., Johnson, J.W., Marsh, B.J., … Horsburgh Jr, C.R. (2006). Environmental risk factors for infection with Mycobacterium avium complex. American journal of epidemiology, 164(1), 32–40. 10.1093/aje/kwj159

Rejas, D., Muylaert, K., De Meester, L. (2005). Nutrient limitation of bacteria and sources of nutrients supporting nutrient-limited bacterial growth in an Amazonian floodplain lake. Aquatic microbial ecology, 39(1), 57–67

Rein, A., Adam, I. K., Miltner, A., Brumme, K., Kästner, M., & Trapp, S. (2016). Impact of bacterial activity on turnover of insoluble hydrophobic substrates (phenanthrene and pyrene) - model simulations for prediction of bioremediation success. Journal of Hazardous Materials, 306, 105–114. 10.1016/j.jhazmat.2015.12.005

Reverter, M., Sarter, S., Caruso, D., Avarre, J. C., Combe, M., Pepey, E., Pouyaud, L., Vega-Heredía, S., de Verdal, H., Zambonino-Infante, J. L. (2020). Aquaculture at the crossroads of global warming and antimicrobial resistance. Nature Communications, 11, 1870. 10.1038/s41467-020-15735-6

Riaz, T., Shehzad, W., Viari, A., Pompanon, F., Taberlet, P., Coissac, E. (2011). ecoPrimers: inference of new DNA barcode markers from whole genome sequence analysis, 39(21), e145–e145. 10.1093/nar/gkr732

Rissanen, A. J., Peura, S., Mpamah, P. A., Taipale, S., Tiirola, M., Biasi, C., … Nykänen, H. (2019). Vertical stratification of bacteria and archaea in sediments of a small boreal humic lake. FEMS Microbiology Letters, 366(5), fnz044. 10.1093/femsle/fnz044

Schaub, R., Avanzi, C., Singh, P. et al. (2020). Leprosy Transmission in Amazonian Countries: Current Status and Future Trends. Current Tropical Medicine Reports, 7, 79–91. 10.1007/s40475-020-00206-1

Schulze-Robbecke, R., Buchholtz, K. (1992). Heat susceptibility of aquatic mycobacteria. Applied and environmental microbiology, 58:1869–1873. 10.1128/aem.58.6.1869-1873.1992

Selman, M., Pardo, A., & King Jr, T. E. (2012). Hypersensitivity pneumonitis: insights in diagnosis and pathobiology. American journal of respiratory and critical care medicine, 186(4), 314–324. 10.1164/rccm.201203-0513CI

Shi, Y., Delgado-Baquerizo, M., Li, Y., Yang, Y., Zhu, Y. G., Peñuelas, J., & Chu, H. (2020). Abundance of kinless hubs within soil microbial networks are associated with high functional potential in agricultural ecosystems. Environment International, 142, 105869. 10.1016/j.envint.2020.105869

Smeulders, M.J., Keer, J., Speight, R.A., Williams, H.D. (1999). Adaptation of Mycobacterium smegmatis to stationary phase. Journal of Bacteriology, 181: 270–283. 10.1128/jb.181.1.270-283.1999

Stackebrandt E., Goebel B.M. (1994). Taxonomic note: A place for DNA-DNA reassociation and 16S rRNA sequence analysis in the present species definition in bacteriology. International Journal of Systematic and Microbiology, 44: 846–849. 10.1099/00207713-44-4-846

Stanford, J.L., Gunthorpe, W.J. (1971). A study of some fast-growing scotochromogenic mycobacteria including species descriptions of Mycobacterium gilvum (new species) and Mycobacterium duvalii (new species). British Journal of Experimental Pathology, 52: 627–637.

Stat, M., Huggett, M. J., Bernasconi, R., DiBattista, J. D., Berry, T. E., Newman, S. J., Harvey, E. S., Bunce, M. (2017). Ecosystem biomonitoring with eDNA: metabarcoding across the tree of life in a tropical marine environment. Scientific Reports, 7(1), 12240. 10.1038/s41598-017-12501-5

Straus, W.L., Ostroff, S.M., Jemigan, D.B., Kiehn, T.E., Sordillo, EM., Armstrong, D., Boone, N., Schneider, N., Kilburn, Silcox, V.A., LaBombardi, V., Good, R.C. (1994). Clinical and epidemiologic characteristics of Mycobacterium haemophilum, an emerging pathogen in immunocompromised patients. Annals of Internal Medicine, 120, 118–125. 10.7326/0003-4819-120-2-199401150-00004

Stubbendieck, R. M., Straight, P. D. (2016). Multifaceted interfaces of bacterial competition. Journal of Bacteriology, 198(16), 2145–2155. 10.1128/jb.00275-16

Tabah, E. N., Johnson, C. R., Degnonvi, H., Pluschke, G., Röltgen, K. (2019). Buruli ulcer in Africa. Buruli Ulcer: Mycobacterium Ulcerans Disease, 43-60. 10.1007/978-3-030-11114-4_2

Taylor, R.H., Falkinham, J.O., Norton, C.D., LeChevallier, M.W. (2000). Chlorine, chloramine, chlorine dioxide, and ozone susceptibility of Mycobacterium avium. Applied and environmental microbiology, 66:1702–1705. 10.1128/AEM.66.4.1702-1705.2000

Tobler, Pfunder, Herzog, Frey, Altwegg. (2006). Rapid detection and species identification of Mycobacterium spp. using real-time PCR and DNA-Microarray. Journal of Microbiological Methods, 66(1), 116–124. 10.1016/j.mimet.2005.10.016

Toju, H., Peay, K.G., Yamamichi, M., Narisawa, K., Hiruma, K., Naito, K., Fukuda, S., Ushio, M., Nakaoka, S., Onoda, Y., Yoshida, K., Schlaeppi, K., Bai, Y., Sugiura, R., Ichihashi, Y., Minamisawa, K., Kiers, E.T., (2018). Core microbiomes for sustainable agroecosystems. Nature Plants, 4, 247–257. 10.1038/s41477-018-0139-4

Toju, H., Tanabe, A.S., Sato, H., (2018). Network hubs in root-associated fungal meta-communities. Microbiome, 6, 116. 10.1186/s40168-018-0497-1

Toranzo, A.E. (2004). Report about fish bacterial diseases. Mediterranean Aquaculture Laboratories, 49-89.

Tortoli, E. (2006). The new mycobacteria: an update. FEMS Immunology & Medical Microbiology, 48, 159–178. 10.1111/j.1574-695X.2006.00123.x

Tufariello, J.M., Kerantzas, C.A., Vilchèze, C., Calder, R.B., Nordberg, E.K., Fischer, J.A., Hartman, T.E., Yang, E., Driscoll, T., Cole, L.E., Sebra, R,. Maqbool, S.B., Wattam, A.R., Jacobs, W.R. (2015). The Complete Genome Sequence of the Emerging Pathogen Mycobacterium haemophilum Explains Its Unique Culture Requirements. mBio, 6(6):e01313–15. 10.1128/mBio.01313-15

Valentini, A., Taberlet, P., Miaud, C., Civade, R., Herder, J., Thomsen, P. F., Bellemain, E., Besnard, A., Coissac, E., Boyer, F., Gaboriaud, C., Jean, P., Poulet, N., Roset, N., Copp, G. H., Geniez, P., Pont, D., Argillier, C., Baudoin, J., … Dejean, T. (2016). Next-generation monitoring of aquatic biodiversity using environmental DNA metabarcoding. Molecular Ecology, 25(4), 929–942. 10.1111/mec.13428

Wagg, C., Schlaeppi, K., Banerjee, S., Kuramae, E.E., van der Heijden, M.G.A., (2019). Fungal-bacterial diversity and microbiome complexity predict ecosystem functioning. Nature Communication, 10, 4841. 10.1038/s41467-019-12798-y

Wang, K., Ye, X., Chen, H., Zhao, Q., Hu, C., He, J., Qian, Y., Xiong, J., Zhu, J., Zhang, D., (2015). Bacterial biogeography in the coastal waters of northern Zhejiang, East China Sea is highly controlled by spatially structured environmental gradients. Environmental Microbiology, 17, 3898e3913. 10.1111/1462-2920.12884

Wit, R., Bouvier, T. (2006). Everything is everywhere, but, the environment selects; what did Baas Becking and Beijerinck really say? Environmental Microbiology, 8(4), 755–758. 10.1111/j.1462-2920.2006.01017.x

Wu, X., Lu, Y., Zhou, S., Chen, L., Xu, B. (2016). Impact of climate change on human infectious diseases: Empirical evidence and human adaptation. Environment International, 86, 14–23. 10.1016/j.envint.2015.09.007

Xiong, J., Liu, Y., Lin, X., Zhang, H., Zeng, J., Hou, J., Yang, Y., Yao, T., Knight, R., Chu, H., (2012). Geographic distance and pH drive bacterial distribution in alkaline lake sediments across Tibetan Plateau. Environmental Microbiology, 14, 2457e2466. 10.1111/j.1462-2920.2012.02799.x

Xiong, J., Ye, X., Wang, K., Chen, H., Hu, C., Zhu, J., Zhang, D., (2014). Biogeography of the sediment bacterial community responds to a nitrogen pollution gradient in the East China Sea. Applied & Environmental Microbiology, 80, 1919e1925. 10.1128/AEM.03731-13

Xu, Y. F., Dong, X. M., Luo, C., Ma, S. N., Xu, J. L., Cui, Y. D. (2022). Nitrogen enrichment reduces the diversity of bacteria and alters their nutrient strategies in intertidal zones. Frontiers in Marine Science, 9, 942074. 10.3389/fmars.2022.942074

Yanong, R.P.E., Pouder, D.B., Falkinham, J.O. (2010). Association of mycobacteria in recirculating aquaculture systems and mycobacterial disease in fish. Journal of Aquatic Animal Health, 22(4), 219–223. 10.1577/H10-009.1

