## Supplementary Information for "Ecological Drivers of Nontuberculous Mycobacteria in Aquatic Systems: Biodiversity, Niche Competition, and Pathogen Emergence"

**Supplementary Figure S1.** Some aquatic sites sampled in French Guiana (FG). A: Amec; B: Remi; C: Cri2; D: Cay3. See Table 1 for site names.

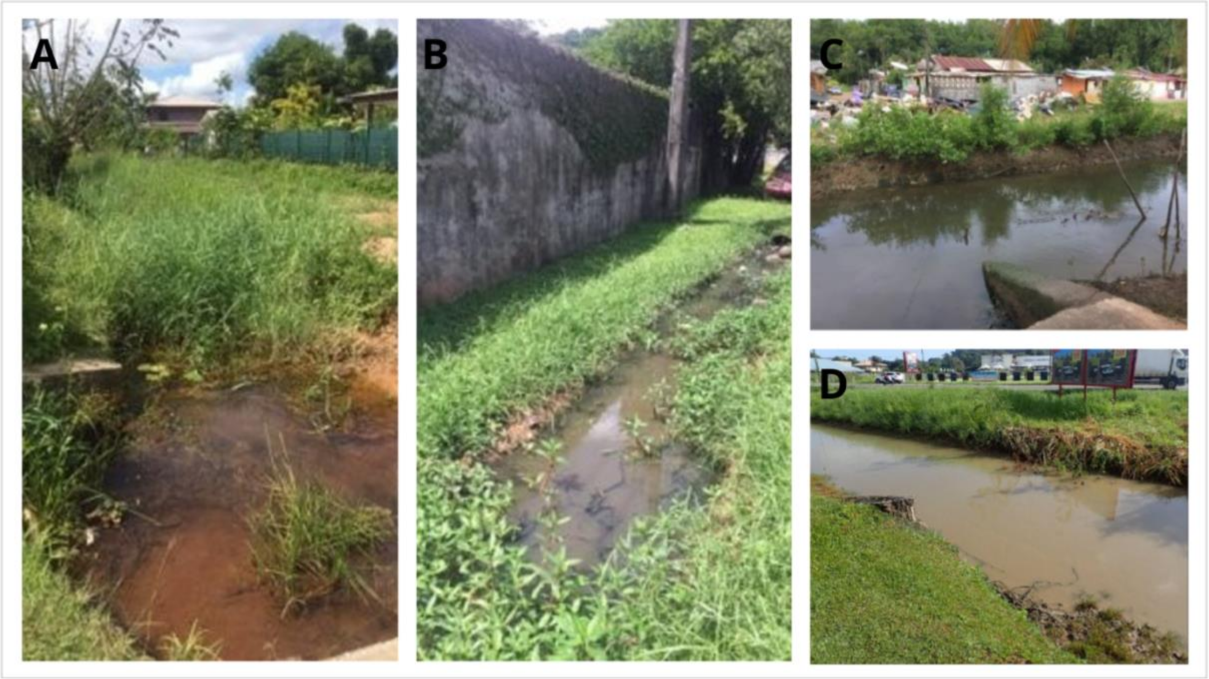

**Supplementary Figure S2.** Some aquatic sites sampled in Côte d'Ivoire (CI). A: ALO01; B: BO01; C: KO01; D: GEO3. See Table 1 for site names.

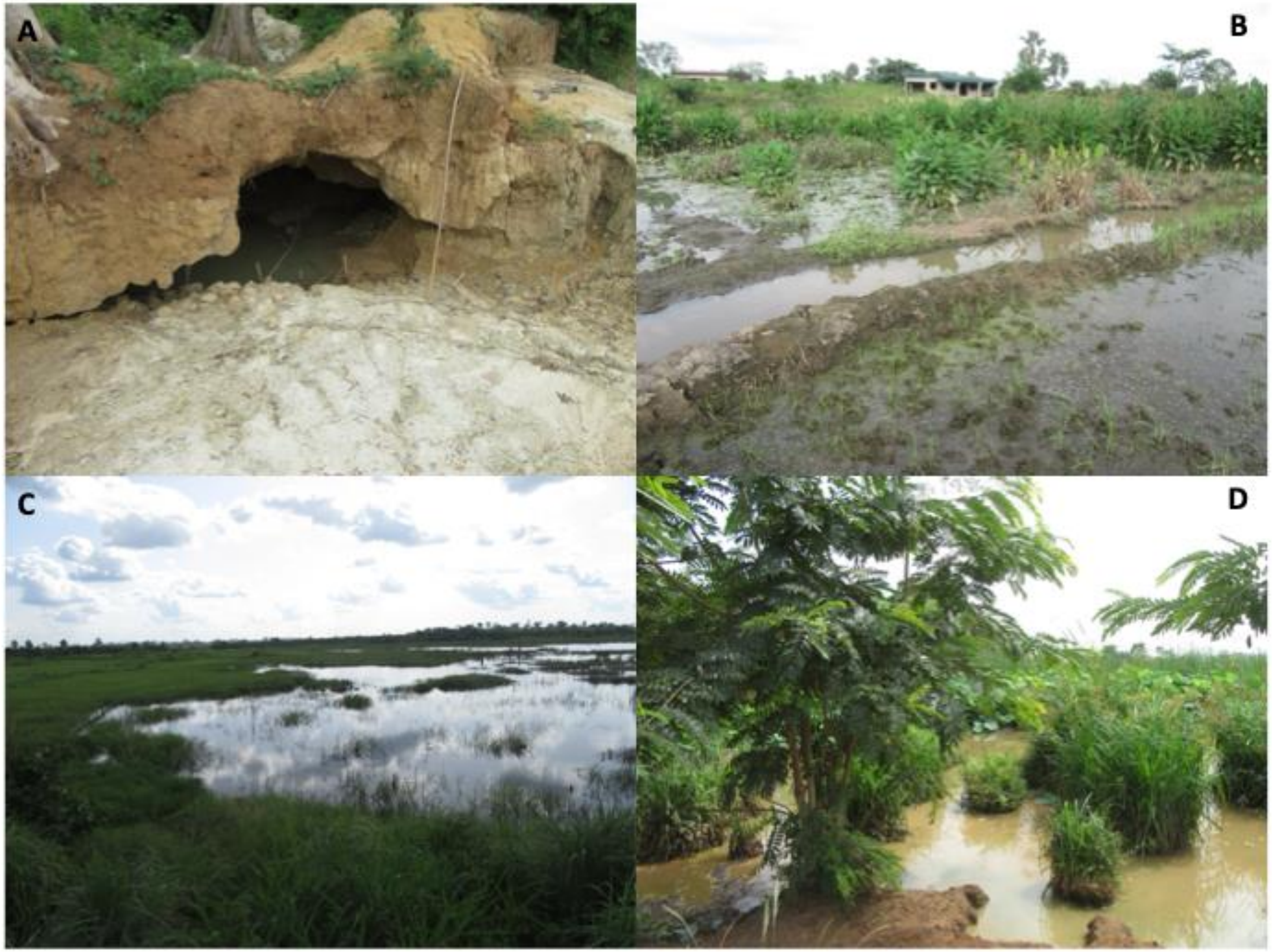

**Supplementary Figure S3.** Reads generated by Illumina MiSeq. The total number of reads is represented by the black lines whilst the red lines show the number of reads assigned to a NTM MOTU after applying the 97% similarity treshold. **A**: reads in water and surface sediment samples in French Guiana; **B**: reads in sediment cores in French Guiana; **C**: reads in water and surface sediment in Côte d'Ivoire.

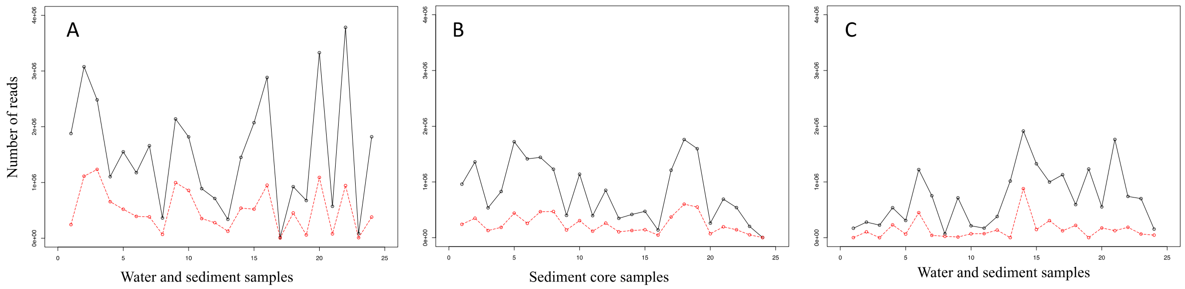

**Supplementary Figure S4.** Ven diagram of the MOTU found in FG and CI.

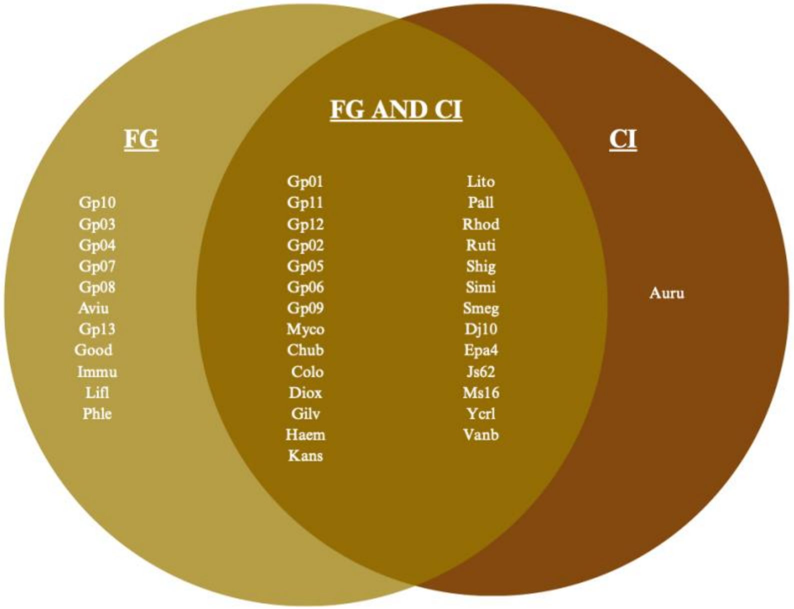

**Supplementary Figure S5.** Score (area under ROC curve) of the contributions of each variable (mycobacterial MOTUs) in the LDA analysis. The higher the score for a MOTU, the more important it is for distinguishing between the two groups of sites observed (rural and urban sites).

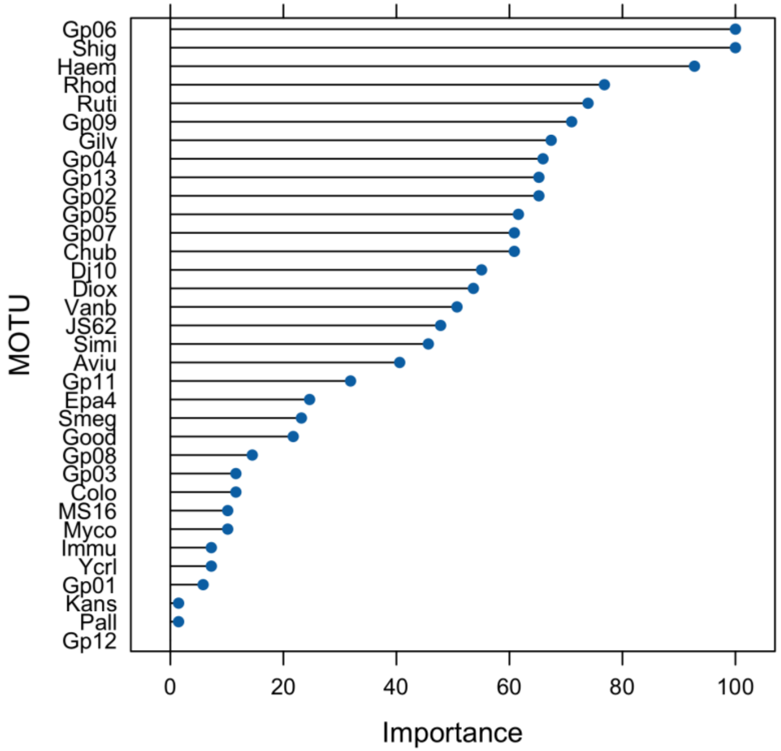

**Supplementary Figure S6.** Distribution of the number of MOTUs per core layer for the 4 sites combined (Amec, Remi, Cay3, Cri2). The Cay3_5-10 cm value was not representative and was thus excluded from the analysis. The anova test performed shows no significative differences in the number of MOTU between sediment layers. The black bar indicates the median and the whiskers the extremum. The circle indicates an extreme value, i.e. greater than 1.5 times the interquartile range.

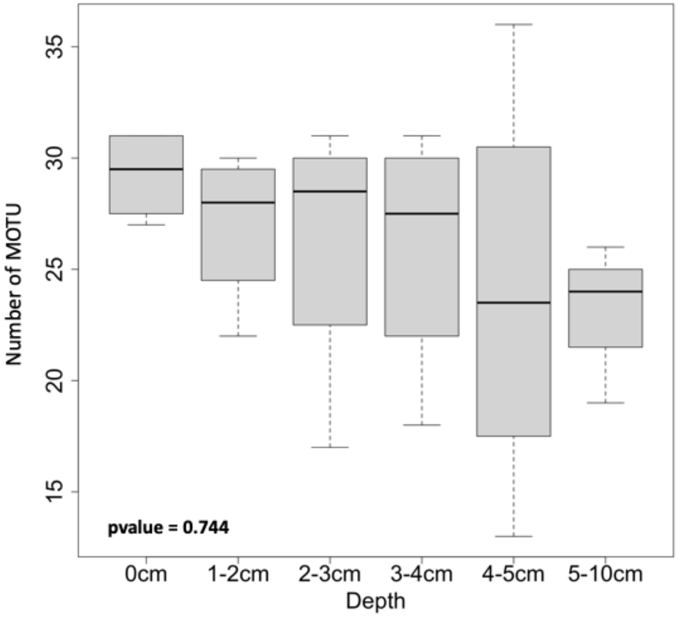

**Supplementary Figure S7.** Distribution of the number of MOTUs per site. The Cay3_5-10cm value was excluded as it was not representative. For the 4 sites, all sediment layers (0-1cm, 1-2cm, 2-3cm, 3-4cm, 4-5cm, 5-10cm) were combined. The Kruskal Wallis test shows differences between the site Amec and the sites Cay3 and Cri2. The black bar indicates the median and the whiskers the extremum. The circle indicates an extreme value i.e. greater than 1.5 times the interquartile range and the segments with "*" represent significant contrasts.

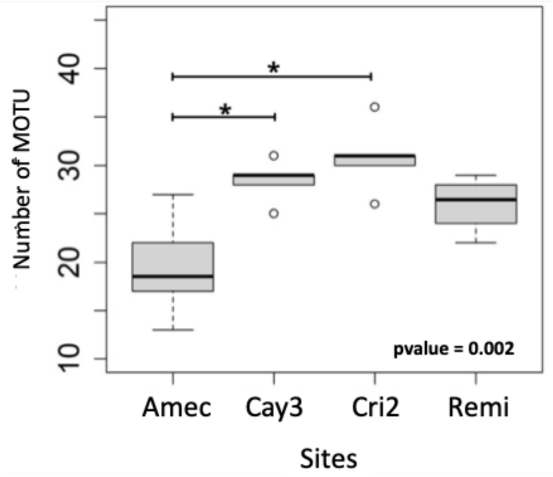

**Supplementary Figure S8.** Hierarchical cluster analysis (HCA) on log10(x+1) transformed core data. The Cay3_5-10cm value was removed since it was not representative. The distance used is Euclidean and the wardD2 jump was chosen. Site names (Amec, Remi, Cri2, Cay3, see Table 1) and sediment layers (1-2cm, 2-3cm, 3-4cm, 4-5cm, 5-10cm) are indicated.

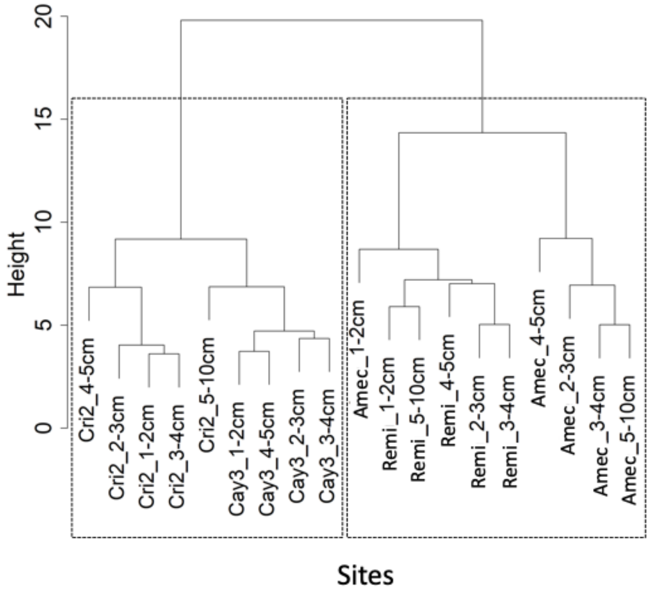

**Supplementary Figure S9.** Score (Areas under ROC curves) of the importance of the LDA variables. The Cay3_5-10cm value, not being representative, was excluded. The higher the score for a variable, the more important it is for the distinction between the two groups of observed sites (Ame/Rem and Cay3/Cri2).

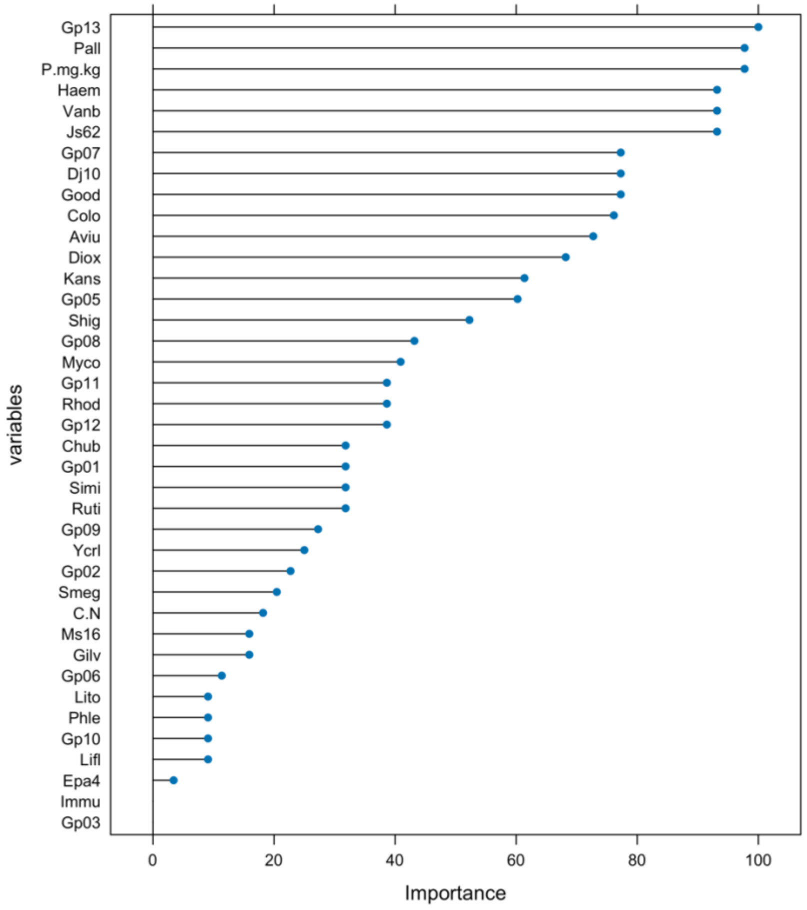

**Supplementary Figure S10.** Distribution of abiotic factors according to the type of site (U: urban or R: rural) collected from the water column in French Guiana (FG). The black bar indicates the median and the whiskers the extremum. The circle indicates an extreme value, i.e. greater than 1.5 times the interquartile range.

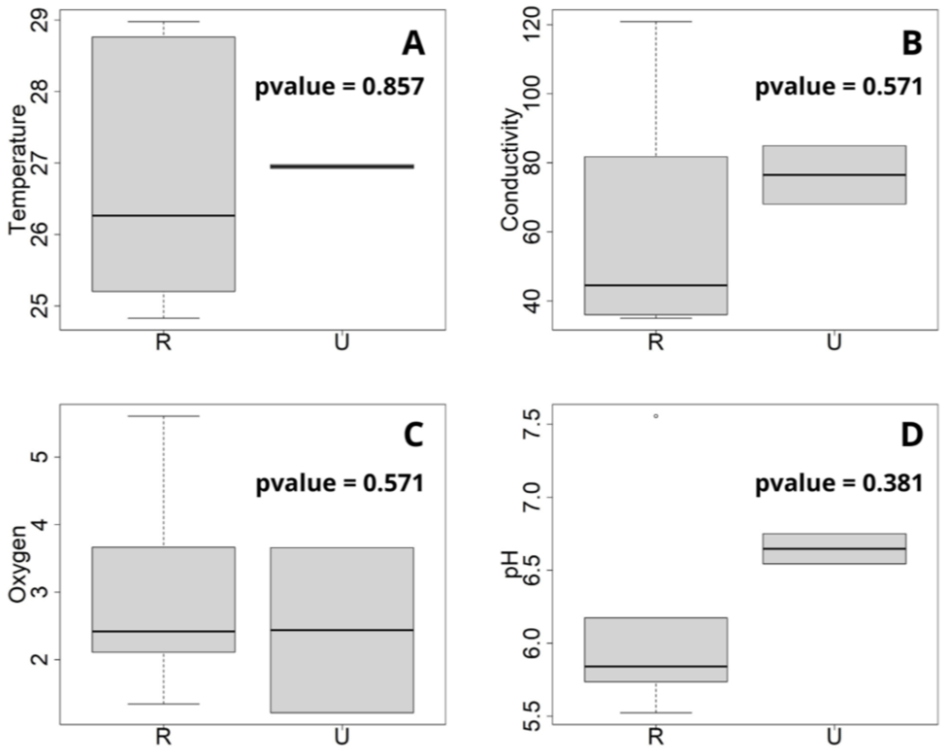

**Supplementary Figure S11.** Substrate distribution (%) for each site and for the sediment layers 0-5cm and 5-10cm. Cl.: Clay; C.Slt.: Coarse silts; C.Sd.: Coarse sands; F.Slt.: Fine silts and F.Sd.: Fine sands.

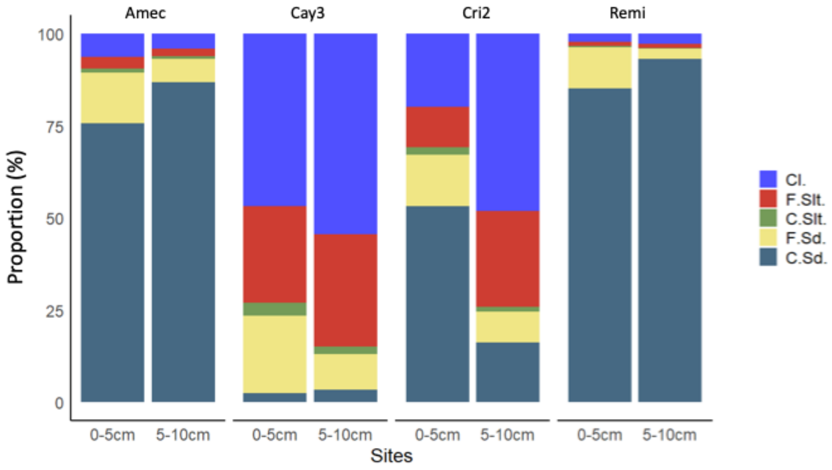

**Supplementary Figure S12.** Score (Air under ROC curve) of the contributions of each MOTU (panel A) and each substrate (panel B) in the RF analyses. The higher the score, the more important it is for the distinction between the two groups of observed sites (Amec/Remi and Cay3/Cri2).

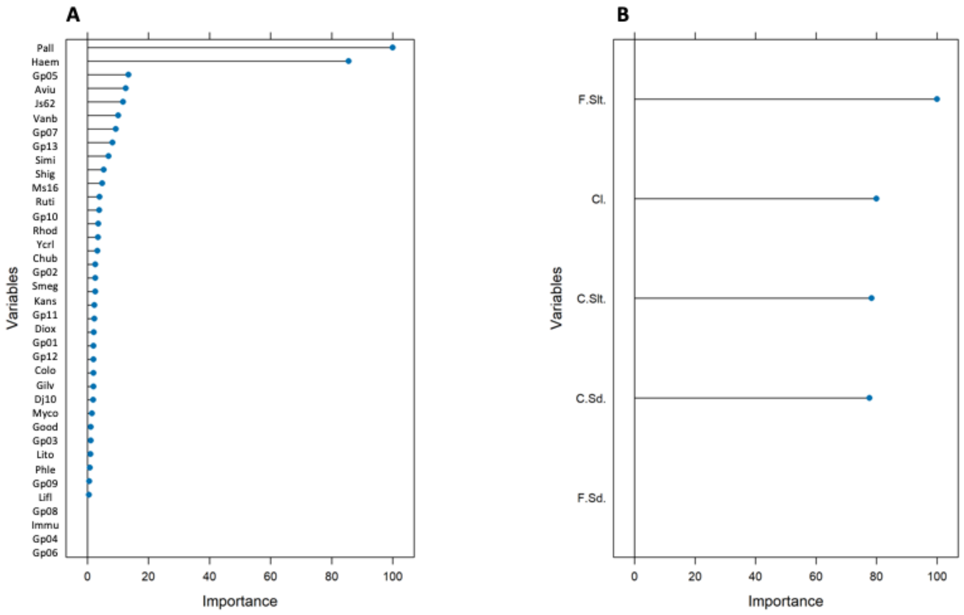

**Supplementary Figure S13.** Distribution of the MOTU Haem (panel A) and Pall (panel B) according to the 4 sites analyzed for sedimentary layers (0-5cm, 5-10cm). The black bar indicates the median and the whiskers indicate the extremum.

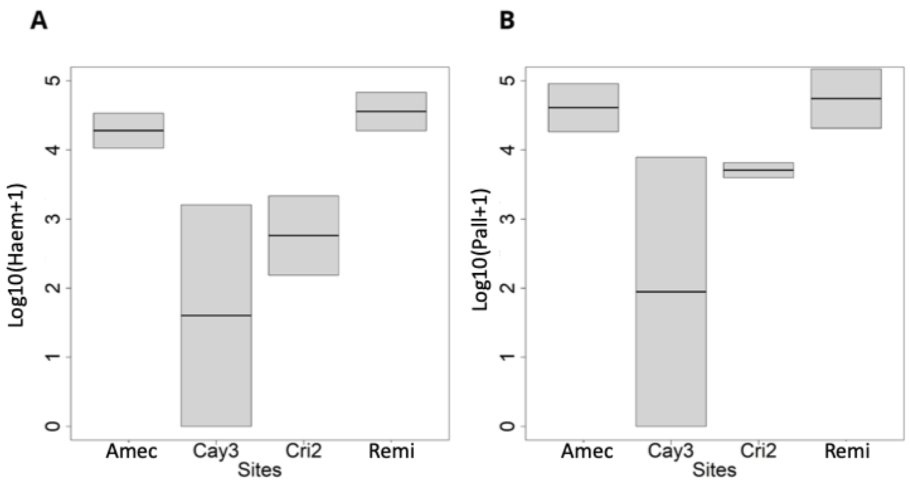

**Supplementary Figure S14.** Distribution of substrate variables (Cl.: Clay; F.Slt.: Fine silts; C.Slt: Coarse silts; F.Sd.: Fine sands; C.Sd.: Coarse sands) according to the 4 sites analyze for sediment layers (0-5cm, 5-10cm). The black bar indicates the median and the whiskers indicate the extremum.

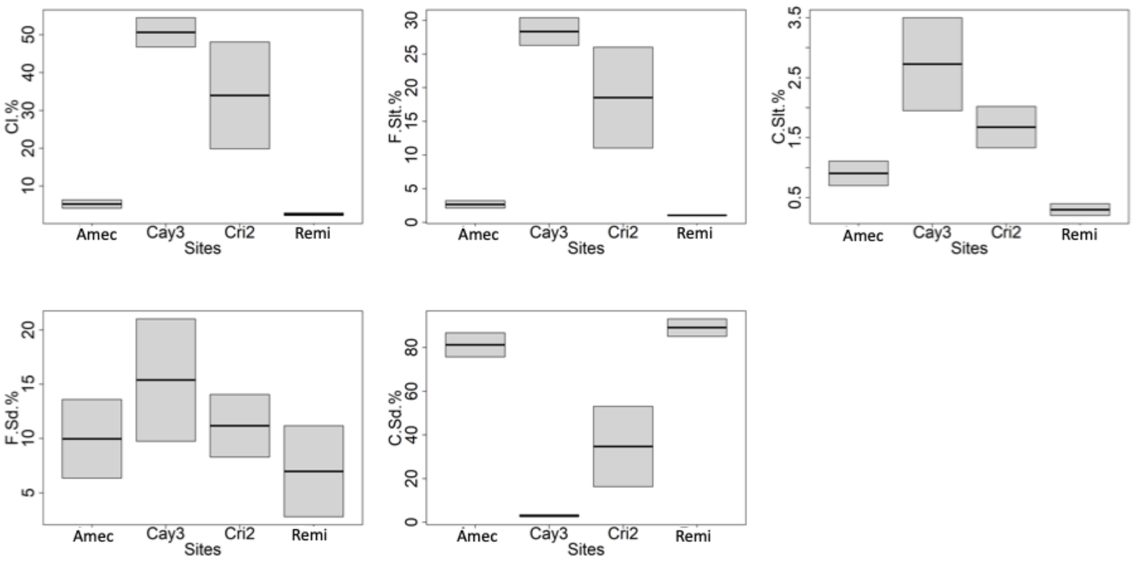

**Supplementary Figure S15.** Score (Air under ROC curve) of the contributions of each MOTU (panel A) and each substrate (panel B) in the RF analyses. The higher the score, the more important it is for the distinction between sediment layers (0-5cm and 5-10cm). The Cay3_5-10cm value was excluded as it was not representative.

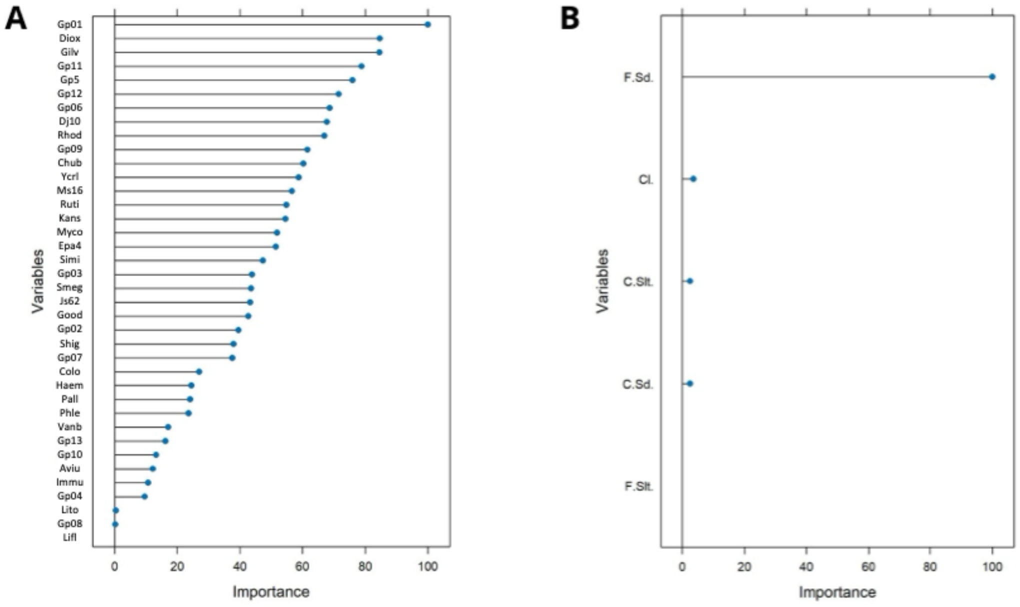

**Supplementary Figure S16.** Distribution of F.Sd. according to sediment layers (0-5cm, 5-10cm). The black bar indicates the median and the whiskers indicate the extremum.

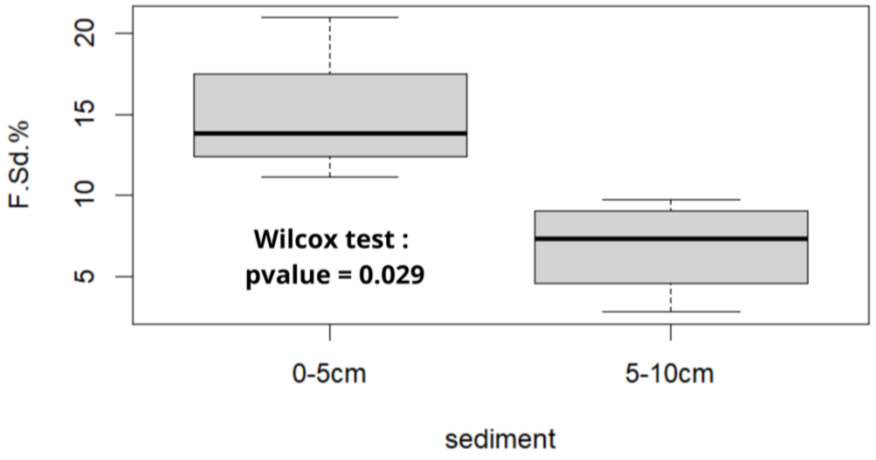

**Supplementary Figure S17.** Distribution of gp1, dio and gil according to sedimentary layers (0-5cm, 5-10cm). The black bar indicates the median and the whiskers represent the extremum.

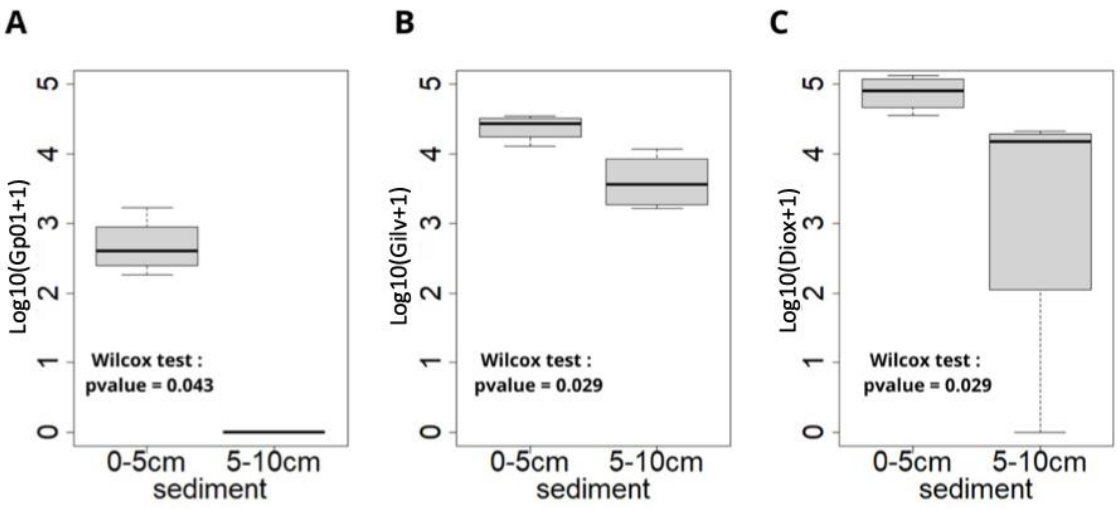

**Supplementary Figure S18.** Accumulation curve of the number of MOTUs in French Guiana (green) and Côte d'Ivoire (pink) across all sampled sites (12 sites for each country).

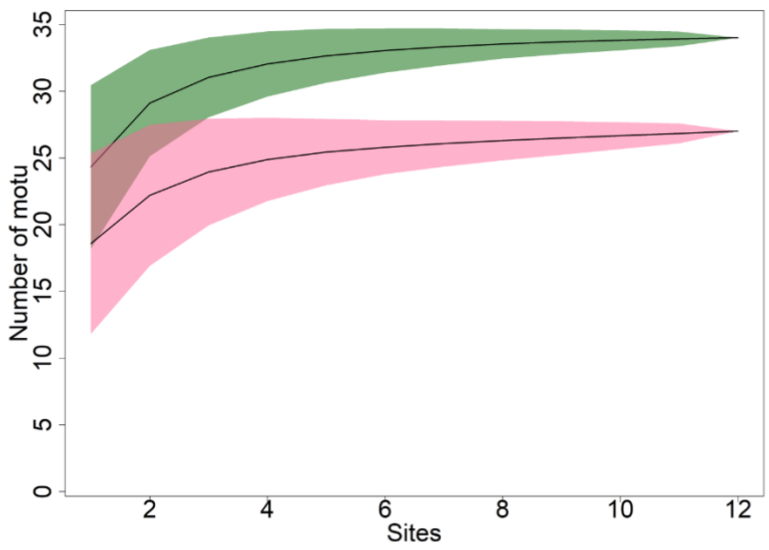

**Supplementary Figure S19.** MOTU prevalence (%) in FG (A) and CI (B).

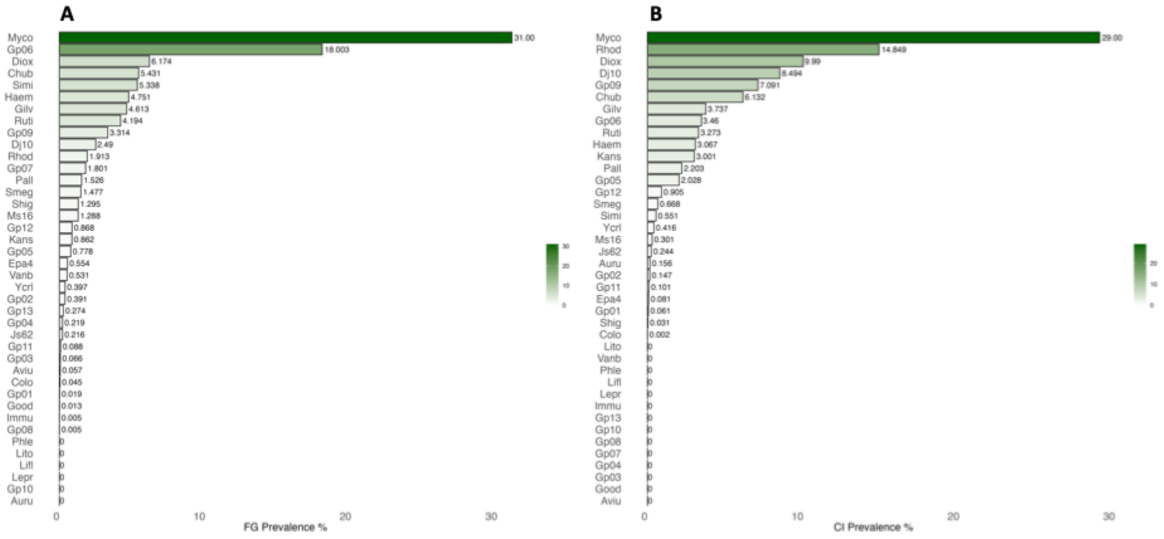

**Supplementary Figure S20.** Score (Area under ROC curves) of the importance of the RF variables of the FG and CI data. The higher the score for a MOTU, the more important it is for the distinction between the two groups of sites of the two countries observed (rural and urban).

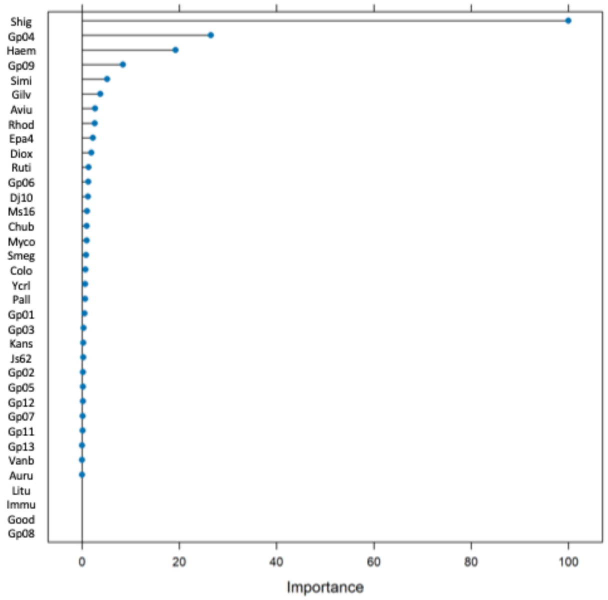

**Supplementary Table S1**. Abiotic parameters of the study sites in French Guiana (FG). W: Water; Temp: Temperature (°C); Cond: Conductivity; $O_2$: Oxygen; P%: Phosphorus (%); C%: Carbon (%); N%: Nitrogen (%); C/N: ration carbon/nitrogen; Cl.%: Clay (%); F.Slt.%: fine silt (%); C.Slt.%: coarse silt (%); F.Sd.%: fine sand (%); C.Sd.%: coarse sand (%). NA: unknown values.

| | | Water physico-chemical parameters | | | | Sediment physico-chemical parameters and granulometry | | | | | | | | |
|---|---|---|---|---|---|---|---|---|---|---|---|---|---|---|
| **Site** | **Habitat/ Layer** | **Temp.** | **Cond.** | **$O_2$** | **pH** | **P%** | **C%** | **N%** | **C/N** | **Cl.%** | **F.Slt.%** | **C.Slt.%** | **F.Sd.%** | **C.Sd.%** |
| **Tona** | W | 28.76275 | 81.81443 | 1.344060 | 5.840268 | NA | NA | NA | NA | NA | NA | NA | NA | NA |
| **Pse1** | W | 24.82847 | 36.06720 | 3.664556 | 5.736446 | NA | NA | NA | NA | NA | NA | NA | NA | NA |
| **Pse3** | W | 26.26319 | 35.02025 | 5.606503 | 5.523926 | NA | NA | NA | NA | NA | NA | NA | NA | NA |
| **Soul** | W | NA | NA | NA | NA | NA | NA | NA | NA | NA | NA | NA | NA | NA |
| **Jojo** | W | 28.97780 | 120.89666 | 2.417446 | 6.174224 | NA | NA | NA | NA | NA | NA | NA | NA | NA |
| **Mato** | W | 25.20238 | 44.52738 | 2.110714 | 7.555952 | NA | NA | NA | NA | NA | NA | NA | NA | NA |
| **Cri3** | W | NA | NA | NA | NA | NA | NA | NA | NA | NA | NA | NA | NA | NA |
| **Cri4** | W | NA | NA | NA | NA | NA | NA | NA | NA | NA | NA | NA | NA | NA |
| **Cay1** | W | NA | NA | NA | NA | NA | NA | NA | NA | NA | NA | NA | NA | NA |
| **Amec** | W | 26.97818 | 84.95678 | 1.212881 | 6.751059 | NA | NA | NA | NA | NA | NA | NA | NA | NA |
| | 1-2 cm | NA | NA | NA | NA | 296.3 | 0.05 | 0.78 | 15.82 | NA | NA | NA | NA | NA |
| | 2-3 cm | NA | NA | NA | NA | 341.8 | 0.08 | 1.55 | 19.76 | NA | NA | NA | NA | NA |
| | 3-4 cm | NA | NA | NA | NA | 358.3 | 0.09 | 1.71 | 19.10 | NA | NA | NA | NA | NA |
| | 4-5 cm | NA | NA | NA | NA | 460.2 | 0.15 | 2.70 | 17.55 | NA | NA | NA | NA | NA |
| | 0-5 cm | NA | NA | NA | NA | NA | NA | NA | NA | 6.34 | 3.22 | 1.11 | 13.60 | 75.73 |
| | 5-10 cm | NA | NA | NA | NA | 216.1 | 0.03 | 0.26 | 7.84 | 4.12 | 2.11 | 0.70 | 6.34 | 86.72 |
| **Remi** | W | 26.92329 | 68.08493 | 3.659041 | 6.543836 | NA | NA | NA | NA | NA | NA | NA | NA | NA |

| | | | | | | | | | | | | | | |
|---|---|---|---|---|---|---|---|---|---|---|---|---|---|---|
| | 1-2 cm | NA | NA | NA | NA | 260.2 | 0.03 | 0.15 | 5.51 | NA | NA | NA | NA | NA |
| | 2-3 cm | NA | NA | NA | NA | 313.9 | 0.03 | 0.19 | 6.43 | NA | NA | NA | NA | NA |
| | 3-4 cm | NA | NA | NA | NA | 314.3 | 0.02 | 0.17 | 6.96 | NA | NA | NA | NA | NA |
| | 4-5 cm | NA | NA | NA | NA | 230.7 | 0.03 | 0.15 | 4.48 | NA | NA | NA | NA | NA |
| | 0-5 cm | NA | NA | NA | NA | NA | NA | NA | NA | 2.19 | 1.1 | 0.4 | 11.17 | 85.14 |
| | 5-10 cm | NA | NA | NA | NA | 114.7 | 0.03 | 0.07 | 2.54 | 2.89 | 1.0 | 0.2 | 2.79 | 93.11 |
| **Cri2** | W | NA | NA | NA | NA | NA | NA | NA | NA | NA | NA | NA | NA | NA |
| | 1-2 cm | NA | NA | NA | NA | 547.1 | 0.05 | 0.72 | 13.24 | NA | NA | NA | NA | NA |
| | 2-3 cm | NA | NA | NA | NA | 571.9 | 0.04 | 0.58 | 13.81 | NA | NA | NA | NA | NA |
| | 3-4 cm | NA | NA | NA | NA | 607.6 | 0.06 | 0.70 | 12.18 | NA | NA | NA | NA | NA |
| | 4-5 cm | NA | NA | NA | NA | 515.0 | 0.07 | 0.94 | 13.29 | NA | NA | NA | NA | NA |
| | 0-5 cm | NA | NA | NA | NA | NA | NA | NA | NA | 19.82 | 11.02 | 2.02 | 14.05 | 53.08 |
| | 5-10 cm | NA | NA | NA | NA | 401.9 | 0.08 | 0.75 | 9.78 | 48.11 | 26.00 | 1.33 | 8.29 | 16.27 |
| **Cay3** | W | NA | NA | NA | NA | NA | NA | NA | NA | NA | NA | NA | NA | NA |
| | 1-2 cm | NA | NA | NA | NA | 769.5 | 0.12 | 1.30 | 11.06 | NA | NA | NA | NA | NA |
| | 2-3 cm | NA | NA | NA | NA | 857.8 | 0.12 | 1.29 | 10.56 | NA | NA | NA | NA | NA |
| | 3-4 cm | NA | NA | NA | NA | 868.0 | 0.14 | 1.86 | 13.58 | NA | NA | NA | NA | NA |
| | 4-5 cm | NA | NA | NA | NA | 819.4 | 0.12 | 1.45 | 12.08 | NA | NA | NA | NA | NA |
| | 0-5 cm | NA | NA | NA | NA | NA | NA | NA | NA | 46.76 | 26.26 | 3.50 | 21.01 | 2.47 |
| | 5-10 cm | NA | NA | NA | NA | 671.5 | 0.11 | 1.13 | 10.68 | 54.52 | 30.39 | 1.95 | 9.75 | 3.39 |

**Supplementary Table S2:** Invertebrate taxa collected in French Guiana (FG) and used to perform the analysis.

| Embranchement | Class | Order | Family | id |
|---|---|---|---|---|
| **ANNELIDA** | CLITELLATA | HAPLOTAXIDA | NA | 1 |
| **ANNELIDA** | CLITELLATA | RHYNCHOBDELLIDA | GLOSSIPHONIIDAE | 2 |
| **ARTHROPODA** | ARACHNIDA | ARANEAE | NA | 3 |
| **ARTHROPODA** | INSECTES | BLATTODEA | NA | 4 |
| **ARTHROPODA** | INSECTES | COLEOPTERA | CURCULIONOIDEA | 5 |
| **ARTHROPODA** | INSECTES | COLEOPTERA | DYTISCIDAE | 6 |
| **ARTHROPODA** | INSECTES | COLEOPTERA | ELMIDAE | 7 |
| **ARTHROPODA** | INSECTES | COLEOPTERA | HALIPLIDAE | 8 |
| **ARTHROPODA** | INSECTES | COLEOPTERA | HYDROCHIDAE | 9 |
| **ARTHROPODA** | INSECTES | COLEOPTERA | HYDROPHILIDAE | 10 |
| **ARTHROPODA** | INSECTES | COLEOPTERA | NA | 11 |
| **ARTHROPODA** | INSECTES | COLEOPTERA | NOTERIDAE | 12 |
| **ARTHROPODA** | INSECTES | COLEOPTERA | SCIRTIDAE | 13 |
| **ARTHROPODA** | INSECTES | COLEOPTERA | SIALIDAE | 14 |
| **ARTHROPODA** | INSECTES | COLEOPTERA | STAPHYLINIDAE | 15 |
| **ARTHROPODA** | INSECTES | DIPTERE | CERATOPOGONIDAE | 16 |
| **ARTHROPODA** | INSECTES | DIPTERE | CHIRONOMIDAE | 17 |
| **ARTHROPODA** | INSECTES | DIPTERE | CULICIDAE | 18 |
| **ARTHROPODA** | INSECTES | DIPTERE | EPHYDRIDAE | 19 |
| **ARTHROPODA** | INSECTES | DIPTERE | NA | 20 |
| **ARTHROPODA** | INSECTES | DIPTERE | TANYPONIDAE | 21 |

| | | | | |
|---|---|---|---|---|
| **ARTHROPODA** | INSECTES | EPHEMEROPTERA | BAETIDAE | 22 |
| **ARTHROPODA** | INSECTES | EPHEMEROPTERA | CAENIDAE | 23 |
| **ARTHROPODA** | INSECTES | EPHEMEROPTERA | EUTHYPLOCIIDAE | 24 |
| **ARTHROPODA** | INSECTES | EPHEMEROPTERA | LEPTOHYPHIDAE | 25 |
| **ARTHROPODA** | INSECTES | EPHEMEROPTERA | LEPTOPHLEBIIDAE | 26 |
| **ARTHROPODA** | INSECTES | EPHEMEROPTERA | NA | 27 |
| **ARTHROPODA** | INSECTES | EPHEMEROPTERA | POLYMITARCIDAE | 28 |
| **ARTHROPODA** | INSECTES | EPHEMEROPTERA | POTAMANTHIDAE | 29 |
| **ARTHROPODA** | INSECTES | HEMIPTERA | BELOSTOMATIDAE | 30 |
| **ARTHROPODA** | INSECTES | HEMIPTERA | CORIXIDAE | 31 |
| **ARTHROPODA** | INSECTES | HEMIPTERA | GERRIDAE | 32 |
| **ARTHROPODA** | INSECTES | HEMIPTERA | HELOTREPHIDAE | 33 |
| **ARTHROPODA** | INSECTES | HEMIPTERA | HYDROMETRIDAE | 34 |
| **ARTHROPODA** | INSECTES | HEMIPTERA | MESOVELIIDAE | 35 |
| **ARTHROPODA** | INSECTES | HEMIPTERA | NAUCORIDAE | 36 |
| **ARTHROPODA** | INSECTES | HEMIPTERA | NEPIDAE | 37 |
| **ARTHROPODA** | INSECTES | HEMIPTERA | NOTONECTIDAE | 38 |
| **ARTHROPODA** | INSECTES | HEMIPTERA | VELIIDAE | 39 |
| **ARTHROPODA** | INSECTES | HYMENOPTERA | FORMICOIDAE | 40 |
| **ARTHROPODA** | INSECTES | LEPIDOPTERA | EREBIDAE | 41 |
| **ARTHROPODA** | INSECTES | LEPIDOPTERA | NA | 42 |
| **ARTHROPODA** | INSECTES | ODONATA | AESHNIDAE | 43 |
| **ARTHROPODA** | INSECTES | ODONATA | COENAGRIONIDAE | 44 |
| **ARTHROPODA** | INSECTES | ODONATA | CORDULIIDAE | 45 |
| **ARTHROPODA** | INSECTES | ODONATA | DICTERIADIDAE | 46 |
| **ARTHROPODA** | INSECTES | ODONATA | GOMPHIDAE | 47 |
| **ARTHROPODA** | INSECTES | ODONATA | LIBELLULIDAE | 48 |

| | | | | |
|---|---|---|---|---|
| **ARTHROPODA** | INSECTES | PLECOPTERA | PERLIDAE | 49 |
| **ARTHROPODA** | INSECTES | TRICHOPTERA | ANOMALOPSYCHIDAE | 50 |
| **ARTHROPODA** | INSECTES | TRICHOPTERA | HYDROPSYCHIDAE | 51 |
| **ARTHROPODA** | INSECTES | TRICHOPTERA | HYDROPTILIDAE | 52 |
| **ARTHROPODA** | INSECTES | TRICHOPTERA | LEPTOCERIDAE | 53 |
| **ARTHROPODA** | INSECTES | TRICHOPTERA | LIMNEPHILIDAE | 54 |
| **ARTHROPODA** | INSECTES | TRICHOPTERA | POLYCENTROPODIDAE | 55 |
| **ARTHROPODA** | MALACOSTRACA | DECAPODA | PALAEMONIDAE | 56 |
| **ARTHROPODA** | MALACOSTRACA | DECAPODA | PARASTACIDAE | 57 |
| **ARTHROPODA** | MALACOSTRACA | DECAPODA | TRICHODACTYLIDAE | 58 |
| **CHORDATA** | AMPHIBIA | ANURA | NA | 59 |
| **MOLLUSCA** | BIVALVE | VENEROIDA | SPHAERIIDAE | 60 |
| **MOLLUSCA** | GASTROPODA | ARCHITAENIOGLOSSA | NA | 61 |
| **MOLLUSCA** | GASTROPODA | BASOMMOTOPHORA | NA | 62 |
| **MOLLUSCA** | GASTROPODA | BASOMMOTOPHORA | PHYSIDAE | 63 |
| **MOLLUSCA** | GASTROPODA | BASOMMOTOPHORA | PLANORBIDAE | 64 |
| **MOLLUSCA** | GASTROPODA | LITTORINIMORPHA | LITTORINIDAE | 65 |
| **MOLLUSCA** | GASTROPODA | LITTORINIMORPHA | NA | 66 |
| **MOLLUSCA** | GASTROPODA | STYLOMMATOPHORA | NA | 67 |

**Supplementary Table S3**. Mycobacterial DNA used as positive control for the 16S rDNA PCR amplification and Illumina sequencing.

| **Species** | **ng/µL** | **A260/A280** |
|---|---|---|
| *M. ulcerans* | 97.9 | 1.299 |
| *M. marinum* | -2.3 | 1.744 |
| *M. fortuitum* | -1.7 | 1.413 |
| *M. avium* | 34.6 | 1.564 |
| *M. chelonae* | -1.1 | 1.853 |
| *M. absessus* | 39.8 | 2.174 |
| *M. smegmatis* | 64.3 | 1.724 |
| *M. lepreae* | 0.7 | 3.643 |
| *M. lepromatosis* | 49.9 | 1.821 |

**Supplementary Table S4**. List of mycobacterial species used in this study and accession Numbers where the 16S sequences were recovered.

| Accession Number | Species | Accession Number | Species | Accession Number | Species |
|---|---|---|---|---|---|
| NC_002944 | Mycobacterium avium | NC_009565 | Mycobacterium tuberculosis | NC_015576 | Mycobacterium sinense |
| NC_002755 | Mycobacterium tuberculosis | NC_010612 | Mycobacterium marinum | NC_015576 | Mycobacterium sinense |
| NC_008769 | Mycobacterium bovis | NC_012943 | Mycobacterium tuberculosis | NZ_CM001225 | Mycobacterium tuberculosis |
| NC_012207 | Mycobacterium bovis | NC_020133 | Mycobacterium liflandii | NZ_CM001226 | Mycobacterium tuberculosis |
| NC_008611 | Mycobacterium ulcerans | NC_010397 | Mycobacterium abscessus | NZ_CM001227 | Mycobacterium tuberculosis |
| NC_008146 | Mycobacterium sp. MCS | NC_022350 | Mycobacterium tuberculosis | NC_017524 | Mycobacterium tuberculosis |
| NC_008146 | Mycobacterium sp. MCS | NC_016768 | Mycobacterium tuberculosis | NC_016604 | Mycobacterium rhodesiae |
| NC_008595 | Mycobacterium avium | NC_018078 | Mycobacterium tuberculosis | NC_016604 | Mycobacterium rhodesiae |
| NC_008596 | Mycobacterium smegmatis | NC_022663 | Mycobacterium kansasii | NC_016804 | Mycobacterium bovis |
| NC_008596 | Mycobacterium smegmatis | NZ_CM000787 | Mycobacterium tuberculosis | NC_015758 | Mycobacterium africanum |
| NC_008726 | Mycobacterium vanbaalenii | NZ_CM000788 | Mycobacterium tuberculosis | NC_015848 | Mycobacterium canettii |
| NC_008726 | Mycobacterium vanbaalenii | NZ_CM000789 | Mycobacterium tuberculosis | NC_017904 | Mycobacterium sp. MOTT36Y |
| NC_008705 | Mycobacterium sp. KMS | NZ_CM001043 | Mycobacterium tuberculosis | NZ_CM001515 | Mycobacterium tuberculosis |
| NC_008705 | Mycobacterium sp. KMS | NZ_CM001044 | Mycobacterium tuberculosis | NC_018027 | Mycobacterium chubuense |
| NC_009077 | Mycobacterium sp. JLS | NZ_CM001045 | Mycobacterium tuberculosis | NC_018027 | Mycobacterium chubuense |
| NC_009077 | Mycobacterium sp. JLS | NC_014814 | Mycobacterium gilvum | NC_017523 | Mycobacterium tuberculosis |
| NC_009525 | Mycobacterium tuberculosis | NC_014814 | Mycobacterium gilvum | NC_017522 | Mycobacterium tuberculosis |
| NC_009338 | Mycobacterium gilvum | NZ_CP012090 | Mycobacterium tuberculosis | NC_016948 | Mycobacterium paraintracellular |
| NC_009338 | Mycobacterium gilvum | NZ_CP003494 | Mycobacterium bovis | NC_016946 | Mycobacterium intracellulare |

| Accession Number | Species | Accession Number | Species | Accession Number | Species |
|---|---|---|---|---|---|
| NC_018143 | Mycobacterium tuberculosis | NC_000962 | Mycobacterium tuberculosis | NC_016947 | Mycobacterium intracellulare |
| NC_018150 | Mycobacterium abscessus | NC_021054 | Mycobacterium tuberculosis | NZ_CM002056 | Mycobacterium tuberculosis |
| NC_018289 | Mycobacterium smegmatis | NC_021194 | Mycobacterium tuberculosis | NZ_CM002058 | Mycobacterium tuberculosis |
| NC_018289 | Mycobacterium smegmatis | NC_021200 | Mycobacterium avium | NZ_CM002059 | Mycobacterium tuberculosis |
| NC_018612 | Mycobacterium indicus pranii | NC_021251 | Mycobacterium tuberculosis | NZ_CM002060 | Mycobacterium tuberculosis |
| NC_016934 | Mycobacterium tuberculosis | NZ_CP009914 | Mycobacterium sp. VKM Ac-1817D | NZ_CM002061 | Mycobacterium tuberculosis |
| NC_023036 | Mycobacterium neoaurum | NZ_CP009914 | Mycobacterium sp. VKM Ac-1817D | NZ_CM002062 | Mycobacterium tuberculosis |

| | | | | | |
|---|---|---|---|---|---|
| NC_023036 | Mycobacterium neoaurum | NZ_CM002022 | Mycobacterium tuberculosis | NZ_CM002063 | Mycobacterium tuberculosis |
| NC_019966 | Mycobacterium sp. JS623 | NC_021715 | Mycobacterium yongonense | NZ_CM002064 | Mycobacterium tuberculosis |
| NC_019966 | Mycobacterium sp. JS623 | NC_021740 | Mycobacterium tuberculosis | NZ_CM002065 | Mycobacterium tuberculosis |
| NC_019951 | Mycobacterium canettii | NZ_CM002048 | Mycobacterium tuberculosis | NZ_CM002066 | Mycobacterium tuberculosis |
| NC_019950 | Mycobacterium canettii | NZ_CM002049 | Mycobacterium tuberculosis | NZ_CM002067 | Mycobacterium tuberculosis |
| NC_019965 | Mycobacterium canettii | NZ_CM002050 | Mycobacterium tuberculosis | NZ_CM002068 | Mycobacterium tuberculosis |
| NZ_CM001762 | Mycobacterium smegmatis | NZ_CM002051 | Mycobacterium tuberculosis | NZ_CM002069 | Mycobacterium tuberculosis |
| NZ_CM001762 | Mycobacterium smegmatis | NZ_CM002052 | Mycobacterium tuberculosis | NZ_CM002070 | Mycobacterium tuberculosis |
| NC_020089 | Mycobacterium tuberculosis | NZ_CM002053 | Mycobacterium tuberculosis | NZ_CM002073 | Mycobacterium tuberculosis |
| NC_020245 | Mycobacterium bovis | NZ_CM002054 | Mycobacterium tuberculosis | NZ_CM002071 | Mycobacterium tuberculosis |
| NZ_CP011883 | Mycobacterium haemophilum | NZ_CM002055 | Mycobacterium tuberculosis | NZ_CM002072 | Mycobacterium tuberculosis |
| NC_020559 | Mycobacterium tuberculosis | NZ_CM002057 | Mycobacterium tuberculosis | NZ_CM002076 | Mycobacterium tuberculosis |

| **Accession Number** | **Species** | **Accession Number** | **Species** | **Accession Number** | **Species** |
|---|---|---|---|---|---|
| NZ_CM002077 | Mycobacterium tuberculosis | NZ_CM002115 | Mycobacterium tuberculosis | NZ_AP014547 | Mycobacterium abscessus |
| NZ_CM002079 | Mycobacterium tuberculosis | NZ_CM002117 | Mycobacterium tuberculosis | NZ_CP002871 | Mycobacterium tuberculosis |
| NZ_CM002074 | Mycobacterium tuberculosis | NZ_CM002113 | Mycobacterium tuberculosis | NZ_CP002882 | Mycobacterium tuberculosis |
| NZ_CM002075 | Mycobacterium tuberculosis | NZ_CM002111 | Mycobacterium tuberculosis | NZ_CP002883 | Mycobacterium tuberculosis |
| NZ_CM002078 | Mycobacterium tuberculosis | NZ_CM002109 | Mycobacterium tuberculosis | NZ_CP002885 | Mycobacterium tuberculosis |
| NZ_CM002080 | Mycobacterium tuberculosis | NZ_CM002112 | Mycobacterium tuberculosis | NZ_CP007803 | Mycobacterium tuberculosis |
| NC_021282 | Mycobacterium abscessus | NZ_CM002108 | Mycobacterium tuberculosis | NZ_CP007809 | Mycobacterium tuberculosis |
| NZ_CM002127 | Mycobacterium tuberculosis | NZ_CM002110 | Mycobacterium tuberculosis | NZ_HG917972 | Mycobacterium marinum |
| NZ_CM002126 | Mycobacterium tuberculosis | NZ_CM002107 | Mycobacterium tuberculosis | NZ_HG917972 | Mycobacterium marinum |
| NZ_CM002125 | Mycobacterium tuberculosis | NZ_CM002106 | Mycobacterium tuberculosis | NZ_CP009100 | Mycobacterium tuberculosis |
| NZ_CM002124 | Mycobacterium tuberculosis | NZ_CM002105 | Mycobacterium tuberculosis | NZ_CP009101 | Mycobacterium tuberculosis |
| NZ_CM002122 | Mycobacterium tuberculosis | NZ_CM002104 | Mycobacterium tuberculosis | NZ_CM002882 | Mycobacterium tuberculosis |
| NZ_CM002121 | Mycobacterium tuberculosis | NZ_CM002102 | Mycobacterium tuberculosis | NZ_CM002883 | Mycobacterium tuberculosis |
| NZ_CM002120 | Mycobacterium tuberculosis | NZ_CM002103 | Mycobacterium tuberculosis | NZ_CM002884 | Mycobacterium tuberculosis |
| NZ_CM002123 | Mycobacterium tuberculosis | NZ_CM002101 | Mycobacterium tuberculosis | NZ_CP009426 | Mycobacterium tuberculosis |
| NZ_CM002119 | Mycobacterium tuberculosis | NZ_CM002100 | Mycobacterium tuberculosis | NZ_CP009427 | Mycobacterium tuberculosis |
| NZ_CM002114 | Mycobacterium tuberculosis | NZ_CM002098 | Mycobacterium tuberculosis | NZ_CP009447 | Mycobacterium abscessus |
| NZ_CM002116 | Mycobacterium tuberculosis | NZ_CM002099 | Mycobacterium tuberculosis | NZ_CP009449 | Mycobacterium bovis |
| NZ_CM002118 | Mycobacterium tuberculosis | NZ_CM002097 | Mycobacterium tuberculosis | NZ_CP009493 | Mycobacterium avium |

| Accession Number | Species | Accession Number | Species | Accession Number | Species |
|---|---|---|---|---|---|
| NZ_CP009407 | Mycobacterium abscessus | NZ_CP009483 | Mycobacterium kansasii | NZ_CP011269 | Mycobacterium fortuitum |
| NZ_CP009408 | Mycobacterium abscessus | NZ_CP007299 | Mycobacterium tuberculosis | NZ_CP011269 | Mycobacterium fortuitum |
| NZ_CP009499 | Mycobacterium intracellulare | NZ_CP010113 | Mycobacterium avium | NZ_CP013049 | Mycobacterium abscessus |
| NZ_CP009494 | Mycobacterium smegmatis | NZ_CP010114 | Mycobacterium avium | NZ_CP010271 | Mycobacterium saopaulense |
| NZ_CP009494 | Mycobacterium smegmatis | NZ_CP015773 | Mycobacterium bovis | NZ_LN831039 | Mycobacterium smegmatis |
| NZ_CP009495 | Mycobacterium smegmatis | NZ_CP010895 | Mycobacterium tuberculosis | NZ_LN831039 | Mycobacterium smegmatis |
| NZ_CP009495 | Mycobacterium smegmatis | NZ_CP010873 | Mycobacterium tuberculosis | NZ_CP013741 | Mycobacterium bovis |
| NZ_CP009496 | Mycobacterium smegmatis | NZ_AM412059 | Mycobacterium bovis | NZ_CP010330 | Mycobacterium tuberculosis |
| NZ_CP009496 | Mycobacterium smegmatis | NZ_CP010946 | Mycobacterium chelonae | NZ_CP010337 | Mycobacterium tuberculosis |
| NZ_CP009613 | Mycobacterium abscessus | NZ_CP011773 | Mycobacterium sp. EPa45 | NZ_CP010338 | Mycobacterium tuberculosis |
| NZ_CP009615 | Mycobacterium abscessus | NZ_CP011773 | Mycobacterium sp. EPa45 | NZ_CP010339 | Mycobacterium tuberculosis |
| NZ_CP009616 | Mycobacterium abscessus | NZ_CP008744 | Mycobacterium bovis | NZ_CP010340 | Mycobacterium tuberculosis |
| NZ_CP009614 | Mycobacterium avium | NZ_CP012044 | Mycobacterium abscessus | NZ_CP014566 | Mycobacterium bovis |
| NZ_HG813240 | Mycobacterium tuberculosis | NZ_CP012095 | Mycobacterium bovis | NZ_CP011022 | Mycobacterium sp. NRRL B-3805 |
| NZ_CP007027 | Mycobacterium tuberculosis | NZ_CP012150 | Mycobacterium goodii | NZ_CP011022 | Mycobacterium sp. NRRL B-3805 |
| NZ_AP014573 | Mycobacterium tuberculosis | NZ_CP012150 | Mycobacterium goodii | NZ_CP014475 | Mycobacterium phlei |
| NZ_AP012555 | Mycobacterium avium | NZ_CP009243 | Mycobacterium bovis | NZ_CP014475 | Mycobacterium phlei |
| NZ_CP009480 | Mycobacterium tuberculosis | NZ_CP012506 | Mycobacterium tuberculosis | NZ_CP010968 | Mycobacterium tuberculosis |
| NZ_CP009482 | Mycobacterium avium | NZ_CP011455 | Mycobacterium bovis | NZ_CP010996 | Mycobacterium simiae |

| Accession Number | Species | Accession Number | Species | Accession Number | Species |
|---|---|---|---|---|---|
| NZ_CP014617 | Mycobacterium africanum | NZ_CP015596 | Mycobacterium sp. YC-RL4 | NZ_CP013475 | Mycobacterium tuberculosis |
| NZ_CP011530 | Mycobacterium immunogenum | NZ_CP015495 | Mycobacterium avium | NZ_CP017920 | Mycobacterium tuberculosis |
| NZ_CP011530 | Mycobacterium immunogenum | NZ_CP011491 | Mycobacterium vaccae | NZ_CP018043 | Mycobacterium sp. WY10 |
| NZ_CP014950 | Mycobacterium abscessus | NZ_CP011491 | Mycobacterium vaccae | NZ_CP018043 | Mycobacterium sp. WY10 |
| NZ_CP014951 | Mycobacterium abscessus | NZ_CP016188 | Mycobacterium abscessus | NZ_CP018303 | Mycobacterium tuberculosis |
| NZ_CP014952 | Mycobacterium abscessus | NZ_CP016189 | Mycobacterium immunogenum | NZ_CP018305 | Mycobacterium tuberculosis |
| NZ_CP014953 | Mycobacterium abscessus | NZ_CP016189 | Mycobacterium immunogenum | NZ_CP018302 | Mycobacterium tuberculosis |
| NZ_CP014954 | Mycobacterium abscessus | NZ_CP016190 | Mycobacterium abscessus | NZ_CP018301 | Mycobacterium tuberculosis |
| NZ_CP014955 | Mycobacterium abscessus | NZ_CP016191 | Mycobacterium abscessus | NZ_CP018300 | Mycobacterium tuberculosis |

| | | | | | |
|---|---|---|---|---|---|
| NZ_CP014956 | Mycobacterium abscessus | NZ_CP016192 | Mycobacterium abscessus | NZ_CP018304 | Mycobacterium tuberculosis |
| NZ_CP014957 | Mycobacterium abscessus | NZ_CP016193 | Mycobacterium abscessus | NZ_CP018778 | Mycobacterium tuberculosis |
| NZ_CP014958 | Mycobacterium abscessus | NZ_CP016396 | Mycobacterium avium | NZ_CP018363 | Mycobacterium avium |
| NZ_CP014959 | Mycobacterium abscessus | NZ_CP016640 | Mycobacterium sp. djl-10 | NZ_CP016972 | Mycobacterium tuberculosis |
| NZ_CP014960 | Mycobacterium abscessus | NZ_CP016640 | Mycobacterium sp. djl-10 | NZ_CP016401 | Mycobacterium caprae |
| NZ_CP014961 | Mycobacterium abscessus | NZ_CP016794 | Mycobacterium tuberculosis | NZ_CM007645 | Mycobacterium tuberculosis |
| NZ_CP010071 | Mycobacterium sp. QIA-37 | NZ_CP016888 | Mycobacterium tuberculosis | NZ_CM007646 | Mycobacterium tuberculosis |
| NZ_CP007220 | Mycobacterium chelonae | NZ_CP012885 | Mycobacterium chimaera | NZ_CP019420 | Mycobacterium sp. MS1601 |
| NZ_CP015596 | Mycobacterium sp. YC-RL4 | NZ_CP011510 | Mycobacterium tuberculosis | NZ_CP019420 | Mycobacterium sp. MS1601 |

| **Accession Number** | **Species** | **Accession Number** | **Species** | **Accession Number** | **Species** |
|---|---|---|---|---|---|
| NZ_CP008702 | Mycobacterium tuberculosis | NZ_CP009180 | Mycobacterium tuberculosis | NZ_CP009195 | Mycobacterium tuberculosis |
| NZ_CP009198 | Mycobacterium tuberculosis | NZ_CP009181 | Mycobacterium tuberculosis | NZ_CP009187 | Mycobacterium tuberculosis |
| NZ_CP009199 | Mycobacterium tuberculosis | NZ_CP009182 | Mycobacterium tuberculosis | NZ_CP009183 | Mycobacterium tuberculosis |
| NZ_CP009200 | Mycobacterium tuberculosis | NZ_CP009184 | Mycobacterium tuberculosis | NZ_CP019882 | Mycobacterium litorale |
| NZ_CP009201 | Mycobacterium tuberculosis | NZ_CP009185 | Mycobacterium tuberculosis | NZ_CP019882 | Mycobacterium litorale |
| NZ_CP009202 | Mycobacterium tuberculosis | NZ_CP009186 | Mycobacterium tuberculosis | NZ_CP020381 | Mycobacterium tuberculosis |
| NZ_CP009203 | Mycobacterium tuberculosis | NZ_CP009188 | Mycobacterium tuberculosis | NZ_CP019883 | Mycobacterium kansasii |
| NZ_CP009204 | Mycobacterium tuberculosis | NZ_CP009189 | Mycobacterium tuberculosis | NZ_CP019884 | Mycobacterium kansasii |
| NZ_CP009205 | Mycobacterium tuberculosis | NZ_CP009194 | Mycobacterium tuberculosis | NZ_CP019885 | Mycobacterium kansasii |
| NZ_CP009206 | Mycobacterium tuberculosis | NZ_CP009196 | Mycobacterium tuberculosis | NZ_CP019886 | Mycobacterium kansasii |
| NZ_CP009207 | Mycobacterium tuberculosis | NZ_CP009197 | Mycobacterium tuberculosis | NZ_CP019887 | Mycobacterium kansasii |
| NZ_CP009172 | Mycobacterium tuberculosis | NZ_CP009190 | Mycobacterium tuberculosis | NZ_CP019888 | Mycobacterium kansasii |
| NZ_CP009173 | Mycobacterium tuberculosis | NZ_CP009191 | Mycobacterium tuberculosis | NZ_CP020821 | Mycobacterium colombiense |
| NZ_CP009174 | Mycobacterium tuberculosis | NZ_CP009193 | Mycobacterium tuberculosis | NZ_CP017593 | Mycobacterium tuberculosis |
| NZ_CP009175 | Mycobacterium tuberculosis | NZ_CP009192 | Mycobacterium tuberculosis | NZ_CP017594 | Mycobacterium tuberculosis |
| NZ_CP009176 | Mycobacterium tuberculosis | NZ_CP017596 | Mycobacterium tuberculosis | NZ_CP017595 | Mycobacterium tuberculosis |
| NZ_CP009177 | Mycobacterium tuberculosis | NZ_CP017597 | Mycobacterium tuberculosis | NZ_CP015964 | Mycobacterium yongonense |
| NZ_CP009178 | Mycobacterium tuberculosis | NZ_CP017598 | Mycobacterium tuberculosis | NZ_CP015965 | Mycobacterium yongonense |
| NZ_CP009179 | Mycobacterium tuberculosis | NZ_CP021122 | Mycobacterium abscessus | NZ_CP020809 | Mycobacterium dioxanotrophicus |

| **Accession Number** | **Species** | **Accession Number** | **Species** |
|---|---|---|---|

| | | | |
|---|---|---|---|
| NZ_CP020809 | Mycobacterium dioxanotrophicus | NZ_AP017901 | Mycobacterium tuberculosis |
| NZ_CP020809 | Mycobacterium dioxanotrophicus | NZ_AP018033 | Mycobacterium tuberculosis |
| NZ_CP019221 | Mycobacterium chimaera | NZ_AP018164 | Mycobacterium shigaense |
| NZ_CP022014 | Mycobacterium tuberculosis | NZ_AP018165 | Mycobacterium stephanolepidis |
| NZ_CP022095 | Mycobacterium avium | NZ_AP017635 | Mycobacterium ulcerans |
| NZ_CP022105 | Mycobacterium avium | NZ_AP018034 | Mycobacterium tuberculosis |
| NZ_CP015267 | Mycobacterium chimaera | NZ_AP018035 | Mycobacterium tuberculosis |
| NZ_CP015272 | Mycobacterium chimaera | NZ_AP018036 | Mycobacterium tuberculosis |
| NZ_CP015278 | Mycobacterium chimaera | NZ_LT549889 | Mycobacterium aurum |
| NZ_CP022223 | Mycobacterium chimaera | NZ_LT549889 | Mycobacterium aurum |
| NZ_CP023149 | Mycobacterium intracellulare | NZ_LT629971 | Mycobacterium rutilum |
| NZ_CP023146 | Mycobacterium intracellulare | NZ_LT629971 | Mycobacterium rutilum |
| NZ_CP023147 | Mycobacterium marseillense | NZ_LT703505 | Mycobacterium chimaera |
| NZ_CP023151 | Mycobacterium chimaera | NZ_LT906469 | Mycobacterium terrae |
| NZ_CP023169 | Mycobacterium tuberculosis | NZ_LT906469 | Mycobacterium terrae |
| NZ_CP023170 | Mycobacterium tuberculosis | | |
| NZ_CP023435 | Mycobacterium pallens | | |
| NZ_CP023435 | Mycobacterium pallens | | |
| NZ_AP017624 | Mycobacterium ulcerans | | |

**Supplementary Table S5**. Known pathogenic environmental mycobacteria with confirmed or suspected waterborne transmission. MAC; *Mycobacterium avium* Complex which include *M. intracellulare*, *M. avium* and its subspecies *avium* (MAA), *paratuberculosis* (MAP) and *sylvaticum*. Non-exhaustive list.

| Organism | Hosts | Diseases | Reservoir/Source of infection | References |
|---|---|---|---|---|
| MAC | Humans | Pulmonary disease (elderly and/or immunocompromised patients) | Hot tub | Mangione et al., 2001; Embil et al., 1997 |
| | Humans | Cutaneous infection | Circulating bath water | Sugita et al., 2000 |
| | Humans | Hypersensitivity pneumonitis | Hot tub | Rickman et al., 2002 |
| | Humans | Cervical lymphadenitis | Drinking stagnant Water | Oloya et al., 2008 |
| *M. avium* | Birds, chickens, pigeons, emus, rheas | Avian mycobacteriosis. Lesions in the liver and gastrointestinal tract | Water contaminated with animal feces | Mycobacteria book |
| | Dogs, cats, armadillos, macaques, marsupial, water buffalo, cattle, pigs, deer, horses | Tuberculosis-like disease | Water contaminated with animal feces | Mycobacteria book |
| *M. bovis* | Humans | Cervical lymphadenitis | Drinking stagnant Water | Oloya et al., 2008 |
| | Cattle, swine, sheep, goats, horses, water buffalo, camels, deer, cats, dogs, etc. | Tuberculosis-like disease | Contaminated water with urine and feces (experimental results) | Cosivi et al., 1995 |
| *M. fortuitum* | Humans | Furunculosis | Whirlpool footbaths at a nail salon | Winthrop et al., 2002 |
| | Humans | Respiratory tract colonization | Hospital ice machine | Labombardi et al., 2002 ; Gebo et al., 2002 |
| | Humans | | Natural waters, Sewage | Jin et al., 1984 |
| | Fish | Fish mycobacteriosis (intestine/liver) | Contaminated water | Phung et al., 2013 |
| *M. chelonae* | Humans | Pseudo-outbreak | Contaminated endoscopy | Kressel & Kidd, 2001 |

| | | | | |
|---|---|---|---|---|
| | Humans | Cutaneous abscesses | Tap water contaminated instruments in liposuction | Meyers et al., 2002 |
| | Fish | Fish mycobacteriosis (intestine/liver) | Contaminated water | Phung et al., 2013 |
| *M. immunogenum* | Humans | Hypersensitivity pneumonitis | Metal removal fluids | Shelton et al., 1999 |
| *M. abscessus* | Humans | Sporotrichoid dermatosis | Public bath, Injections | Lee et al., 2000 ; Zhibang et al., 2002 |
| | Fish | Fish mycobacteriosis (intestine/liver) | Contaminated water | Phung et al., 2013 |
| *M. marinum* | Humans | Cutaneous infection | Aquarium management | Dorronsoro et al., 1997 |
| | Humans | Ulcerated nodule | Aquarium | Speight & Williams, 1997 |
| | Fish | Fish mycobacteriosis (intestine/liver) | Contaminated water | Phung et al., 2013 |
| *M. kansasii* | Humans | Cellulitis | Swimming at a beach | Hsu et al., 2002 |
| | | | Drinking water | Kaustova et al., 1981 |
| | | | Hospital water | Wright et al., 1985 |
| | | | Injections | Domergue et al., 2001 |
| | | | Industrial water | Chobot et al., 1997 |
| *M. ulcerans* | Humans | Ulcerative disease | Contaminated water bodies | Ross et al., 1997; Combe et al., 2017 |
| | Mammals | Ulcerative disease | Contaminated water bodies | Combe et al., 2017 |
| *M. szulgai* | Humans | Keratitis | Contamination from ice water | Holmes et al., 2002 |
| *M. palstre* | Humans | Cervical lymphadenitis | Contaminated water | Torkoo et al., 2002 |
| *M. gordonae* | Fish | Fish mycobacteriosis (intestine/liver) | Aquarium water | Phung et al., 2013; Primm et al., 2004 |

| | | | | |
|---|---|---|---|---|
| *M. scrofulaceum* | Humans | Lung lesions, lymphadenitis, granulomatous hepatitis, osteomyelitis, subcutaneous abscesses, renal infection | Hospital Tap water | Hsueh et al., 1996; Primm et al., 2004 |
| | Fish | Fish mycobacteriosis (intestine/liver) | Contaminated water | Toranzo et al., 2005 |
| *M. terrae* | Humans | Pulmonary and cutaneous infections | Fish tanks | Mycobacteria book |
| | | | Natural waters | Tuffley & Holbeche, 1980 |
| | | | Drinking water | Jin et al., 1984 |
| | | | Drinking water biofilm | Schulze-Robbecke et al., 1992 |
| | | | Hospital water | Lockwood et al., 1989 |
| | | | Recreational water | Dailloux et al., 1980 |
| | | | Injections | Zenone et al., 1999 |
| | | | Damp buildings | Huttunen et al., 2001 |
| *M. xenopi* | Humans | Pulmonary disease | Natural waters | Torkoo et al., 2000 |
| | | | Drinking water | Sniadack et al., 1993 |
| | | | Hospital water | Wright et al., 1985 |
| | | | Hospital equipment | Bennett et al., 1994 |
| | | | Hot water system | Wright et al., 1985 |
| | | | Recreational water | Slosareck et al., 1994 |
| *M. tusciae* | Humans | Cervical lymphadenitis | Drinking water (Tap water) | Tortoli et al., 1999 |
| *M. simiae* | Humans | Cervical lymphadenitis | No information | Cruz et al., 2007 |
| | Fish | Fish mycobacteriosis (intestine/liver) | Contaminated water | Toranzo et al., 2005 |
| *M. genavense* | Humans | Gastro-intestinal disorder | Drinking water, Hospital water | Ristola et al., 1999; Hillebrand-Haverkort et al., 1999 |

| | | | | |
|---|---|---|---|---|
| | Birds | Avian mycobacteriosis. Lesions in the liver and gastrointestinal tract | Water distribution systems | Portaels et al., 1996 |
| *M. intracellulare* | Humans | Pulmonary disease | Drinking water biofilm, Recreational water | Falkinham et al., 2001; Saito & Tsukamura, 1976 |
| | Birds, poultry | Tuberculosis-like disease | Contaminated water | Dhama et al., 2011 |
| | Pigs | Lymphadenitis | Contaminated water | Thorel et al., 2001 |
| *M. malmoense* | Fish | Fish mycobacteriosis (intestine/liver) | Contaminated water | Phung et al., 2013 |
| *M. gastri* | Fish | Fish mycobacteriosis (intestine/liver) | Contaminated water | Phung et al., 2013 |
| *M. segmatis* | Fish | Fish mycobacteriosis (intestine/liver) | Contaminated water | Phung et al., 2013 |
| | Humans | Skin or soft-tissue infections | Water (suspected only) | Wallace et al., 1988 |
| *M. bohemicum* | Fish | Fish mycobacteriosis (intestine/liver) | Contaminated water | Phung et al., 2013 |
| *M. phlei* | Fish | Fish mycobacteriosis (intestine/liver) | Contaminated water | Phung et al., 2013 |
| *M. poriferae* | Fish | Fish mycobacteriosis (intestine/liver) | Contaminated water | Toranzo et al., 2005 |
| | Humans | Pulmonary disease | Water (suspected only) | Ballester et al., 2011 |
| *M. triplex-like* | Fish | Fish mycobacteriosis (intestine/liver) | Contaminated water | Toranzo et al., 2005 |
| | Humans | Pulmonary disease | | Suomalainen et al., 2001 |
| *M. neonarum* | Fish | Fish mycobacteriosis (intestine/liver) | Contaminated water | Toranzo et al., 2005 |
| *M. trivale* | Fish | Fish mycobacteriosis (intestine/liver) | Water from aquaculture systems | Yanong et al., 2010 |
| *M. haemophilum* | Fish | Fish mycobacteriosis (intestine/liver) | Water from aquaculture systems, biofilm (tank surface water) | Yanong et al., 2010 |
| *M. gilvum* | Humans | Ulcerative disease | Contaminated water bodies | Combe et al., 2020 ; this study |
